# Blood biomarkers and breed genetics of aging in pet dogs

**DOI:** 10.64898/2026.03.07.710297

**Authors:** Vista Sohrab, Michelle E. White, Benjamin R. Harrison, Rob Bierman, Abbey Marye, Kathleen Morrill Pirovich, Diane P. Genereux, Kate Megquier, Xue Li, Brittney Kenney, Cindy Reichel, Dog Aging Project Consortium, Noah Snyder-Mackler, Joshua M. Akey, Daniel E. L. Promislow, Frances L. Chen, Elinor K. Karlsson

## Abstract

Pet dogs share human-like environments while aging on a compressed timescale, making them a powerful translational model for aging research. Using genomic and phenotypic data from 7,627 dogs in the Dog Aging Project, including 976 profiled for 159 blood metabolites and clinical analytes, we generated the first GWAS catalog in dogs. Blood traits map to orthologous loci in dogs and humans, indicating deeply conserved pathways. Breed ancestry explains substantial variance in blood traits, and selection on visible characteristics such as fur type has pleiotropic metabolic effects. Leveraging mosaic ancestry in mixed-breed dogs and longitudinal mortality data, we identify blood traits elevated in short-lived breeds that predict individual mortality risk — including globulin and potassium — and protective traits enriched in long-lived breeds, such as ethanolamine. Although some aging-associated traits relate to growth hormone pathways, many do not, indicating that aging in dogs is multifactorial. These findings establish dogs as a translational system for identifying genetic determinants and biomarkers of aging relevant to extending healthy lifespans.

## Main Text

Aging is characterized by progressive declines in physiological and cognitive function, widespread metabolic remodeling, and an increased susceptibility to chronic disease, placing a substantial burden on individuals, families, and healthcare systems(*1–5*). The rate and trajectory of these changes vary widely among individuals, reflecting a complex interplay of genetic, environmental, and lifestyle factors(*6*, *7*). Large-scale genetic studies, including genome-wide association studies (GWAS), have identified thousands of loci associated with exceptional longevity and age-related diseases (*8–12*). However, the biological mechanisms linking these loci to functional decline remain poorly understood, limiting their translation into effective therapeutic strategies.

Genetic studies of blood traits offer a complementary and potentially more mechanistic approach to understanding aging than studies of longevity alone. Many circulating blood traits are linked to aging-related pathways. Clinical analytes such as serum chemistry profiles and complete blood counts are shaped by both genetic variation (*13–16*) and age-related physiological change(*17–20*) and serve as biomarkers of disease risk that may reflect processes contributing to pathology (*21–23*). The plasma metabolome, which comprises thousands of small molecules that reflect metabolic activity, predicts mortality, frailty, and functional decline in both dogs and humans (*24–28*). Identifying genetic variation underlying individual differences in blood trait levels and longitudinal trajectories can yield new interventions for aging-related diseases, as exemplified by LDL cholesterol and the development of statins for cardiovascular disease (*29*).

In humans, identifying modifiable metabolic or environmental factors that influence aging is constrained by the long human lifespan(*30*). Pet dogs (also known as companion dogs) share many features of human aging: they exhibit heterogeneous aging trajectories shaped by genetic, environmental, and lifestyle factors (*31–33*), yet age on an accelerated timescale, with lifespans of roughly 9–15 years (*34–36*). As household animals, dogs experience many of the same dietary, social, and environmental exposures as their owners and often receive comparable medical care(*37*, *38*). This close human–dog relationship enables collection of high-resolution longitudinal data, including veterinary clinical records, plasma metabolomic profiling, and owner-reported measures of physical and cognitive function (*39–43*). Aging-associated measures such as plasma metabolite mortality hazard ratios are strongly correlated between dogs and humans, supporting the translational relevance of this system(*44*).

Blood traits in dogs, as in humans, vary with age, sex, and size, but in dogs they also vary with breed (*43*, *45–47*). Modern dog breeds are inbred populations created within the last 175 years through severe population bottlenecks and ongoing selective breeding for physical and physiological traits (*48–51*). In the United States, most dogs are either single-breed (“purebred”) or mixed-breed animals with ancestry from multiple breeds(*52*). This unusual population history alters the allele frequency spectrum, enabling well-powered GWAS in smaller cohorts than are typically required in humans(*53*, *54*).

Single-breed dogs tend to have shorter lifespans than mixed-breed dogs(*34*), and some large-bodied breeds are particularly short-lived. Variation in adult body size among breeds is strongly associated with genes in the growth hormone/insulin-like growth factor 1 (GH/IGF-1) pathway (*55–57*). In rodent models, manipulation of this pathway profoundly affects aging (*58–61*), leading to speculation that selection on GH/IGF-1 variants contributes to the shortened lifespan of some dog breeds(*62*). However, body size alone does not account for the full variation in lifespan across breeds (*63*, *64*). Most mixed-breed dogs in the United States have ancestry from four or more breeds, often spanning a wide range of body sizes(*52*), creating a natural experiment in which the effects of size can be separated from other breed-associated genetic influences on aging.

Here, we leverage the unique combination of breed ancestry, environmental diversity, and deep longitudinal phenotyping in dogs from the Dog Aging Project to identify blood traits associated with variation in lifespan and to evaluate their relevance for human aging.

## Dog Aging Project

The Dog Aging Project is the largest study of the biological, genetic, and environmental factors that influence aging in dogs(*65*). As of December 2025, the project has enrolled 52,705 dogs. Here, we analyze the subset of 7,627 dogs with genetic data from the 47,444 dogs enrolled through the end of 2023. DNA samples were collected by owners using saliva swabs, sequenced (mean depth of 0.79x ± 0.41x; range 0.14x to 5.36x), and aligned to the CanFam4.0 reference genome(*66*). Genotypes were imputed using a panel of 1,929 deeply sequenced (12.95x to 31.06x) dogs and related canids (*67*). Our final dataset includes 29,089,701 biallelic SNPs, of which 9,871,985 are common (minor allele frequency ≥ 0.01). In dogs, we previously showed that low-pass sequencing with imputation achieves array-comparable genotype accuracy while generating marker densities 10- to 50-fold higher, providing sufficient resolution to capture most common variation(*52*).

### Genetically determined breed ancestry

The sequenced dogs ranged in age from a few weeks to 21 years at enrollment (mean ± SD, 4.6 ± 3.5 years) and are younger than the full cohort of enrolled dogs (6.8 ± 4.3 years)(Fig. 1A,B; fig. S1A). Approximately half are female (48.8%), and 82% are sterilized, similar to the U.S. pet dog population (fig. S1B)(*68*). We estimated breed ancestry using supervised admixture analysis against a reference panel comprising 109 common breeds selected from the 360 breeds recognized worldwide, together with village dogs and wolves(*69*). We classified dogs as single-breed or mixed using owner-reported information, which we have previously shown to be reliable(*52*, *69*) and which allows our dataset to include breeds not represented in the reference panel.

**Fig. 1.**
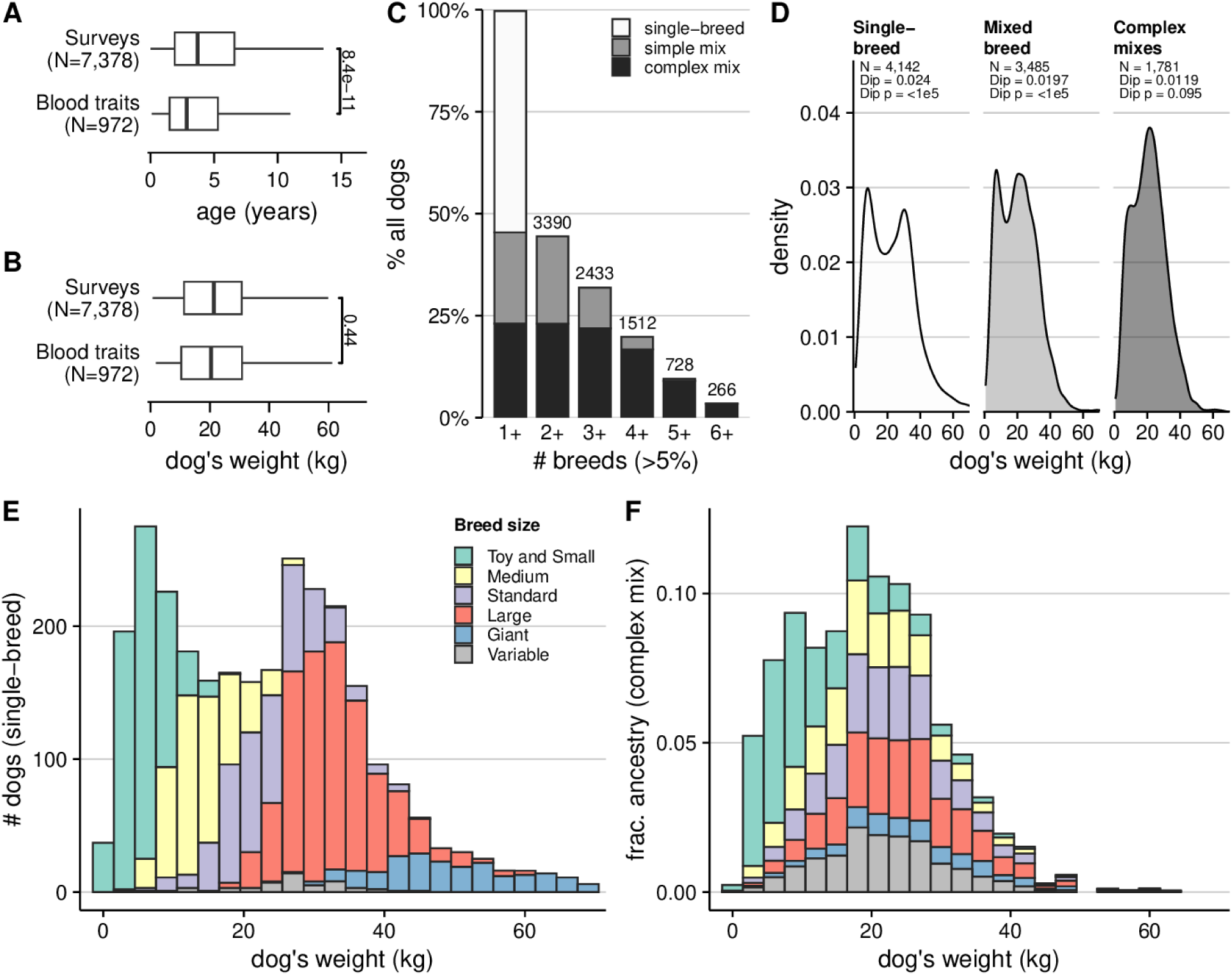
Demographic and genetic structure of the study population. **(A)** Dogs with blood trait data are somewhat younger than the full enrolled population and **(B)** do not differ in body weight. (**C)** Nearly half of dogs (45%) have detectable ancestry (>5%) from more than one breed, including simple mixes (dogs with >40% ancestry from any single breed) and more complex mixes. **(D)** Body weight distributions stratified by ancestry complexity show bimodality in single-breed dogs and in all mixed-breed dogs, but not among complex mixes, using Hartigan’s dip test for unimodality (100,000 bootstrap replicates). **(E)** In single-breed dogs, body weight is correlated with size specified in the breed conformation standard. **(F)** In complex mixed-breed dogs, the relationship between body size and breed ancestry is less structured, with large-breed ancestry present in small dogs and vice versa, and few dogs of exceptionally small or large size.

Mixed-breed dogs have broadly similar breed ancestry proportions to those observed among single-breed dogs. Overall, 45.7% of sequenced dogs are mixed-breed, and half of mixed-breed dogs (51%) are “complex mixes” with no more than 40% ancestry from any breed (Fig. 1C). Breeds that are common among single-breed dogs also comprised a larger proportion of mixed-breed ancestry (Spearman’s ρ = 0.70). Amongst the most common breeds, ancestry from golden retrievers and labrador retrievers is more common in single-breed dogs, and ancestry from American pit pull terriers, poodles, toy poodles, and Chihuahuas is more common in mixed-breed dogs (fig. S2).

Breed-related population structure influences the distribution of adult body weight, producing deviations from the approximately normal distribution expected in a randomly mating population (*70*). In single-breed dogs, the popularity of very small and very large breeds produces a bimodal weight distribution (n = 4,163; Hartigan’s dip test, p < 1 × 10⁻⁵) (Fig. 1D,E). In contrast, the weight distribution in complex-mix dogs does not reject unimodality (n = 1,760; dip p = 0.089), and dogs with mixed ancestry rarely have extreme body sizes (Fig. 1D,F). Breed ancestry explained less variance in body weight in mixed-breed dogs (R^2^ = 0.75) than in single-breed dogs (R^2^= 0.84), and even less in complex mixes (R^2^ = 0.68); all associations were significant (permutation p≤1x10^-5^; table S1). This pattern supports our conceptual framework: because the tight coupling of body size and genetic ancestry in single-breed dogs(*71*) is disrupted in mixed-breed dogs, they provide an opportunity to disentangle the effects of size from other breed-associated factors that influence aging.

### Phenotypes

All owners completed baseline surveys (959 questions) about their dog’s environment, lifestyle, health, and behavior at enrollment. We selected 154 questions from the baseline survey for genetic analysis: all questions with standardized response types, meaningful variation between individuals, a sufficient number of responses, and potential to be heritable (data S1; fig. S3). This included 26 questions on physical activity, 40 questions on behavioral and cognitive health, and 29 questions on environment, medical history, diet and weight (*65*, *72*). We also included two surveys added a year into the project that consequently have fewer responses: the Dog Obesity Risk and Appetite (DORA) survey (34 questions and 4,895 dogs)(*73*) and the Monash Dog Owner Relationship Scale (MDORS; 25 questions and 4,915 dogs)(*74*)(table S2).

We collected blood-based molecular phenotypes (blood traits) for a subset of dogs (fig. S1). Dogs with blood trait data are younger (3.8 ± 3.1SD years) than the full sequenced cohort (4.6 ± 3.5 SD years) but are otherwise similar (Fig. 1AB). After excluding traits whose distributions were dominated by rare, extreme values and therefore poorly suited for GWAS, we analyzed 159 blood traits: 36 clinical analytes measured in up to 971 dogs (complete blood counts [CBC] and serum chemistry profiles [SCP]) and 123 plasma metabolites measured in 937 dogs (table S2).

### Blood traits are more heritable than survey traits

Blood traits show higher SNP-based heritability (h²_SNP) than survey-derived traits (median h²_SNP = 0.39 vs. 0.15; permutation p < 1 × 10⁻5) (Fig. 2A). We estimated h^2^_SNP_using restricted maximum likelihood with correction for linkage disequilibrium(*75*), adjusting for age, body weight, sex (sterilization status is modeled as part of sex), and – for blood traits – hours fasted. Consistent with the high heritability of height in pet dogs (*52*), body weight is highly heritable(h^2^_SNP_ = 0.81 ± 0.02; N=7378). Two blood traits show even higher heritability: serum creatinine (h^2^_SNP_ = 0.88 ± 0.13; N=953) and cystathionine (h^2^_SNP_ = 0.87 ± 0.12; N=924)(Fig. 2A)(*76*, *77*). In total, 29 blood traits have heritability estimates exceeding 50%, including 9 clinical analytes and 20 plasma metabolites (fig. S4).

**Fig. 2.**
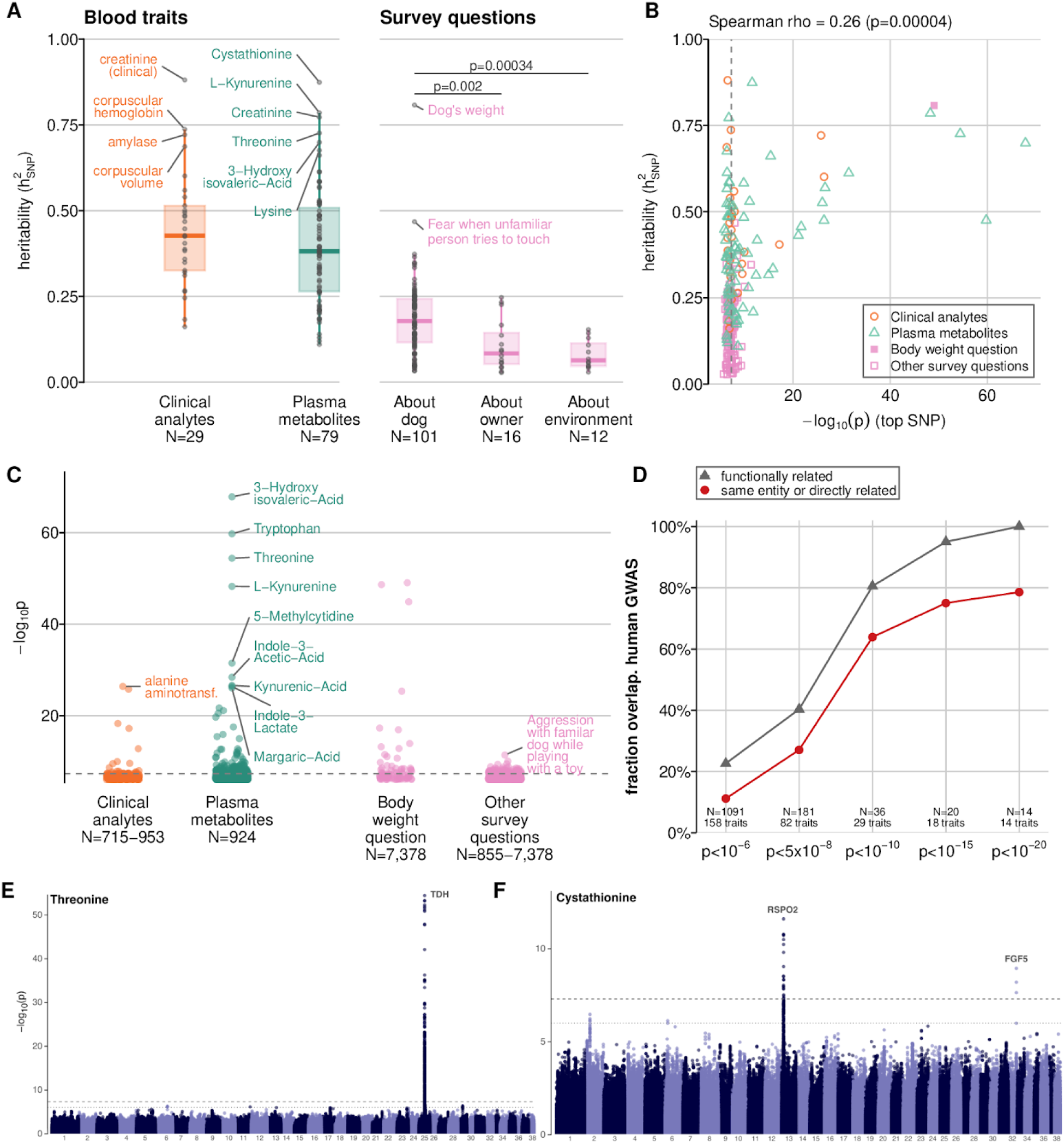
Heritability and GWAS of blood traits and survey questions in dogs. **(A)** SNP-heritability for blood traits and for survey questions shows that questions about the dog are more heritable than those about the owner’s relationship with the dog (“owner”; (74)) or the dog’s environment. Brackets indicate two-sided permutation tests (100,000 permutations) comparing median heritability. Boxplots show median, interquartile range, and whiskers (1.5× IQR), with the most heritable traits labeled. **(B)** Traits with higher SNP heritability exhibit stronger top genome-wide associations (Spearman ρ; two-sided permutation test, 100,000 permutations). **(C)** Top associations for blood traits and survey questions. **(D)** Fraction of dog GWAS regions exceeding the indicated significance threshold that overlap a genome-wide significant human GWAS signal for the same or a functionally related trait. N indicates the number of dog GWAS regions above the threshold. **(E)** GWAS for threonine levels (924 dogs) and **(F)** cystathionine levels (924 dogs).

With the exception of body weight, none of the survey-derived traits exceed 50% heritability (Fig. 2A). This pattern likely reflects a focus on traits that are strongly influenced by non-genetic factors. The most heritable survey traits concern eating behaviors, size-influenced behaviors, and behaviors affecting ease of ownership, such as fearfulness and biddability (fig. S4, data S2). In contrast, survey questions related to the dog’s environment show lower heritability, and those from a survey focused on the owner–dog relationship show the lowest heritability.

## Building the first GWAS catalog for dogs

To systematically explore the genetic architecture of canine traits, we conducted GWAS for each available phenotype and generated the first comprehensive GWAS catalog for dogs (<u>broad.io/dogPheWeb</u>) using a dog-adapted implementation of the PheWeb software(*78*). This interactive platform currently provides variant–phenotype associations for 154 survey-based traits and 159 blood-based traits (fig. S5).

To mitigate the effects of complex population structure, we performed GWAS using the leave-one-chromosome-out mixed linear model (MLMA-LOCO) implemented in GCTA, incorporating the same covariates used in our heritability analyses(*52*, *79*). Given the diverse breed ancestries in our cohort and the correspondingly short linkage disequilibrium (*50*, *52*, *80*), we adopted standard human GWAS thresholds (p < 5.0×10⁻⁸ for genome-wide significance; p < 1.0×10⁻⁶ for suggestive associations) (*81*, *82*)(data S3).

In total, we identified 298 genome-wide significant loci with a mean size of 283 kb (SD = 378 kb), defined by clumping nearby SNPs (<250 kb) and in linkage disequilibrium (r² > 0.2)(*83*, *84*). Traits with higher heritability are more likely to have at least one genome-wide significant locus (logistic regression OR = 9.5, p = 0.0036), and have a greater number of significant loci overall (quasi-Poisson β = 3.1, p = 2.6×10^-8^) (Fig. 2B). Body weight illustrates this pattern clearly, yielding 33 distinct associations (Fig. 2C), including replicating published associations at the genes IGF1 (p=9.0x10^-50^), HMGA2 (p=2.35x10^-49^) and LCORL (1.33x10^-45^) (*52*, *55*, *56*, *77*, *85*). We also identified a previously unreported association near microRNA-497 (p = 1.3x10^-14^), a regulator of IGF1R adjacent to an insulin-signaling gene (*86*, *87*).

The remaining 153 survey questions have 76 significant associations, with 67% having no significant hits, 22% having one significant hit, and 11% having two or more. The significance of associations tends to be lower for survey-based traits (excluding body weight) than for blood traits (Fig. 2C). The top association is for the survey question “Aggression with a familiar dog while playing with a toy” (p_GWAS_ = 4.1x10^-12^), and maps to a locus that in humans is associated with circulating sex hormone–binding globulin levels (rs79391862-A, p = 2x10^-121^(*88*)), a phenotype that primarily reflects metabolic and endocrine status but may influence behavior indirectly.

In contrast, blood-based traits yield more, and more significant, associations than survey traits (Fig. 2C). Among the 159 blood traits, we identified 189 genome-wide significant associations (1.19 ± 2.35 per trait), with 82 blood traits (52%) having at least one association. The strongest associations are for plasma metabolites, and map to genes with well-established biochemical connections. For example, we replicate a previous association between the gene GPT and circulating alanine aminotransferase levels(*54*). The most significant association is between the metabolite 3-hydroxyisovaleric acid and a SNP in the gene MCCC2, which encodes an enzyme that prevents accumulation of this metabolite.

### Locus heterogeneity and pleiotropy in blood traits

The genetic architecture of blood traits in dogs shows extensive locus heterogeneity and pleiotropy, mirroring patterns observed in human metabolomic and hematologic GWAS(*29*, *89*, *90*). Among the 82 traits with at least one significant association, 37 map to multiple independent loci, led by the metabolites 3-hydroxyisovaleric acid (17 loci), 5-methylcytidine (14 loci), and S-methylcysteine (13 loci). Thirteen of the 189 genomewide significant loci associate with more than one blood trait.

The most pleiotropic locus was a 95-kb region on chromosome 13 containing ALB, which encodes albumin. Variants at this locus were associated with tryptophan levels (p = 1.8x10^-60^), albumin/globulin ratio (p = 1.1x10^-17^), albumin levels (p = 1.9x10^-13^) and globulin levels (p = 2.7x10^-8^). This pattern is biologically expected: ∼90% of circulating tryptophan is albumin-bound(*91*, *92*), so genetic variants that alter albumin concentration or binding capacity simultaneously affect tryptophan availability. Consistent with this mechanism, the ALB locus was also associated with metabolites known to bind albumin (fig. S6), including L-kynurenine (p = 5.6x10^-49^), indole-3-acetic acid(p = 4.1x10^-29^), indole-3-lactate(p = 4.2x10^-27^), and indole-3-propionate (p = 2.3x10^-13^)(*93*, *94*).

### Overlap of blood trait loci in dogs and humans

The loci most significantly associated with blood traits in dogs are orthologous to loci implicated in human studies, pointing to shared biological mechanisms and underscoring the value of dogs as a translational model. We compared the dog associations with those reported at orthologous human GWAS loci (±250 kb) across 25 studies (*95–120*) (data S4). Amongst all 181 dog loci, 27% overlap a human association for the same or a directly related trait. This overlap rises sharply when restricting to the most significant dog signals: among the 14 loci with p < 1x10^-20^ in dogs, 79% are associated with the same or a directly related trait in humans (Fig. 2D).

Despite these broad similarities, several metabolic pathways show clear species-specific differences. In dogs, one of the strongest GWAS signals is the association between threonine levels and TDH, the gene encoding L-threonine dehydrogenase (p = 3.9x10^-55^) (Fig. 2E). In humans, TDH is a non-functional pseudogene that is not associated with threonine or any related traits(*120*, *121*). Threonine levels in humans are instead associated with loci such as the gene GCKR, which modulates hepatic metabolic flux (*104*, *122*). Additionally, dogs lack the strong association between tryptophan levels and the gene TDO2 observed in humans. TDO2 encodes the rate-limiting enzyme of the kynurenine pathway and is a primary regulator of systemic tryptophan turnover (*123*). In dogs, the top signal instead maps to the gene encoding albumin (ALB; p = 1.8x10^-59^), which can influence tryptophan levels by altering the bound versus free fraction.

## Breed shapes blood traits in single-breed dogs

Breed ancestry is the dominant determinant of variation in both plasma metabolites and clinical analytes in dogs. To assess the amount of variation explained by breed compared to other individual traits like age, weight and sex, we compared dogs with single-breed ancestry (N=357-475 dogs in each of 81-87 breeds for each blood trait), using an analysis of variance (type II ANOVA)(data S5). Breed explained on average 25.1 ± 5.1% of the variance, larger than for age (1.9 ± 2.5%), sex (1.8 ± 1.8%), or weight (0.5 ± 0.7%) (Fig. 3A). As expected, more heritable traits have more variance explained by breed (slope = 0.22, *p* < 0.0001) but not by age, sex, or weight (slopes = –0.003, 0.016, and 0.001; *p* = 0.83, 0.22, and 0.95, respectively)(Fig. 3B).

**Fig. 3.**
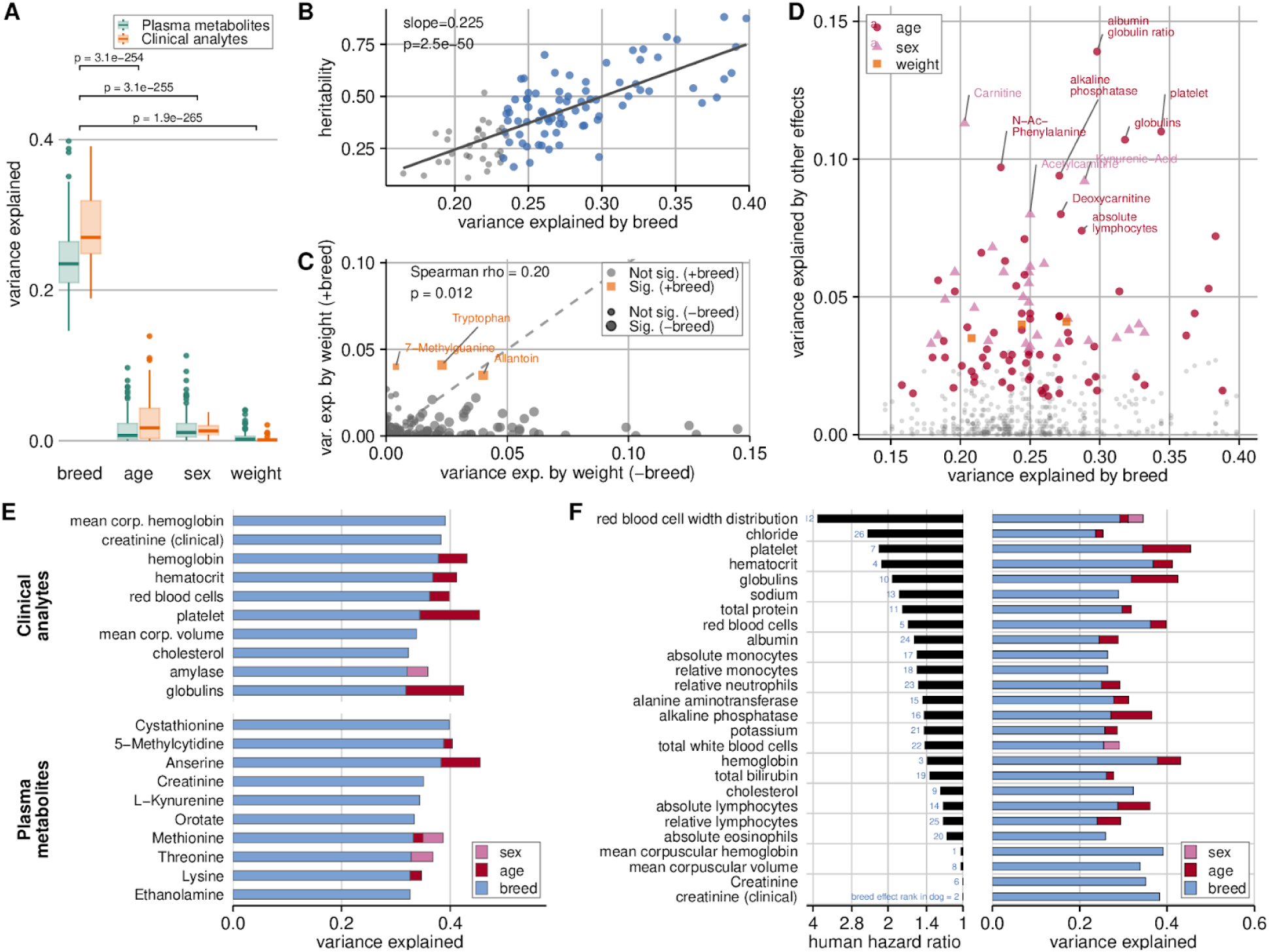
Breed explains substantial variation in blood traits in single-breed dogs. **(A)** Breed explains more variance than age, sex, or body weight (generalized η², type II ANOVA). Differences were tested using a mixed-effects model with phenotype as a random effect; p values are from post hoc contrasts comparing breed with each effect. **(B)** Variance explained by breed increases with trait heritability (GREML–LDS constrained). The line shows a linear fit; slope and p value are from a mixed-effects model testing the heritability–effect interaction. **(C)** Including breed reduces variance attributed to body weight for some traits, consistent with shared breed–weight structure. Weight variance is shown for models with (+breed) and without (−breed) breed; traits with FDR-adjusted p < 0.05 are highlighted. **(D)** Age, sex, and weight generally explain less variance than breed. Grey points indicate non-significant effects (pFDR > 0.05) (pFDR > 0.05); the 10 strongest effects are labeled. **(E)** Variance partitioning across age, breed, sex, and weight for the 20 traits most influenced by breed, shown as stacked bars; none show a significant weight effect. **(F)** Comparison of dog and human effects for clinical analytes. Left, published human hazard ratios; right, variance explained by breed in dogs.

Body weight explains only a small fraction of the variance in any blood trait, in part due to the strong correlation between breed and size (Fig. 3A)(*71*). Multicollinearity diagnostics show that this correlation does not generally obscure independent weight effects in our models (variance inflation factor, VIF = 2.72). For a subset of blood traits, however, inclusion of breed markedly attenuates the apparent effect of body weight. For example, in models excluding breed, weight explains 14.5% of the variance in creatinine levels, a metabolite reflecting total muscle mass (*124*); when breed is included, weight explains nearly none of the variance (0.4%), and breed explains 38.3%. Across the 14 blood traits for which weight explains more than 5% of the variance in models excluding breed, inclusion of breed reduces the weight effect by an average of 94.1% ± 6.7%, leaving only 0.01%–1.4% of the variance explained (Fig. 3C).

Overall, a significant (p_FDR_<0.05) amount of variation is explained by breed for 96 blood traits, more than for age (N=68), sex (N=28) or weight (N=3)(Fig. 3D). For example, age explains substantial variation in albumin-globulin ratio (13.9%; p_FDR_ = 2.7x10^-11^) and platelets (11.0%, p_FDR_ = 8.5x10^-7^), which are both linked to aging in humans (*125–127*). Sex explains variation in carnitine levels (11.3%; p = 7.5x10^-8^), which is linked to sexual dimorphism in humans(*128*), and in kynurenic acid (effect size = 9.2%; p =3.1x10^-6^), which is a tryptophan metabolite modulated by testosterone and estrogen (*129*).

### Selection for fur type in breeds influences longevity-related blood trait

The influence of breed on blood traits in dogs is likely driven in part by historic selection on aesthetic and functional characteristics. The strong effect of breed on creatinine levels is consistent with selection on size and muscle mass (*130–132*). Similarly, breed explains 32.1% of variance in amylase levels (p_FDR_ = 2.0x10^-5^), a trait previously shown to vary between breeds due to copy-number variation at the AMY2B gene related to adaptation to starch-rich diets (*133*, *134*).

For at least one metabolite, a large effect of breed may be the pleiotropic consequences of selection on a seemingly unrelated aesthetic trait. The two loci associated with cystathionine levels, which is the second-most breed influenced blood trait (40% of variance explained,p_FDR_ = 2.1x10^-10^), are both in genes that determine fur characteristics: the hair growth pattern gene RSPO2 (p = 2.6 × 10^-12^) and the hair length gene FGF5 (p = 1.2x10^-9^)(*135*) (Fig. 2F, Fig. 4A,B). This differs from humans, where cystathionine levels are associated with variation in the gene CTH, which encodes cystathionine γ-lyase(*104*). In a subset of 50 mixed and single-breed dogs with both metabolite measurements and photographs, dogs with the RSPO2-linked furnishings trait (beards and bushy eyebrows) and dogs with longer fur had significantly higher cystathionine levels (Fig. 4C; table S3). Furthermore, the variants most associated with cystathionine levels were also associated with the corresponding fur trait (fig. S7; data S6). Finally, the cystathionine associated SNPs are differentiated between breeds according to their fur characteristics (Fig. 4D; data S7), consistent with evidence of selection at both loci (*136*).

**Fig. 4.**
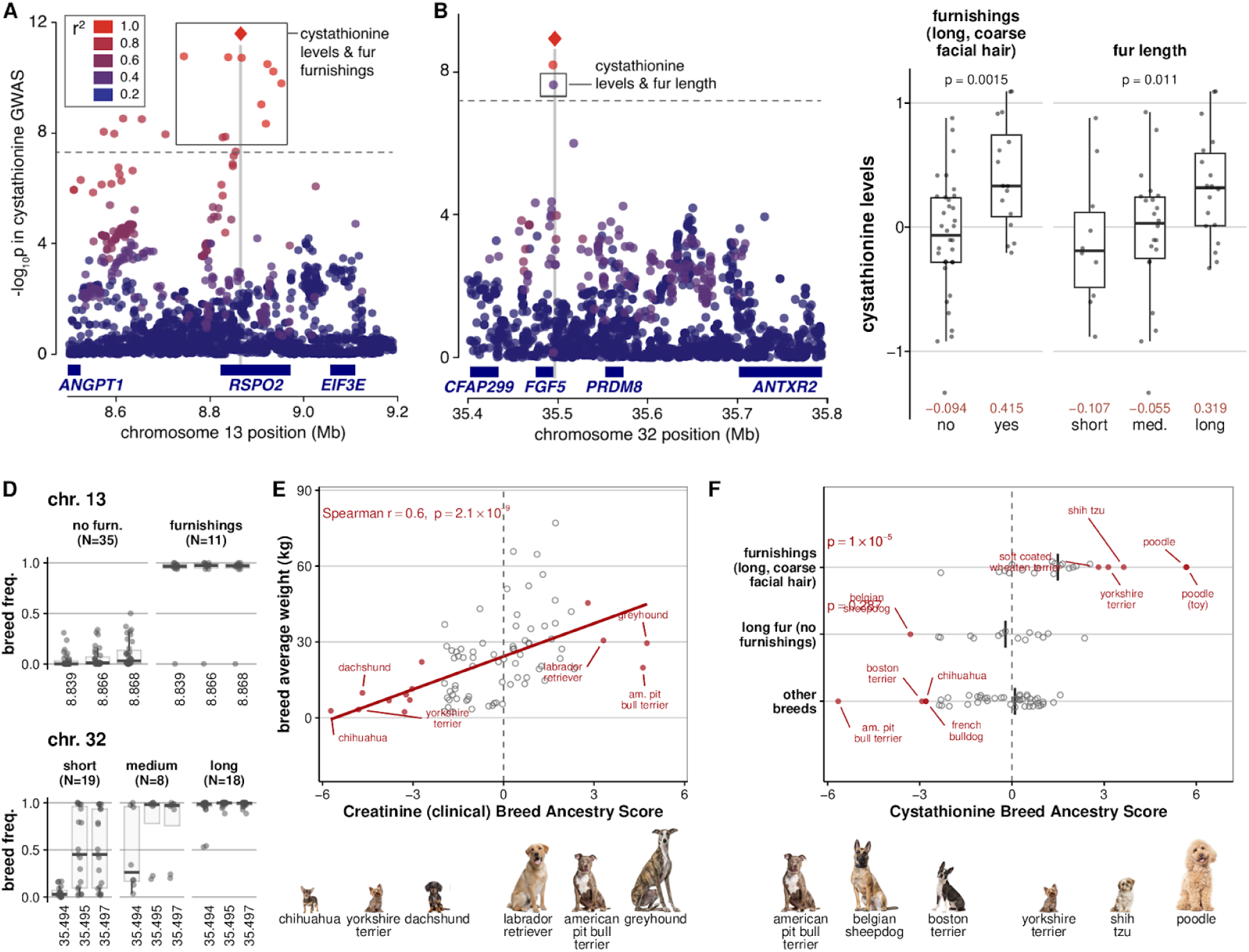
Ancestry composition links variation in blood traits to breed differences. **(A,B)** Regional association plots show two cystathionine loci colocalizing with canonical fur loci for furnishings (chr13) and fur length (chr32). Boxes denote SNPs significantly associated with both cystathionine levels and the indicated fur trait in 50 dogs phenotyped for both. **(C)** Normalized cystathionine levels by fur phenotype in 50 dogs. Points represent individuals; boxes show medians and interquartile ranges. P values are from a Wilcoxon rank-sum test (furnishings) and Kendall’s rank correlation (fur length); group means are shown in red. **(D)** Breed-level minor allele frequencies for the top cystathionine-associated variants stratified by fur phenotype; boxes summarize breed distributions within each class. **(E)** Breed-ancestry effects on creatinine estimated using linear mixed-effects regression. The ancestry score is the REML t statistic; the dashed line indicates no breed effect (t = 0). Points represent breeds (open gray, not significant; filled red, pFDR < 0.05). The line shows the relationship between breed effect size and mean breed body weight. **(F)** Breed-ancestry effects on cystathionine are stronger in breeds with furnishings and long fur. Breeds with the largest effects are labeled; text reports the two-sided t-test p value.

### Aging-related blood traits are not disproportionately influenced by breed

In analyses focused only on single-breed dogs, the blood traits most strongly influenced by breed were not disproportionately related to aging or longevity. We defined aging-related blood traits as the 32 traits used in two validated blood-based aging clocks in humans (Aging.AI(*22*) and PhenoAge(*137*); table S4). These traits were not more strongly influenced by breed than other blood traits, based on a linear model comparing gene-effect scores while controlling for phenotype category (clinical analyte versus plasma metabolite) (β = 0.019, p_FDR_ = 0.40). Within this set of aging-clock traits, the variance explained by breed was not higher for traits conferring greater mortality hazard ratios in humans (Spearman’s ρ = −0.08, p_FDR_ = 0.91).

## Breed ancestry proportions disentangle breed effects

To identify blood traits that underlie lifespan differences between dog breeds, we leveraged the genetic diversity of the pet dog population by incorporating mixed-breed dogs. This increased sample size and statistical power and enabled us to disentangle breed effects from body size. For each blood trait, we estimated how ancestry from individual breeds predicts deviation from the population mean using linear mixed-effects models, yielding a Breed Ancestry Score that quantifies the effect of breed ancestry on trait levels(*52*). We then tested whether variation in these Breed Ancestry Scores was explained by breed lifespan (data S8).

We validated this approach using blood traits known to correlate with phenotypic differences between breeds. For example, breed weight explains 25.4% of the variance in breed ancestry scores for creatinine, a biomarker of muscle mass (*124*), with ancestry from heavier breeds predicting higher creatinine levels (n=82; Fig. 4E). Similarly, ancestry from breeds with furnishings and long fur predicts higher cystathionine levels, consistent with pleiotropy at the two associated loci (Fig. 4F).

## Blood trait variation and breed lifespan

Overall, variation in Breed Ancestry Scores for 19 blood traits is significantly explained (p_FDR_ < 0.05) by either body weight (17 traits) or lifespan (3 traits, including one also associated with weight) in a model including both variables (Fig. 5A, fig. S8, data S9). In contrast to single-breed analyses, the effect of body weight is no longer masked by its strong correlation with breed. Ancestry from longer-lived breeds was associated with lower potassium levels, higher sodium–potassium ratios, and higher chloride levels (β = –0.51 to 0.54; pFDR = 0.03 for all; Fig. 5B).

**Fig. 5.**
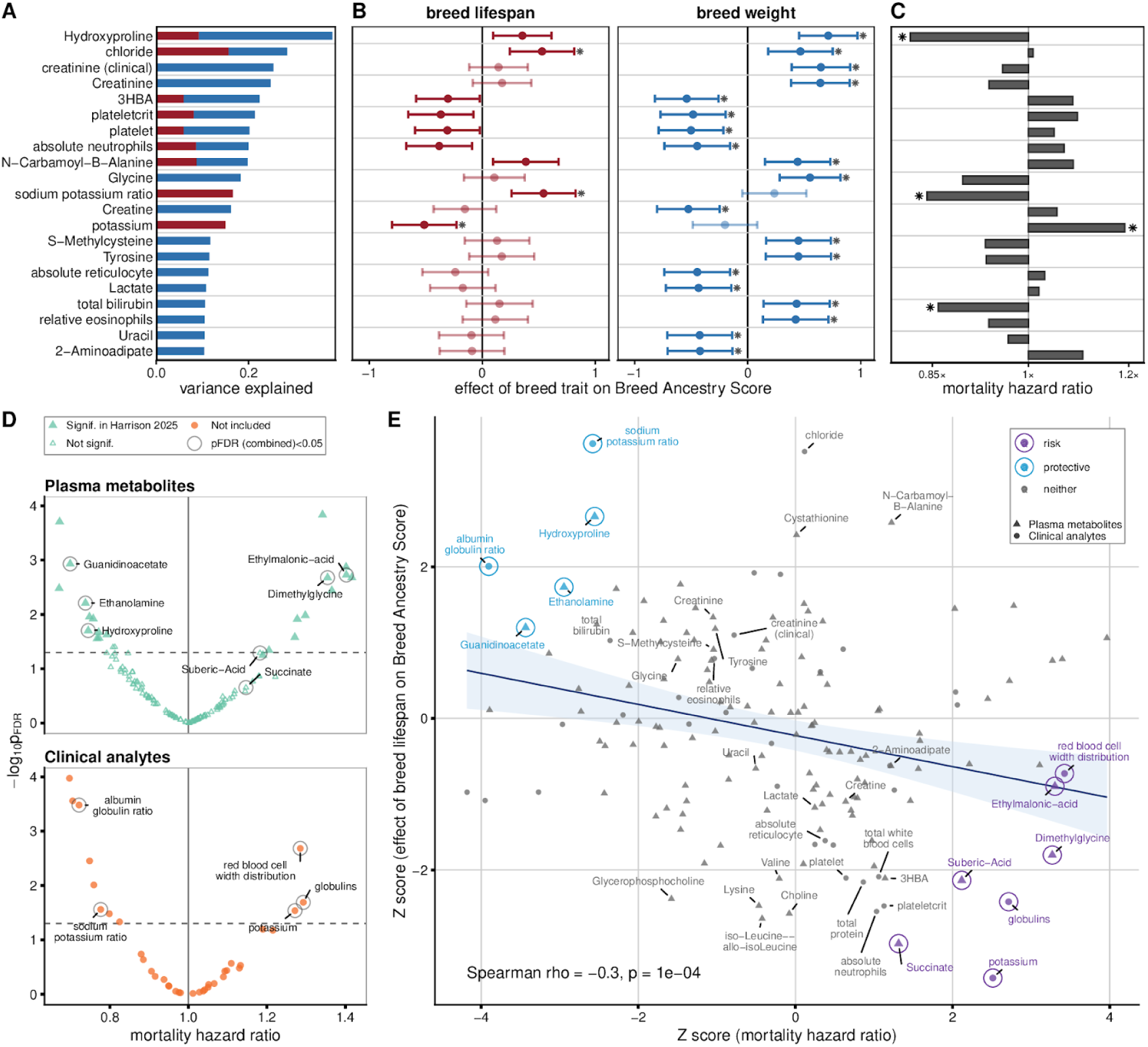
Breed Ancestry Scores distinguish blood traits predictive of mortality. **(A)** Blood traits with significant breed ancestry score variation (pFDR < 0.05) explained by breed lifespan (red) and/or breed body weight (blue). Stacked bars show the variance explained by each covariate, expressed as generalized η² from linear mixed-effects models (Breed Ancestry Score ∼ breed weight + lifespan). **(B)** Effect estimates for the traits in (A), shown as β ± 1.96 s.e. from the same models and faceted by covariate; asterisks indicate pFDR ≤ 0.05. **(C)** Mortality hazard ratios for individual dogs; asterisks denote pFDR ≤ 0.05. **(D)** Mortality risk in individual dogs faceted by trait type (circled and labeled points; combined pFDR ≤ 0.05). **(E)** Concordance between individual-level mortality hazard ratios and breed-level effects. Points represent breeds (circled, pFDR ≤ 0.05; labeled, included in **(A)**), with a linear fit (blue line) and 95% confidence interval (shaded).

### Correlation of breed lifespan effect with mortality hazard

We estimated all-cause mortality hazard ratios for all 159 blood traits—including plasma metabolites and clinical analytes—in 957 dogs with three years of longitudinal follow-up, during which 11% died. Using an approach we previously developed (*44*), we modeled mortality risk as a function of each trait while adjusting for age at enrollment, body weight, and sex (data S10). Thirty-four traits were significantly associated with mortality risk (p_FDR_ < 0.05)(fig. S9), including several traits whose levels vary with breed ancestry and lifespan (Fig. 5C,D).

Breed-lifespan effects and mortality hazard ratios are more strongly correlated (Spearman rho = –0.31; p = 5.5x10^-5^) (Fig. 5E) than breed-weight effects (Spearman rho = -0.2, p= 0.013). In other words, traits for which ancestry from short-lived breeds predicts higher values also tend to confer higher mortality risk in individual dogs (“risk” traits), and vice versa for “protective” traits. To confirm that this relationship did not reflect the breed ancestry of the dogs in the Cox analysis, for a subset of five significant blood traits (p_FDR_<0.05), we assessed whether breed ancestry explained mortality risk by adjusting for ancestry from the nine most influential breeds. All five blood traits remained significant even after adjusting for ancestry (fig. S10; table S5).

### Growth-hormone related metabolites are not exceptional

Plasma metabolites known to be affected by growth hormone/insulin-like growth factor 1 (GH/IGF-1) signaling are not more strongly implicated in lifespan than other metabolites. Of the 123 plasma metabolites analyzed, 24 have been shown to change in response to perturbation of the GH/IGF-1 pathway in rodent models(*138–142*)(table S6). GH–related metabolites had lower absolute mortality hazard ratios (mean |z| = 0.96 vs. 1.35; Wilcoxon p=0.0092). Breed ancestry scores for GH-linked metabolites and other metabolites are similarly correlated with breed lifespan (fig. S11).

### Identifying risk and protective blood traits

To identify traits associated with lifespan at both breed and individual levels, we combined the z-scores from the breed-level and individual-level analyses (data S11), yielding 13 traits with p_FDR_<0.05 (six protective, seven risk-associated). For example, both globulin and potassium were risk-associated: ancestry from short-lived breeds predicted higher levels, and elevated levels in individual dogs were linked to increased mortality (Fig. 6). This mirrors human aging, where elevated globulin (HR > 1.9) and elevated potassium (HR > 1.4) increase mortality risk (*143*, *144*).

**Fig. 6.**
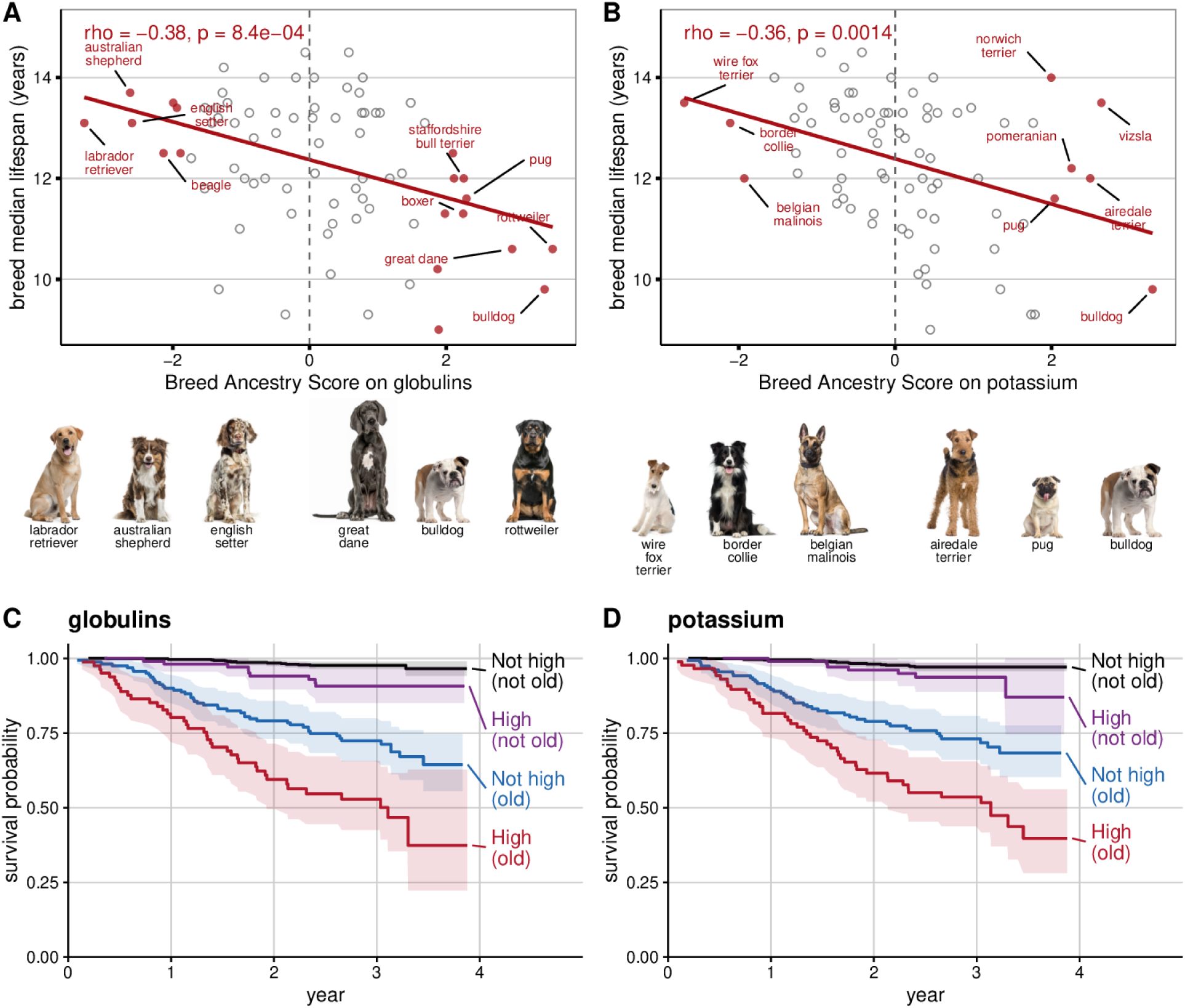
Two blood traits higher in short-lived breeds are associated with lower probability of survival. **(A)** Breed Ancestry Scores (REML t) are plotted against breed lifespan for globulin levels and **(B)** potassium levels. Points are breeds (red, p_FDR_≤0.05; gray, p_FDR_>0.05, top breeds labeled) with linear fit (red line) and Spearman’s correlation (ρ). Vertical dashed gray line denotes no ancestry effect. Images depict representative breeds with the strongest effects. **(C–D)** Kaplan–Meier survival curves for dogs stratified by age at enrollment (old, ≥7 years; not old, <7 years) and by normalized abundance of globulins or potassium (high, top quartile) as a function of years since enrollment (*44*).

Eight plasma metabolites—four risk-associated and four protective—were implicated in the combined analyses (Fig. 5E, fig. S12). Risk-associated metabolites included suberic acid, a dicarboxylic fatty acid indicative of mitochondrial stress that accumulates with age in humans (*145*); dimethylglycine, a marker of renal dysfunction (*146*) associated with increased all-cause mortality(*147*, *148*); and ethylmalonic acid and succinate, both markers of mitochondrial dysfunction. Protective metabolites included ethanolamine, which extends lifespan in yeast, fruit flies, and mammalian cell lines by enhancing autophagy(*149*); deoxycarnitine, a biomarker of carnitine pathway flux, reductions of which have been linked to frailty, insulin resistance, cardiovascular disease, and muscle weakness(*150*, *151*); and guanidinoacetate, the immediate precursor of creatine, with reduced creatine levels associated with sarcopenia, fatigue, and cognitive decline(*152–154*).

One of the strongest protective candidates was hydroxyproline, which is not a canonical aging biomarker in humans. Hydroxyproline derives predominantly from collagen degradation and may reflect turnover of collagen-rich tissues(*155*). Hydroxyproline levels were higher in dogs with more ancestry from long-lived breeds, and were associated with lower mortality risk in individual dogs.

Not all traits influenced by breed lifespan were associated with mortality in individual dogs, or vice versa. For example, although dogs with more ancestry from short-lived breeds had lower chloride levels (β = 0.62, p = 6.5x10^-4^, p_FDR_ = 0.038; Fig. 5B), consistent with human studies showing low chloride predicts higher mortality (*156*), chloride levels in individual dogs were not associated with survival in our Cox analysis. FAD (flavin adenine dinucleotide), a key mitochondrial redox cofactor whose depletion contributes to metabolic and cellular aging (*157*), is associated with survival in individual dogs (hazard ratio = 1.37; pFDR = 0.0036), but the Breed Ancestry Score was not correlated with breed lifespan (β = −0.041; p = 0.83).

Pseudouridine showed a similar pattern: it was strongly predictive of survival in individual dogs, paralleling its association with mortality risk in humans (*26*), but the Breed Ancestry Score did not correlate with breed lifespan (β = 0.172; p = 0.29).

## Discussion

By integrating genomic, metabolic, and clinical data from thousands of dogs, we establish the domestic dog as a uniquely powerful system for dissecting the genetic and physiological architecture of aging. We present the first GWAS catalog for dogs, as well as the largest genome-wide study of blood traits, including both clinical analytes and plasma metabolites, in any nonhuman species. Many blood traits showed substantial heritability, mirroring similar studies in humans, and proved highly amenable to GWAS. Despite a study size far smaller than the typical human GWAS, more than half of the blood traits had at least one significant association.

In both dogs and humans, many blood traits are influenced by common, large-effect variants, enabling robust genetic associations even in modestly sized cohorts (*99*, *111*). In dogs, we detected strong signals with fewer than 1000 individuals, nearly an order of magnitude less than typically required in human studies(*14*), reflecting the demographic history of dog breeds. Severe bottlenecks, high inbreeding, and intense artificial selection have altered allele frequencies (*50*, *51*, *158*), substantially boosting GWAS power (*52*, *80*). This architecture positions blood traits as endophenotypes effective for linking genetic variation to complex phenotypes such as disease risk and aging in dogs.

Nearly all of the strongest dog GWAS signals for blood traits overlap loci associated with the same or closely related traits in humans, highlighting conserved biological mechanisms and reinforcing the translational utility of dogs. The small number of exceptions highlight pathways that may have diverged between dogs and humans, potentially reflecting species-specific metabolic adaptations. For example, threonine levels in dogs, but not humans, are associated with genetic variation in TDH, a gene that has evolved into a nonfunctional pseudogene in humans. Tryptophan levels in humans, but not dogs, are significantly associated with genetic variation in TDO2 (*122*), suggesting functional differences in this pathway.

Selection on aesthetic traits and genetic drift in dog breeds has reshaped metabolic pathways. Breed explains substantial variance in blood traits, significantly influencing 89% of clinical analytes and 60% of plasma metabolites. For the two most heritable traits, creatinine (h^2^_SNP_ = 0.88) and cystathionine (h^2^_SNP_ = 0.87), breed alone accounts for nearly 40% of total variance. For creatinine, this likely reflects breed differences in body size and muscle mass, with ancestry from heavier breeds predicting higher creatinine levels. However, creatinine levels are not associated with all-cause mortality risk in individual dogs (Fig. 5E), indicating that breed-driven differences in this biomarker do not directly translate into differences in lifespan. The two loci associated with cystathionine map to genes underlying breed-defining fur characteristics and are associated with fur type in our cohort, illustrating pleiotropic effects driven by selection on visible traits. Yet cystathionine levels are likewise not associated with mortality risk (Fig. 5E). This contrasts with humans, where elevated cystathionine has been linked to cardiovascular disease and mortality (*159*), and may reflect the lower prevalence of cardiovascular disease in dogs (*160*).

Pleiotropy between cystathionine metabolism and fur traits is biologically plausible. Cystathionine is a key intermediate in the transsulfuration pathway linking methionine metabolism to cysteine biosynthesis, and genetic disorders of cystathionine metabolism include symptoms of hair breakage and alopecia(*161*). Cysteine availability could influence hair structure by upregulating keratin synthesis (*162*) or by affecting disulfide bond formation that confers strength and resilience to hair (*163*). The two loci associated with both cystathionine levels and hair traits in dogs map to genes with established roles in hair biology: FGF5, which regulates hair follicle cycling (*164*), and RSPO2, which enhances Wnt/β-catenin signaling in the hair follicle niche (*165*).

By analyzing all dogs — both single- and mixed-breed — we uncover size-related influences on blood traits that were not detectable in our analyses of single-breed dogs, where body size is tightly correlated with breed. Mixed-breed dogs carry a mosaic of ancestry from multiple breeds, creating a natural experiment in which incremental contributions of ancestry from individual breeds can be evaluated. Using this framework, we found that creatinine is strongly associated with breed body size, consistent with its role as a biomarker of muscle mass (*124*). We also saw that dogs with more ancestry from large breeds tend to have higher bilirubin levels, notable given that we also show higher bilirubin predicts lower all-cause mortality risk in dogs (fig. S9), as it does in humans(*166*). In contrast, our analyses of single-breed dogs only showed no effect of weight on creatinine, bilirubin, or most other blood traits once breed was included in the model (Fig. 3C).

Leveraging the Dog Aging Project’s large, deeply phenotyped longitudinal cohort, we identify a more complex relationship between breed ancestry and aging than the simple observation that “big dogs die young”(*167*). Ancestry from both giant breeds (e.g., great Danes) and small breeds (e.g., pugs) predicts elevated globulin levels—a marker of inflammation or infection—and reduced survival (*143*) (Fig. 6A). Ancestry from brachycephalic breeds of different sizes (e.g., pugs and bulldogs) is associated with higher potassium levels (Fig. 6B), which in turn predicts reduced survival (Fig. 6D). This may reflect the consequences of congenital respiratory impairment, as chronic airway obstruction can lead to respiratory acidosis and secondary elevations in blood potassium (*168*, *169*). However, ancestry from non-brachycephalic small breeds (e.g., pomeranians) and larger breeds (e.g., Airedale terriers) is also associated with higher potassium levels(Fig. 6B), suggesting other mechanisms likely contribute.

These results highlight the potential for admixture mapping to amplify the power of dogs for genetic investigation, provided the computational challenges posed by high-dimensional ancestry can be addressed. Admixture mapping leverages local ancestry in admixed individuals to identify trait-associated loci, but existing methods are designed to model only a relatively small number of source populations. In the Dog Aging Project, modeling ancestry for 80% of dogs would require inclusion of at least 70 distinct source populations (breeds)(fig. S13). New methods capable of accommodating this high-dimensional ancestry space could further increase the power of dog GWAS, particularly for blood traits with strong breed effects, and identify the source populations contributing to lifespan-associated variation.

Together, our results demonstrate how the genetic diversity of dogs can be leveraged to identify biomarkers of aging, assess their predictive value in individuals, and pinpoint conserved physiological pathways. This dual-level approach provides a framework for mechanistic dissection of aging biology and prioritization of intervention targets. With human-like physiological complexity, and a compressed lifespan, the domestic dog is an exceptional model for both discovery and translation in aging research.

## Acknowledgments

We thank Dog Aging Project participants, their dogs, community veterinarians, and all Dog Aging Project team members. We also thank the Darwin’s Ark project participants and team members. We thank the Terra platform at the Broad Institute and the Arizona State University Research Computing Facility for computational support.

## Funding

National Institutes of Health grant U19 AG057377 (DELP)

National Institutes of Health grant R01 CA255319 (EKK)

National Institutes of Health grant R37 CA218570 (EKK)

Glenn Foundation for Medical Research

Tiny Foundation Fund at Myriad Canada

WoodNext Foundation

Dog Aging Institute

## Author contributions

Conceptualization: VS, JMA, FLC, EKK

Investigation: VS, MEW, BRH, RB, AM, KMP, DPG, KM, XL, CR, FLC, EKK

Software: VS, RB, KMP

Data Curation: VS, BRH, AM, XL, BK, CR

Resources: NSM, JMA, DELP, EKK

Supervision: NSM, JMA, DELP, FLC, EKK

Writing – original draft: VS, MEW, FLC, EKK

Writing – review & editing: VS, MEW, BRH, RB, AM, KMP, DPG, KM, XL, BK, CR, NSM, JMA, DELP, FLC, EKK

## Competing interests

DELP is on the Scientific Advisory Board for WndrHLTH Club, Inc., and serves as Treasurer of the Dog Aging Institute. KMP holds employment, advisory roles, and stock options in Colossal Biosciences (not involved in the funding, design, or execution of this study).

## Data, code, and materials availability

Raw sequencing data have been deposited in the NCBI Sequence Read Archive (SRA) under accession number PRJNA800779. All raw and processed datasets as well as GWAS summary statistics are available from Dryad (DOI: 10.5061/dryad.sxksn03hq). Code for GWAS and heritability pipeline (https://doi.org/10.5281/zenodo.15392359) and for analyses and figure generation (https://doi.org/10.5281/zenodo.18868204) are available on GitHub.

## Dog Aging Project Consortium

Joshua M. Akey¹, Rozalyn M. Anderson², Elhanan Borenstein³, Marta G. Castelhano⁴, Amanda E. Coleman⁵, Kate E. Creevy⁶, Matthew D. Dunbar⁷, Virginia R. Fajt⁸, Jessica M. Hoffman⁹, Erica C. Jonlin¹⁰, Matt Kaeberlein¹⁰, Elinor K. Karlsson¹¹, Kathleen F. Kerr¹², Jing Ma¹³, Evan L. MacLean¹⁴, Stephanie McGrath¹⁵, Natasha J. Olby¹⁶, Daniel E.L. Promislow¹⁷, May J. Reed¹⁸, Audrey Ruple¹⁹, Stephen M. Schwartz²⁰, Sandi Shrager²¹, Noah Snyder-Mackler²², M. Katherine Tolbert⁶

1. Lewis-Sigler Institute for Integrative Genomics, Princeton University, Princeton, NJ, USA
2. University of Wisconsin Madison, Madison, WI, USA
3. Department of Clinical Microbiology and Immunology, Gray Faculty of Medical and Health Sciences, Tel Aviv University, Tel Aviv, Israel
4. Cornell Veterinary Biobank, College of Veterinary Medicine, Cornell University, Ithaca, NY, USA
5. Department of Small Animal Medicine and Surgery, College of Veterinary Medicine, University of Georgia, Athens, GA, USA
6. Department of Small Animal Clinical Sciences, Texas A&M University School of Veterinary Medicine & Biomedical Sciences, College Station, TX, USA
7. Center for Studies in Demography and Ecology, University of Washington, Seattle, WA, USA
8. Department of Veterinary Physiology and Pharmacology, Texas A&M University School of Veterinary Medicine & Biomedical Sciences, College Station, TX, USA
9. Department of Biological Sciences, Augusta University, Augusta, GA, USA
10. Department of Laboratory Medicine and Pathology, University of Washington School of Medicine, Seattle, WA, USA
11. Genomics & Computational Biology, UMass Chan Medical School, Worcester, MA, USA
12. Department of Biostatistics, University of Washington, Seattle, WA, USA
13. Division of Public Health Sciences, Fred Hutchinson Cancer Research Center, Seattle, WA, USA
14. College of Veterinary Medicine, University of Arizona, Tucson, AZ, USA
15. Department of Clinical Sciences, College of Veterinary Medicine and Biomedical Sciences, Colorado State University, Ft. Collins, CO, USA
16. Department of Clinical Sciences, College of Veterinary Medicine, North Carolina State University, Raleigh, NC, USA
17. Jean Mayer USDA Human Nutrition Research Center on Aging at Tufts University, Boston, MA, USA
18. Department of Medicine, Division of Gerontology and Geriatric Medicine, University of Washington School of Medicine, Seattle, WA, USA
19. Department of Population Health Sciences, Virginia-Maryland College of Veterinary Medicine, Virginia Tech, Blacksburg, VA, USA
20. Epidemiology Program, Fred Hutchinson Cancer Center, Seattle, WA, USA
21. Collaborative Health Studies Coordinating Center, Department of Biostatistics, University of Washington, Seattle, WA, USA
22. School of Life Sciences, Arizona State University, Tempe, AZ, USA

## Notes

https://doi.org/10.5061/dryad.sxksn03hq

https://doi.org/10.5281/zenodo.15392359

https://doi.org/10.5281/ZENODO.18868204

### Competing Interest Statement

Daniel E.L. Promislow is on the Scientific Advisory Board for WndrHLTH Club, Inc., and serves as Treasurer of the Dog Aging Institute. Kathleen Morrill Pirovich holds employment, advisory roles, and stock options in Colossal Biosciences (not involved in the funding, design, or execution of this study).

### Summary of Updates

Data, code, and materials availability section of main text updated to include DOIs, since references were missing in previous submission

