## Supplementary Materials and Methods for "Blood biomarkers and breed genetics of aging in pet dogs"

Supplementary Materials for:
**Blood biomarkers and breed genetics of aging in pet dogs**

### Materials and Methods

Sample and data collection

In brief, dogs included in this genetic analysis represent the subset of Foundation companion dogs enrolled in the Dog Aging Project (DAP). Owner informed consent for participation in the Dog Aging Project was administered at the time of enrollment prior to any data or sample collection (*1*). The University of Washington IRB deemed the recruitment of dog owners for the Dog Aging Project, and the administration and content of the DAP Health and Life Experience Survey (HLES), as human subjects research that qualifies for Category 2 exempt status (IRB ID no. 5988, effective 10/30/2018). All study-related procedures involving privately owned dogs were approved by the Texas A&M University IACUC, under AUP 2021-0316 CAM (effective 12/14/2021).

Survey data were collected via the REDCap Electronic Data Capture system and veterinary electronic medical records were required to join sampled cohorts of the Dog Aging Project that involved biospecimen collection. Owners of dogs enrolled in any of these sampled cohorts received a saliva swab for DNA collection. Low-coverage (1x) whole genome sequencing is performed, and the resulting genetic data is provided to DAP for ancestry calling and genetic analysis (*1*, *2*).

A subset of 976 genotyped dogs are members of the Precision cohort with biospecimens (blood, urine, fecal, and hair samples) collected annually at their veterinary visits (*1*, *3*). All biospecimens are sent by veterinarians to the Texas Veterinary Medical Diagnostic Laboratory (TVMDL) for routine clinicopathologic assays, including conducting complete blood count (CBC) and chemistry profile testing. Approximately 0.5 mL of extracted blood is run on an Advia 120 Hematology System (Siemens Medical Solutions, Malvern, PA), and ~0.5 mL of serum is run on a DxC700AU Chemistry Analyzer (Beckman Coulter, Brea, CA) for CBC and chemistry profiling, respectively(*3*). Plasma samples were sent to the Promislow lab at the University of Washington in preparation for targeted metabolomic profiling using liquid chromatography-mass spectrometry (LC–MS) at Northwest Metabolomics Research Center (NMRC)(*3*, *4*).

Genetic Data

7,627 low-coverage sequencing FASTQ files (available on SRA under PRJNA800779) were aligned to the canFam4 (UU_Cfam_GSD_1.0 + ROSY) genome assembly using NVIDIA Clara Parabrick's (version 4.0) fq2bam wrapper for BWA-MEM on the Sol supercomputer of Arizona State University (*5*). The parameter --bwa-options='-K 100000000 -Y' was included in the pbrun fq2bam command as recommended by the Dog10K consortium's genome alignment code (https://github.com/jmkidd/dogmap). We performed genotype imputation with GLIMPSE2 (*6*). Briefly, the Dog10K phased imputation reference panel of 1929 dogs was used to create chunks and binary reference panels for each autosome (chromosomes 1 to 38). The 7,627 binary alignment map files (BAMs) were provided via a list to GLIMPSE2_phase for phasing and imputation. The resulting individual binary variant call format files (BCFs) per chunk were ligated together into a single BCF file containing the imputed genotype calls for all dogs.

Breed ancestry reference panel and ancestry calling

The breed ancestry reference panel consisted of 109 modern breeds with at least 4 dogs per breed, village dogs (4 from Nigeria, 5 from Vietnam and 55 from China), and wolves (19 North American wolves and 25 Eurasian wolves)(*2*). Breed ancestry was inferred for 7,587 dogs using ADMIXTURE (*7*), which is further described in Sexton et al(*2*) and based on the methodology developed for Darwin's Ark study (*8*). The remaining forty dogs in our genetic set are missing ancestry calls but will be available in subsequent ancestry data releases from the Dog Aging Project.

Single-breed status and standardized breed assignment

We used owner-reported breed status and breed labels as the primary source of breed information, with limited corrections and validation using sequencing-based ancestry. Owner-reported breed names were standardized and harmonized to correct common misspellings and near-duplicate labels. Dogs were classified as single-breed if owners reported them as such, or if ≥85% of genetic ancestry was assigned to a single breed. For dogs classified as single-breed, the standardized breed (Data S14) was the cleaned owner-reported breed, with two exceptions: First, when owner-reported and sequencing-derived breeds differed only by size variant (e.g., poodle vs. poodle (toy)) and genetic ancestry overwhelmingly supported one breed (≥85%), we assigned the sequencing-derived breed. Second, for dogs reported by owners as mixed-breed but with ≥85% of genetic ancestry assigned to a single breed, we assigned that sequencing-derived breed as the standardized breed.

Size distribution

To assess whether body size distributions were consistent with a single mode, we applied Hartigan's dip test for unimodality to each phenotype (R package diptest), treating a dip-test $P < 0.05$ as evidence against unimodality.

To quantify how well breed ancestry predicts body weight, we analyzed the relationship between ancestry-derived predicted weight and observed body weight in dogs > 2 years old with age, sex, sterilization status and body weight information. Predicted body weight was calculated as the sum across contributing breeds of relative ancestry proportion (assigned as 1 for single-breed dogs) multiplied by breed-specific weight estimates (table S12). Linear regression modeled observed weight as a function of predicted weight, sex, and sterilization status, with model fit summarized by $R^2$. Significance of the predicted-weight effect was assessed using Freedman–Lane permutation tests (100,000 permutations, fixed seed, +1 correction).

Criteria for phenotype inclusion

We retained ordinal and binary (yes/no) survey questions. We excluded questions if they were free-text, nested, dog and owner demographics, and environmental questions. Questions that had greater than 15% missingness (NAs) as well as ordinal survey questions with unbalanced phenotype distributions, defined as a single answer having at least 85% of the responses, were also excluded. There were a few exceptions to the criteria mentioned above where we included questions that could influence dog health or be affected by the intrinsic characteristics of the dog, which may be impacted by genetics. Quantitative survey questions with a heavy-tailed distribution (kurtosis > 7) were also excluded. The inclusion criteria and reasoning for individual survey questions can be found in Data S1. Similarly, we excluded 14 metabolites and 6 blood analytes (1 complete blood count and 5 blood chemistry analytes) with heavy-tailed distributions (kurtosis > 10) from further analysis. Kurtosis results can be found in DataDryad.

Data cleaning

Survey data from the 2022 Dog Aging Project Data Release were re-coded to ensure all ordinal question responses were in logical order. The lowest response for any particular question was coded as 0 with the highest response coded as n-1, where n = the total number of response options. "Unknown" responses were uninformative and therefore converted to missing values (NAs). The survey responses were re-coded for genetic analysis where DAP_coded_response values were converted to the coded_response column values available in Data S16. After cleaning, we had a total of 1022 survey questions available for analysis. We excluded 868 questions for the following reasons: high levels of missingness with many cases arising from nested survey questions that were not shown to dog owners based on their response to a prior question (N = 480), demographic or environmental (N = 197), responses were the same for many dogs leading to insufficient phenotypic variability (N = 32), and responses not suitable for GWAS (e.g. free text, categorical, or nested; N = 159). The specific reasoning for exclusion of each individual question can be found in Data S1.

After re-coding survey responses, they were z-transformed using the *scale* function in R. Raw complete blood count (CBC) data were square-root transformed, and serum chemistry profile (SCP) data were natural log transformed. For metabolites, the LC-MS peaks were integrated to give metabolite count data that were then data transformed. Residuals of a regression model were taken to correct both main effects of batch and run-order effects, which ultimately resulted in normalized peak areas(*4*). Each metabolite was scaled to unit variance by batch. Missing values were imputed by 10-nearest neighbor mean-imputation, and the effects of the remaining technical covariates, travel time, hemolysis and arrival temperature on metabolite abundance were corrected for using linear regression(*4*). This data processing resulted in normalized baseline metabolite measurements for 936 dogs(*4*).

Heritability and GWAS

We developed our heritability and GWAS workflow using the Nextflow workflow management system to facilitate reproducibility when using different computing environments and scalability by parallelizing across multiple phenotypes. The software used in the workflow is Plink (v1.90b6.21 and v2.00a5LM) and Genome-wide Complex Trait Analysis (GCTA v1.94.1). The workflow includes genetic set quality control, followed by SNP-based heritability estimation using restricted maximum likelihood (REML), and GWAS via mixed linear model leave-one-chromosome-out approach. It follows the methodology performed in the genetic analysis of Darwin's Ark (*8*). The code can be found at https://github.com/VistaSohrab/dog-gwas-heritability-nextflow and each step from Genetic set filtering until GWAS is described in more detail below.

Filtering and GRM construction

We first filter the PLINK genetic set input file based on the params.maf (minor allele frequency threshold), params.geno (genotyping rate threshold), and params.hwe (Hardy-Weinberg equilibrium deviation p-value threshold) declared in nextflow script or provided with

"--maf", "--geno", and "--hwe" in nextflow run command at time of submission. Biallelic SNPs with minor allele frequency less than 0.01 and missing in greater than 5% of individuals were excluded. SNPs with extreme deviation from Hardy-Weinberg equilibrium, given p-values below $1x10^{-20}$ in the exact test with mid-p adjustment, and excess heterozygosity were excluded. The following nextflow parameters were used: params.maf = 0.01, params.geno = 0.05, params.hwe = 1e-20, as already defined in the main script dog_gwas_heritability_multiple_phenotypes.nf. After quality control of genotypic data, we retained 9,867,131 autosomal biallelic SNPs for 7,627 dogs. We then construct the genetic relatedness matrix (GRM) from the filtered PLINK genetic set files created. We calculated a GRM for the 7,627 sequenced dogs in our genetic set of 9,867,131 autosomal biallelic SNPs.

Heritability analysis

We estimate SNP-based heritability ($h^2_{SNP}$) while accounting for linkage disequilibrium to avoid inflation by non-independence of linked SNPs (LD-corrected SNP-based heritability). LD scores were calculated in 250 kb regions using a block size of 10,000 kb with an overlap of 5,000 kb between blocks. Four LD-stratified genetic relationship matrices (GRM) were generated for 7,627 dogs upon LD score stratification of SNPs. We calculated $h^2_{SNP}$ with standard errors using the GREML-LDMS method incorporating multiple GRMs corresponding to sets of LD-score SNPs stratified by quartiles (*9*). However, LD correction was not applied to 76 traits where restricted maximum likelihood failed to converge or more than half of the LD-stratified variance components were constrained. Therefore, we also conducted the standard GREML heritability analysis using the single GRM constructed from 9,867,131 autosomal SNPs with minor allele frequency (MAF) >1%. All models included covariates for age, weight, and sex status (sex and sterilization status) with hours fasted also included for blood traits.

GWAS

We performed genome-wide mixed linear model-based association on autosomal chromosomes using the "leave-one-chromosome-out" approach (MLMA-LOCO) implemented in GCTA (*10*). Sex and sterilization status (sex_class_at_HLES variable in the 2022 Dog Aging Project DogOverview dataset) was the categorical covariate while age and weight at time of survey completion were quantitative covariates for survey-based phenotypes. When running GWAS on weight, age was the only quantitative covariate. For blood-based phenotypes, the categorical covariate included was sex and sterilization status (sex_class_at_HLES variable in 2023 Dog Aging Project DogOverview dataset), while quantitative covariates included age at date of blood collection, baseline weight at time of survey completion, and the number of hours fasted prior to blood collection.

We defined associations by clumping SNPs in LD ($r^2 > 0.2$) (params.clump_rsquared = 0.2) that were within 250 kb (params.clump_kb = 250) of the associated index SNPs, defined as having p-value less than 1e-6 (params.clump_pval = 1e-6).

PheWeb

PheWeb is designed to display human GWAS, supporting the display of 23 autosomes; therefore, the code was adapted to allow the display of 38 autosomes. RSIDs do not exist in dogs and therefore were removed as a requirement, and hyperlinks to the UCSC genome browser point to the corresponding canFam4 UCSC genome browser for users to view the genomic region a variant of interest resides in. These changes to standard PheWeb are available as a github repository here: https://github.com/AkeyLab/DAP_pheweb.

PheWeb thresholds for genome-wide significance ($p = 5 \times 10^{-8}$) and suggestive associations ($p = 1 \times 10^{-6}$) conventionally used in human GWASs (*11*, *12*) remain unchanged. The search bar in the phenotypes page can be used to filter the table by phenotype name, sample size number, gene name, and phenotype categories. The phenotype categories that can be searched include the following survey section names: Dog Demographics, Dog Behavior, Dog Environment, Physical Activity, Health Status, Medication and Preventatives, Diet, Eating Behavior, and Dog Owner Relationship. The molecular phenotypes can be searched via Complete Blood Count or Blood Chemistry Panel, while metabolites have further classifications such as amino acids and peptides, purines, pyrimidines, fatty acids, and fatty esters that can be used to filter the phenotype table on PheWeb.

### Defining blood trait GWAS regions in dogs

All analyses were performed in R (version 4.5.2). We included all suggestive dog GWAS regions ($p < 1x10^{-6}$) for phenotypes classified as Clinical analytes or Plasma metabolites in Table S1, using regions defined in Data S3. Regions shorter than 1 kb were padded to 1 kb, yielding 1,133 dog region–trait pairs. These regions were lifted from the dog reference genome (CanFam4.0) to the human genome (hg38) using UCSC liftOver with the chain file canFam4ToHg38.over.chain.gz and parameters -minMatch=0.1 (13, 14). Of the 1,133 regions, 1,091 (96.4%) successfully lifted. Lifted regions were then expanded to a total width of 500 kb.

### Defining blood trait GWAS regions in humans

Human associations were extracted from the NHGRI–EBI GWAS Catalog file gwas_catalog_v1.0-associations_e113_r2025-02-08.tsv downloaded from https://www.ebi.ac.uk/gwas/docs/file-downloads. To restrict the analysis to blood and metabolic traits measured in non-disease-affected adults, we filtered this set to 304,480 significant ($p < 5x10^{-8}$) associations reported in a manually curated list of 25 publications with metabolomic GWAS results for healthy individuals (*13–37*). We further filtered the list down to 186,964 associations by keeping only blood traits measured in non-diseased individuals (Data S4). We generated more uniform and consistent trait names by processing the original names through the ChatGPT API ("gpt-4.1-mini") using the system prompt shown below. This is implemented in run.GET_HUMAN_GWAS_NAMES.R.

### Prompt used for human blood trait standardization

"You classify trait names measured in blood and return a canonical name and an include flag.
Workflow: (1) Decide whether INCLUDE is TRUE or FALSE; (2) if INCLUDE = FALSE, do not refine the name, set canonical_name equal to the input and provide a brief reason; (3) if INCLUDE = TRUE, generate a cleaned, standardized canonical_name.
INCLUDE = TRUE only for direct blood measurements: serum or plasma metabolites (small molecules, amino acids, lipids, acylcarnitines); clinical chemistry analytes (e.g., creatinine, urea, glucose, calcium, bilirubin, ALT, AST); complete blood count traits (e.g., RBC, WBC,

neutrophils, eosinophils, platelets, hematocrit, MCV); serum or plasma proteins, cytokines, complement factors, and hormones.
INCLUDE = FALSE for non-blood or context-specific traits: urine, CSF, tissue, or fecal measurements; disease-specific or cohort-qualified traits (e.g., "in chronic kidney disease," "in elite athletes"); traits not measured in blood (e.g., height, diabetes, bone density, psychiatric traits); unknown feature identifiers (e.g., X-11787, QI7389); environmental, behavioral, or questionnaire-based traits.
Canonical name rules (applied only when INCLUDE = TRUE): normalize to the underlying biochemical or clinical entity; remove measurement context terms (e.g., serum, plasma, levels, concentrations); remove transformations (e.g., minimum, maximum, mean, inverse-normal transformed); remove disease qualifiers; for complex lipids, remove chain-length or unsaturation annotations (e.g., 38:3, 18:1/20:4, O-36:2) and retain only the lipid class; collapse protein isoforms, subfamilies, and member numbers to the core family name; keep chemical names unchanged; if ambiguous but clearly a blood measurement, select the simplest reasonable canonical name.
Special case when INCLUDE = FALSE: set canonical_name equal to the input and provide a brief explanation for exclusion.
Output format: return a single JSON object containing the input trait name, canonical_name, include flag, and reason."

Defining human/dog overlap

We identified 21,100 overlaps between the 1,091 lifted dog regions and the 186,964 human GWAS intervals using bedtools intersectBed (v2.31.1)(*38*). The resulting table is provided as Data S4. Out of 1133 suggestive region/trait pairs found in dogs, 1091 lifted over and 501 overlapped a significant association in humans. The total number of overlaps was 21,732. This is implemented in script run.GWAS_INTERSECT.R.

We characterized relationships between dog and human traits with overlapping GWAS signals using the OpenAI Chat Completions API (model gpt-5.1) as a structured classification engine via the httr2 and jsonlite packages in R. We provided a detailed system prompt that defined a strict 1–5 scale describing the biological relationship between two blood-measured traits: (1) same measure or strict superset/subset, (2) same specific metabolic or hematologic pathway or direct component (including enzyme–substrate/cofactor and direct clinical readouts), (3) functionally related but not direct pathway neighbors, (4) systemic or state-level correlation only, and (5) no clear relationship. Each request included a single trait pair and instructed the model to return only a JSON object containing the fields trait1, trait2, score, and a free-text rationale. To maximize efficiency, we scored overlaps with the strongest associations in human GWAS first and evaluated weaker associations only when no score of 1 or 2 was identified for the corresponding strong association. In total, 1,863 trait pairs were scored (37 score=1; 137 score=2; 746 score=3; 839 score=4; 104 score=5) (see S_TRAIT_MATCH.tsv in DataDryad). We then identified all overlaps (see S_GWAS_RAW_OVERLAP.tsv in DataDryad) where the traits in dog and human are related to one another. These results are in Data S4, and in Fig. 2 panel D, with pairs with scores of ≤2 described as "same entity or directly related" and scores of ≤3 described as "functionally related". Scores of 4 or 5 are not considered as matches.

Prompt used for cross-species trait relationship scoring

"You score pairs of blood-measured biological traits on a strict 1–5 scale describing their biological relationship.
Score definitions:
1 = Same measure: identical molecule or entity, or a strict subset/superset; identical text ignoring case and whitespace; aggregate versus component of the same measure (e.g., total WBC vs neutrophils; BUN vs urea).
2 = Same specific pathway or direct component: traits in the same defined metabolic or hematologic pathway, including substrate–product relationships within two steps; shared canonical pathway modules (e.g., glycolysis, TCA cycle, pentose phosphate pathway, urea cycle, β-oxidation, carnitine shuttle, bile acid synthesis or transport, amino acid catabolism including BCAA, glutathione cycle, tryptophan–kynurenine or indole pathways, one-carbon or methylation cycle, heme–bilirubin metabolism, purine or pyrimidine metabolism); enzyme–substrate or enzyme–cofactor pairs; metabolite–conjugate pairs; metabolite–direct clinical chemistry readouts; direct lineage relationships (e.g., total WBC vs lymphocytes; RBC vs reticulocytes); metabolite–primary carrier or transport protein in blood (e.g., fatty acids vs albumin).
3 = Functionally related: traits involved in the same biological process but not direct pathway neighbors, such as oxidative stress, immune activation, thrombosis, gut–liver axis function, amino acid homeostasis, or red blood cell turnover.
4 = Systemic or state-level correlation: traits that covary due to shared physiological state, including renal or hepatic function, inflammation, cardiometabolic status, nutritional state, or disease severity, without a direct mechanistic pathway connection.
5 = No clear relationship: none of the above criteria apply.
Quick scoring checks: score 1 for identical or strict component/superset relationships; score 2 for shared defined pathway modules, enzyme–substrate or cofactor relationships, direct clinical readouts, or primary carrier–metabolite pairs; score 3 for shared biological processes without close pathway linkage; score 4 for systemic correlation only; score 5 otherwise.
Output format: return only a JSON object containing trait1, trait2, score, and a brief rationale."

<u>ANOVA on blood traits in single breed dogs</u>

We fit linear models for each blood trait (CBC, SCP, and metabolites), with breed as the primary factor of interest across all single breed dogs in R v4.4.3. Breeds with at least 2 or more representative dogs were retained in the analysis. Age, weight, sex and sterilization status were also included as covariates. We assessed the significance of each predictor using an analysis of variance (ANOVA). We used the anova_test function of the Rstatix package v0.7.3 with the following formula: blood trait ~ age + breed + weight + sex status. We also ran ANOVA excluding breed from the formula (blood trait ~ age + weight + sex status) to understand the effects of the other covariates in the absence of breed. ANOVA results are in Data S5.

To compare how strongly different predictors (breed, age, sex, and body weight) contributed to variation in blood-based traits, we analyzed the ANOVA-derived proportion of variance explained (ges) using a linear mixed-effects model implemented in the R packages lme4 v1.1.37 and lmerTest v3.1.3. The model treated Effect (breed, age, sex, weight) as a fixed effect and included a random intercept for phenotype to account for the repeated-measures structure (each phenotype contributes four variance components). Estimated marginal means and pairwise contrasts were obtained using emmeans v2.0.0, including focused comparisons of breed versus each alternative predictor. Statistical significance of fixed effects and contrasts was based on

Satterthwaite-adjusted degrees of freedom, and p-values reported in the text reflect these mixed-model estimates.

To evaluate how strongly variance explained (ges) depended on trait heritability (her) for each predictor, we used the same linear mixed-effects model, and included fixed effects for heritability, predictor identity (Effect: breed, age, sex, weight), and their interaction, with phenotype included as a random intercept (lmer(ges ~ her * Effect + (1 | phenotype))). Marginal slopes of ges versus her for each predictor were estimated using emmeans (emtrends), providing effect-specific trends and confidence intervals.

Regional association of RSPO2 and FGF5 with fur traits in 50 dogs

Photographs of 50 dogs with blood traits were used to phenotype their furnishings, fur length, and fur curl. We ran quantitative regional association analysis for fur length and case-control regional association analysis with Fisher's exact test for furnishings and fur curl using PLINK v1.90b7.2 64-bit (11 Dec 2023). The RSPO2 region was defined as --chr 13 --from-bp 6400000 --to-bp 11400000, while the FGF5 region spanned --chr 32 --from-bp 33000000 --to-bp 38000000. Regional association results are reported in Data S6. For the top 6 SNPs associated with cystathionine levels in RSPO2 and FGF5, we calculated breed allele frequencies across 3,594 single-breed dogs from 49 breeds that have at least 20 representative individuals with PLINK v1.90b7.2 64-bit (11 Dec 2023). Breed allele frequencies can be found in Data S7.

Linear Mixed Effects Regression Analysis (LMER)

To measure the relationship of genetic breed ancestry with blood traits, we constructed LMER models for all dogs, including both single-breed dogs and those with mixed breed ancestry in R v4.4.0 (r-4.4.0-gcc-12.1.0) using the lme4_1.1-35.5 and lme4qtl_0.2.2 packages. We included only breeds that have a standard deviation of ≥5% ancestry across dogs carrying that breed. For each LMER model, the blood trait of interest was the response variable with breed ancestries as fixed effects and age tertiles and relatedness derived from the genetic relatedness matrix (GRM) as random effects. These models were built with REML to obtain unbiased estimates, standard deviations, and Wald statistics (t.val, or Breed Ancestry Score) for the fixed effects of breed on blood traits with ANOVA performed to obtain the breed F statistics. To obtain the likelihood ratio for each breed, we constructed models using maximum likelihood (ML) with and without the breed. We then performed ANOVA to report p-values for the likelihood tests conducted between the ML models. These results can be found in Data S8. The LMER methodology and implementation was similar to that performed for Darwin's Ark(*8*).

Relating breed-level genetic effects to body size and lifespan

To quantify whether breed-level associations between genetic breed ancestry and blood traits are related to breed body size and lifespan, we analyzed breed-specific LMER summary statistics from Data S8. For each phenotype, we modeled the breed-level association strength (REML.t.val, or Breed Ancestry Score) as a function of breed standard body weight (breed.weight.kg) and breed median lifespan(*39*) using (i) an ANOVA framework (rstatix) and (ii) linear regression on standardized variables to obtain standardized effect sizes and 95% confidence intervals. Analyses were performed separately for each phenotype after harmonizing breed names across tables and excluding missing values. P-values from ANOVA were adjusted for multiple testing using the Benjamini–Hochberg procedure within each model term (Effect) across phenotypes. Results are in Data S9.

Survival analysis

We performed time-dependent Cox proportional hazard modeling for all blood traits using the coxph function of the survival package v3.8-3 in R v4.4.3. We analyzed time-to-death of dogs with the initial timepoint ($t_0$) referring to the date of initial blood draw. Observations were censored at either the last blood draw or survey collection timepoint if the dog was known to be alive. We adjusted for dogs' scaled age, sex and either scaled weight at $t_0$ for metabolites or weight category at time of enrollment in the Dog Aging Project for clinical analytes as fixed effects with scaled blood trait values as time-varying covariates updated at each annual follow-up blood draw. We further adjusted the resulting p-values using the Benjamini-Hochberg method. Cox regression results are in Data S10.

Inclusion of breed ancestry score as fixed effect for risk and protective blood traits

For two of the risk-associated and three protective blood traits, we also ran the Cox model with an additional fixed effect referred to as the composite ancestry score, which was the sum of ancestry from the top nine most strongly associated breeds with the analyte. The composite ancestry score was derived by summing the proportion ancestry from the top 9 high-risk breed ancestries, defined as the 9 breeds with the highest absolute value of the REML t-value (REML.tval) having p-value (ML.anova.pvalue) < 0.05, for each dog.

Annotation of GH/IGF-1 metabolites

To annotate whether circulating metabolites are affected by perturbation of the growth hormone/insulin-like growth factor-1 (GH/IGF-1) axis, we curated published metabolomic studies of rodent models with established alterations in GH/IGF-1 signaling using a semi-manual literature review that focused on validated GH/IGF-1–perturbed mouse models (*40*). Studies were included if they reported targeted or untargeted metabolomic measurements of at least one of the 123 plasma metabolites analyzed in this study (Table S1), measured in plasma or serum. Tissue-only studies were excluded.

We identified relevant studies that used one of the following validated GH/IGF-1–perturbed rodent models (*40*): Ames dwarf (Prop1–/–); Snell dwarf (Pit1–/–); GHRKO (Ghr–/–); GHKO (Gh–/–); GHRH-KO; GHRHR-KO (lit/lit); adult-onset GHRKO (aGHRKO); GHA mice; bGH transgenic (MT1-bGH); PEPCK-bGH; and GHRH-overexpressing mice. We using a semi-manual approach that combined manual search with AI-assisted tools:

Manual searches with the Google Scholar and PubMed search platforms, used the following search string: (metabolomics OR "targeted metabolomics" OR "untargeted metabolomics" OR "metabotyping") AND (plasma OR serum) AND ("growth hormone" OR GH OR IGF-1) AND (mouse OR mice OR rodent) AND (GHRKO OR GHRH OR "growth hormone receptor" OR "GHRH knockout" OR "growth hormone transgenic" OR Ames OR Snell OR bGH). Subsequent search strings combined a list of 15 specific GH/IGF-1 mouse model names with terms related to metabolomics and blood samples. Example: "GHRKO model" AND (plasma OR serum) AND (metabolomics).

AI-assisted searches using the DeepResearch AI platform (OpenAI GPT-5.2, Dec 10th, 2025) and Google Scholar Labs (Dec 10th, 2025) using the following prompt:

"Conduct a systematic literature review to identify studies that report individual metabolite changes in plasma or serum in the following transgenic rodent models with perturbed GH/IGF-1 activity: Ames dwarf (Prop1–/–); Snell dwarf (Pit1–/–); GHRKO (Ghr–/–); GHKO (Gh–/–);

GHRH-KO; GHRHR-KO (lit/lit); Adult-onset GHRKO (aGHRKO); GHA mice; bGH transgenic (MT1-bGH); PEPCK-bGH; GHRH-overexpressing; any other transgenic rodent model with documented alteration of the GH/IGF-1 axis. Include all peer-reviewed studies up to Dec 10th, 2025 and provide citations.". Five studies met inclusion criteria (*41–45*).

Integrating breed-level effects with individual-level mortality associations

We combined the breed-ancestry standardized effect estimates and p-values from Data S9 (model REML.t.val ~ breed.weight.kg + lifespan) and the Cox proportional hazard ratios and p-values from Data S10 and computed signed z-scores for breed-level effects (z_lm) and mortality associations (z_cox) by combining two-sided p-values with the sign of the estimated effect. To quantify concordance between breed-level and individual-level signals, we calculated a combined z-statistic z_comb = (-z_lm + z_cox)/sqrt(2), from which we derived two-sided p-values and Benjamini–Hochberg FDR-adjusted q-values. To assess whether concordance depended on whether the trait was a plasma metabolite or clinical analyte, we regressed z_lm and z_cox on phenotype category, correlated the resulting residuals using Spearman's ρ within each model Effect, and summarized n, ρ, and p-value. The final integrated table (phenotype, Effect, z_lm, z_cox, support score, combined statistics, and FDR) is in Data S11.

### Supplementary Text

Validation of low-pass sequencing + imputation in dogs

Low-pass sequencing with imputation provides substantially higher marker density in dogs compared to available genotyping arrays, making it essential for GWAS in highly admixed and diverse cohorts where linkage disequilibrium (LD) is short. In previous work(*8*), we showed that genotyping arrays capture only 40–80% of genomic diversity in mixed-breed dogs, whereas low-pass sequencing followed by imputation tags over 94% of variants. Our original Darwin's Ark dataset used loimpute (*46*) for imputation. We sequenced dogs to a depth of 1.0x ± 0.6x (±SD) and imputed against a reference panel of 435 canids with mean coverage 22.9x. In that dataset, we found 98.3% ± 0.7% concordance between low-pass and high-coverage (>30x) whole-genome sequencing for SNPs with minor allele frequency (MAF) > 0.02.
Here, we sequenced dogs to slightly lower depth of 0.79x ± 0.41x (±SD), and imputed genotypes with the Dog10K reference panel(*47*) using GLIMPSE2(*6*). Dog10K includes 1,611 dogs from 321 breeds, 309 village dogs, and 9 wolves, including the dogs from our original imputation panel. Thus, our new imputation reference panel is bigger and captures more diversity than our original imputation panel. The breeds in Dog10K include 85% of the breeds in Darwin's Ark dogs and collectively represent 86% of Darwin's Ark breed ancestry. The large number of breeds in our imputation reference panel means we likely capture nearly all common variation in any missing breeds, as genetic variants private to a single breed are rare and low frequency, reflecting the recent shared ancestry among breeds(*8*, *48*).

Comparison of sterilized and intact dogs

The reproductively intact and sterilized dogs had different population demographics. While the sterilized dogs were more representative of the pet dog population as a whole, the intact dogs were enriched for young, single-breed animals. Most reproductively intact dogs were single-breed (85.5%) and male (59%). Less than half of sterilized dogs were single-breed (41%)

and 49% were male. The intact dogs were also significantly younger than the sterilized dogs under two years of age at enrollment (61%; mean age 2.6 years versus 5.0 years in the full cohort; t-test $p = 1.68 \times 10^{-151}$).

To maximize our sample size, we included all dogs in all analyses. In the Cox regression analysis, sex and sterilization status were assessed separately, and as sterilization status was not significant, only sex was included as one of the covariates in survival analyses. In all other analyses that included sex as a covariate, we also included sterilization status by coding sex as: Male_neutered, Male_intact, Female_spayed, Female_intact.

Survival analysis accounting for breed ancestry

To confirm that the individual analytes were predictive of mortality after accounting for breed ancestry, we reran the Cox model with a composite ancestry score that was the sum of ancestry from the top nine most strongly associated breeds with a particular analyte. The strongly associated breeds were selected from those having the greatest absolute value Breed Ancestry Score (REML.t.val) and significantly associated with the blood trait (ML.anova.p $< 0.05$). Composite ancestry score was not significantly associated with survival for any of the blood traits tested. All traits remained significantly associated with risk of death even after accounting for breed composition (Figure S10).

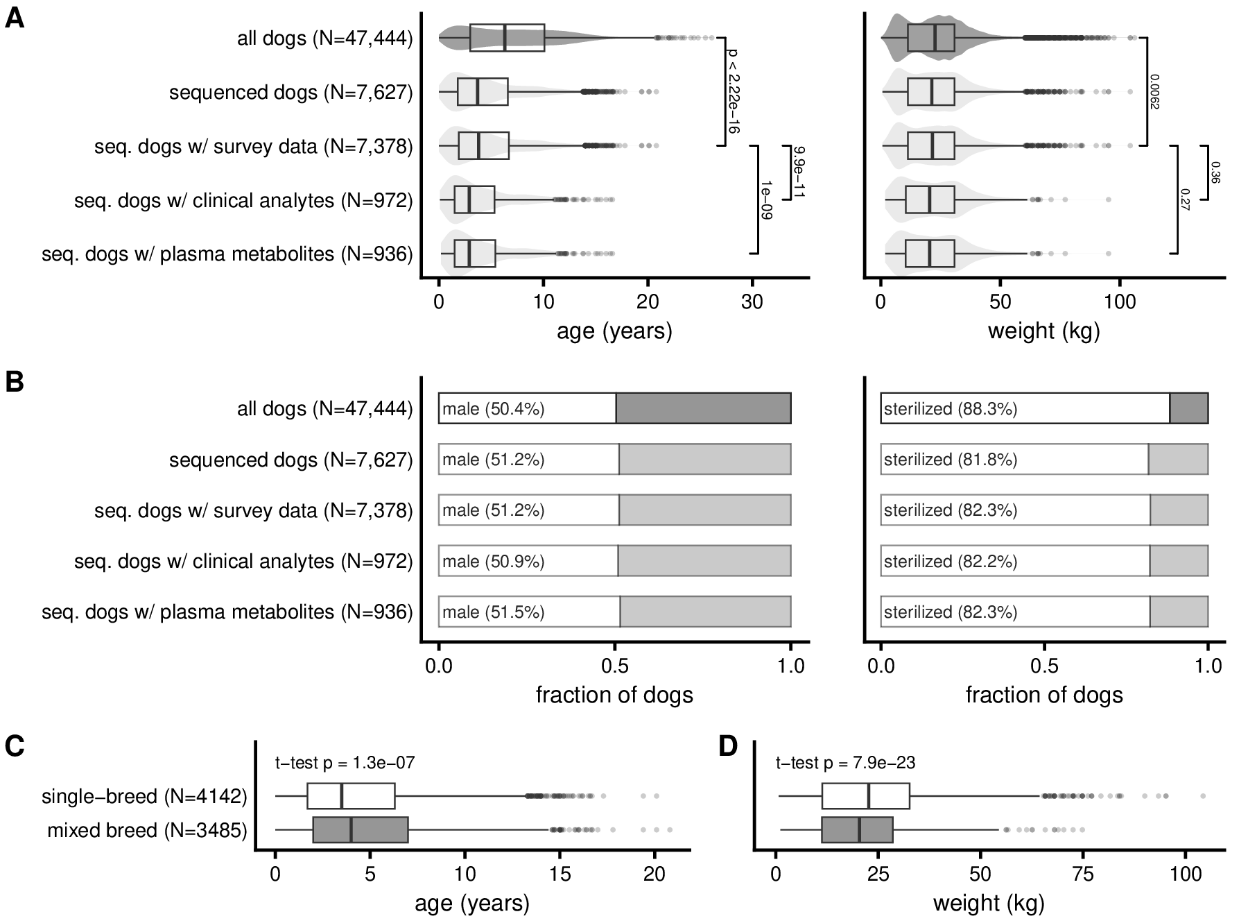

**Fig. S1. Demographic characteristics of sequenced and surveyed dogs from the Dog Aging Project. (A)** Violin and box plots show the distribution of age and weight across subsets of dogs, including all Dog Aging Project (DAP) dogs, dogs with genome sequence data, and subsets with additional phenotype data (e.g., survey data, CBC, serum chemistry, plasma metabolites) at enrollment. P values were computed using two-sided Wilcoxon rank-sum tests. **(B)** Proportions of male and sterilized dogs are shown as stacked bars; group differences were assessed using $\chi^2$ tests. **(C)** Single-breed dogs are slightly younger than mixed-breed dogs ($t = 5.28$, $df = 7301.5$, $p = 1.3 \times 10^{-7}$). **(D)** Single-breed dogs are heavier than mixed-breed dogs ($t = -9.87$, $df = 7570.1$, $p = 7.9 \times 10^{-23}$). Box plots indicate medians and interquartile ranges.

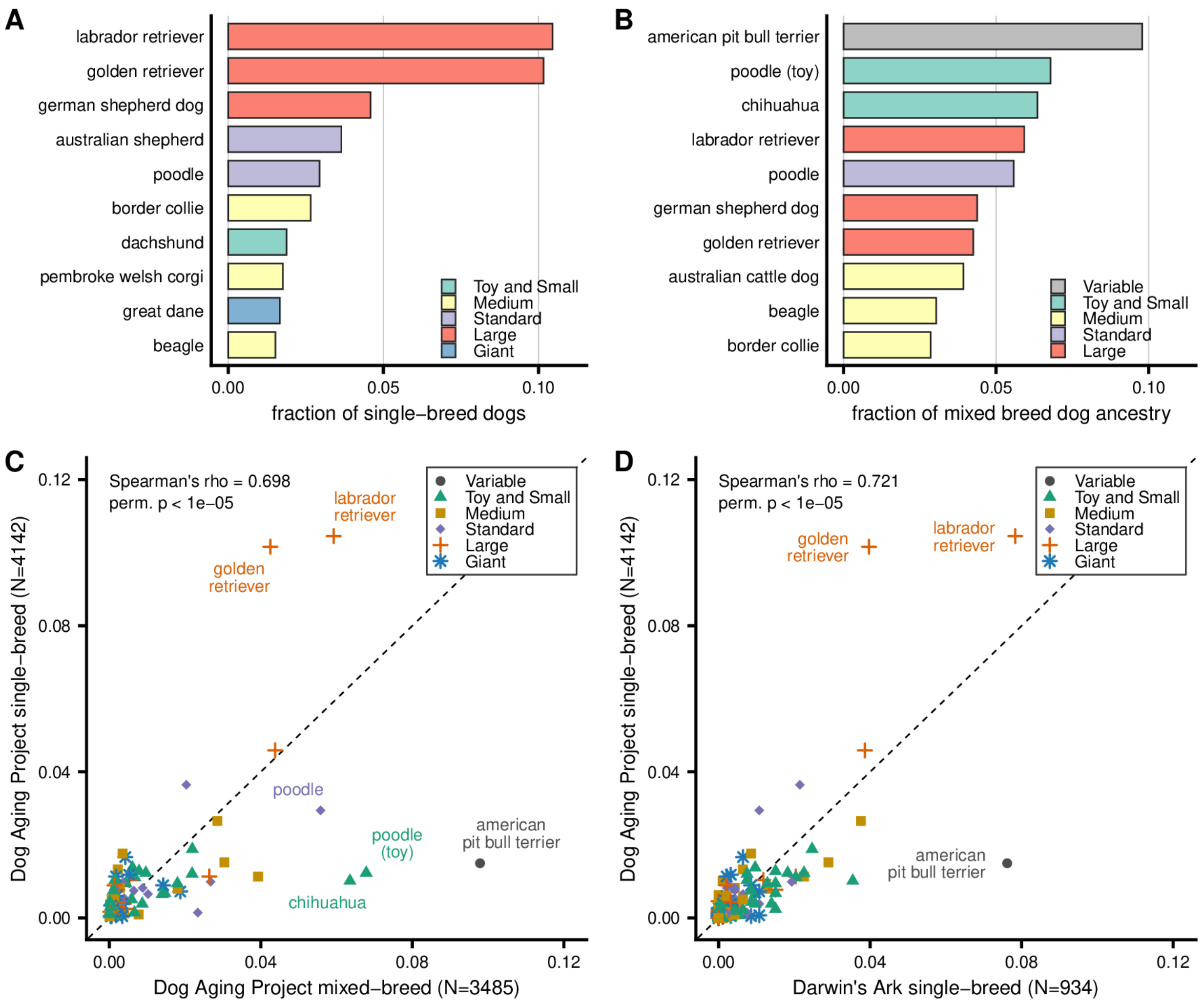

**Fig. S2. Breed composition of the Dog Aging Project cohort. (A)** The ten most common single-breed dogs in the sequenced Dog Aging Project cohort, shown as a fraction of all single-breed dogs. Bars are colored by breed size classification. **(B)** The ten most common breed ancestries among mixed-breed dogs, based on aggregated genetic ancestry proportions. **(C)** Correlation between breed frequencies among single-breed dogs and mixed-breed dogs in the Dog Aging Project and **(D)** between single-breed dogs in the Dog Aging Project and Darwin's Ark cohorts. Each point represents a breed. The dashed line indicates a 1:1 correspondence. Breed labels are shown for those with >4% frequency in the DAP cohort. Perm. p is p value from two-sided permutation test with 100,000 permutations.

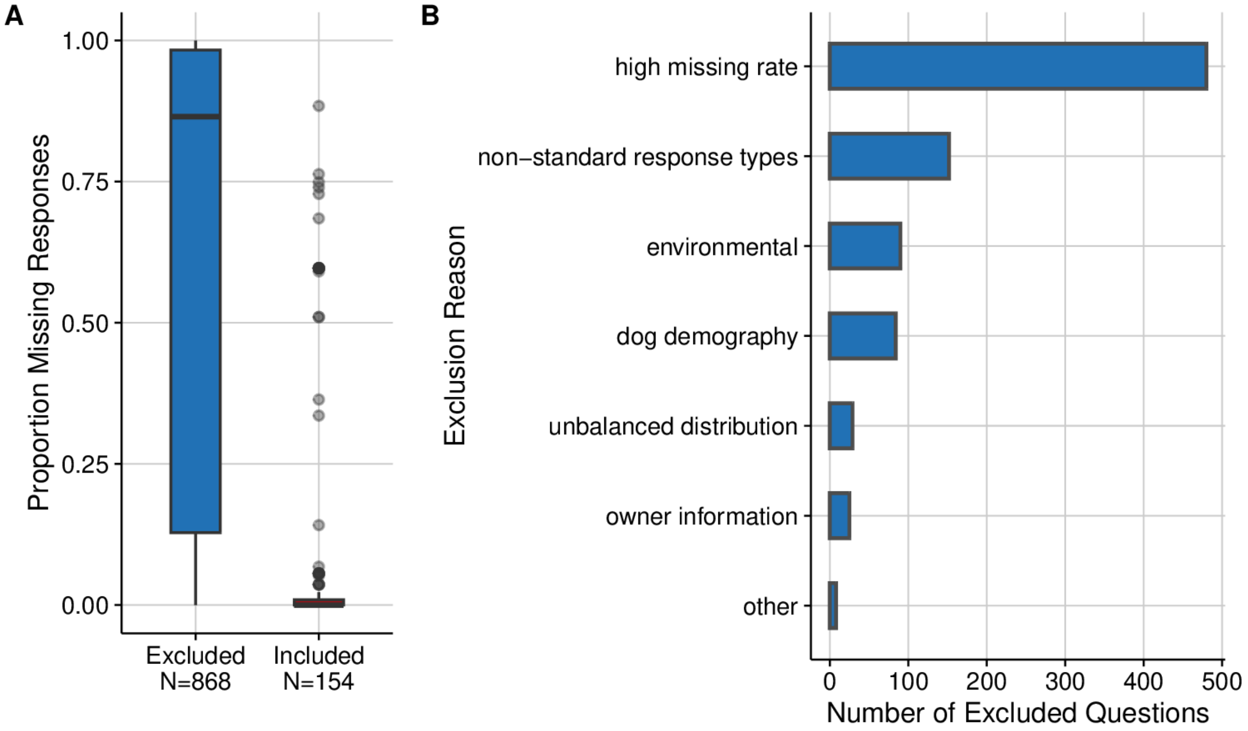

**Fig. S3. Criteria for excluding survey questions from genetic analyses. (A)** The proportion of missing responses (questions with NA values) across baseline Dog Aging Project survey questions for those excluded and included in our genetic analysis. Box plots show median and interquartile ranges. Points indicate outliers as depicted in our included survey questions that have high proportion missingness. **(B)** Bar chart showing exclusion reasonings and the number of excluded questions that fall into each category for the 868 excluded questions.

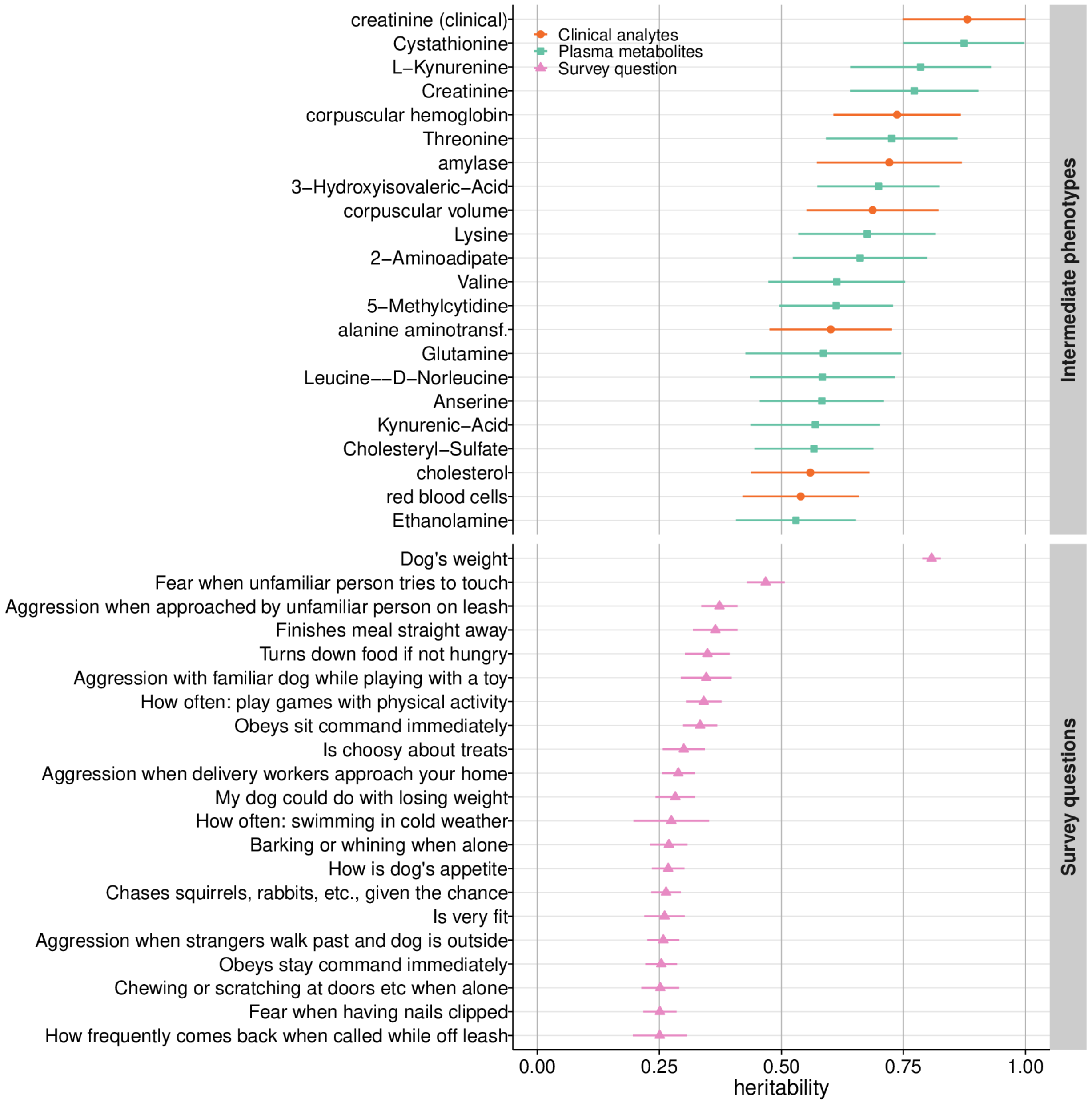

**Fig. S4. Most heritable traits in the Dog Aging Project cohort.** We estimated SNP-based heritability ($h^2$) and standard error for 313 traits using GREML with LD score correction. Traits are grouped as either Survey questions or Intermediate phenotypes (blood traits), and only those in the top 20% of heritability within each group are shown. Points represent $h^2$ estimates; horizontal lines indicate standard errors. Trait labels are ordered by decreasing heritability. Categories are colored and shaped by phenotype category. Traits related to dog-owner relationship (mdors) and environment (prefix de_) were excluded.

**A**

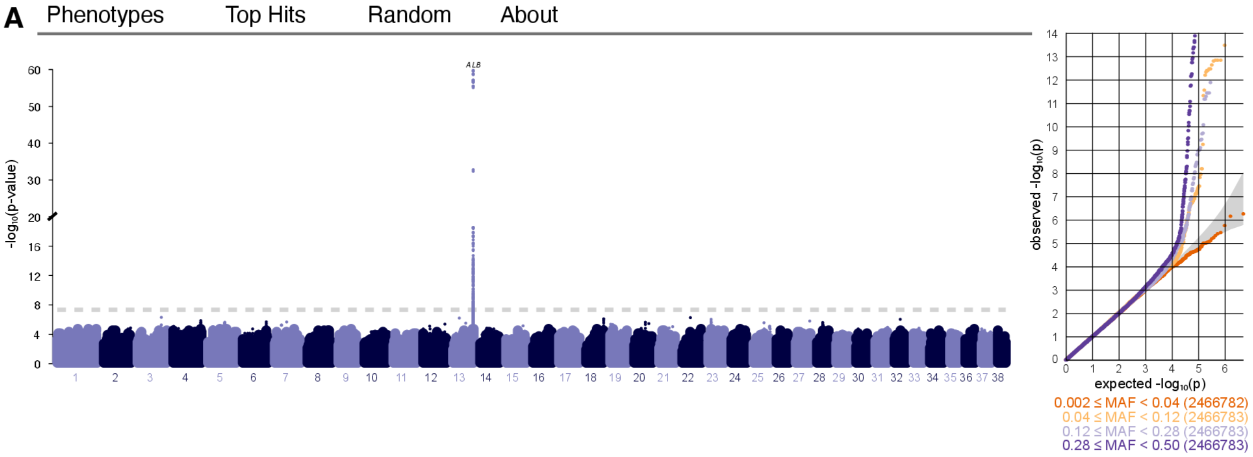

| Variant | Nearest Genes | MAF | P-value | Effect size |
|---|---|---|---|---|
| 13:63,214,897 T / A | ALB | 0.34 | 1.8e-60 | 0.22 |
| 3:74,664,897 G / A | TBC1D1 | 0.007 | 5.4e-7 | -0.40 |
| 22:26,322,324 A / G | LOC111091737 | 0.29 | 5.9-e7 | -0.072 |

**B** 13 : 63,214,897 T / A

Nearest gene: ALB

MAF ranges from 0.33 to 0.37

View on CanFam4 UCSC

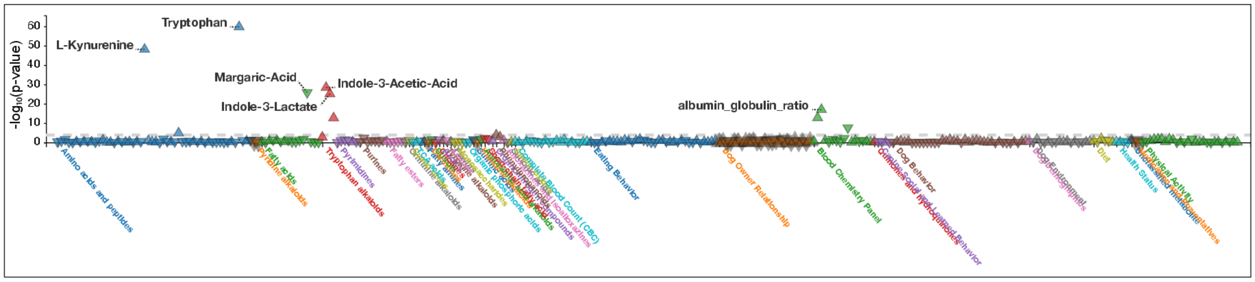

| Category | Phenotype | P-value | Effect size | Number of samples |
|---|---|---|---|---|
| Amino acids and peptides | Tryptophan | 1.8e-60 | 0.22 | 924 |
| Amino acids and peptides | L-Kynurenine | 8.0e-49 | 0.25 | 924 |
| Tryptophan alkaloids | Indole-3-Acetic-Acid | 4.1e-29 | 0.27 | 924 |

**Fig. S5. Dog Aging Project PheWeb for interactive browsing of GWAS results. (A)** GWAS results for tryptophan displayed in PheWeb, including a Manhattan plot, quantile–quantile plot, and a table of the top tryptophan-associated SNPs. **(B)** PheWAS results for the top tryptophan-associated SNP, shown as a Manhattan plot of associations across all other phenotypes and a table summarizing its strongest phenotype associations.

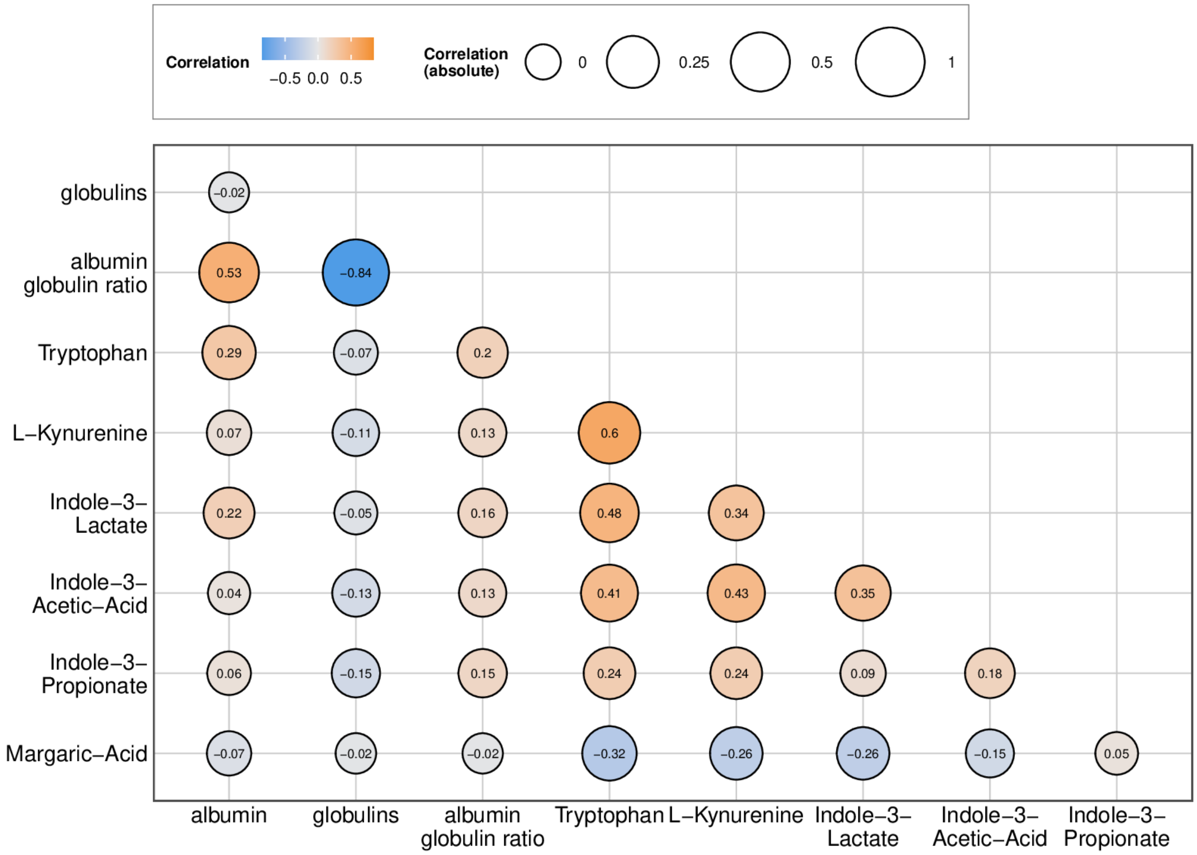

**Fig. S6. Phenotypic correlations across traits with genome-wide significant association at ALB locus.** We calculated phenotype correlation coefficients across all blood traits that had the ALB locus as genome-wide significant in our association studies. Pearson's correlation coefficients are displayed in the circles for pairwise comparison between traits. Shades of blue refer to negative phenotypic correlations while shades of red refer to positive phenotypic correlations.

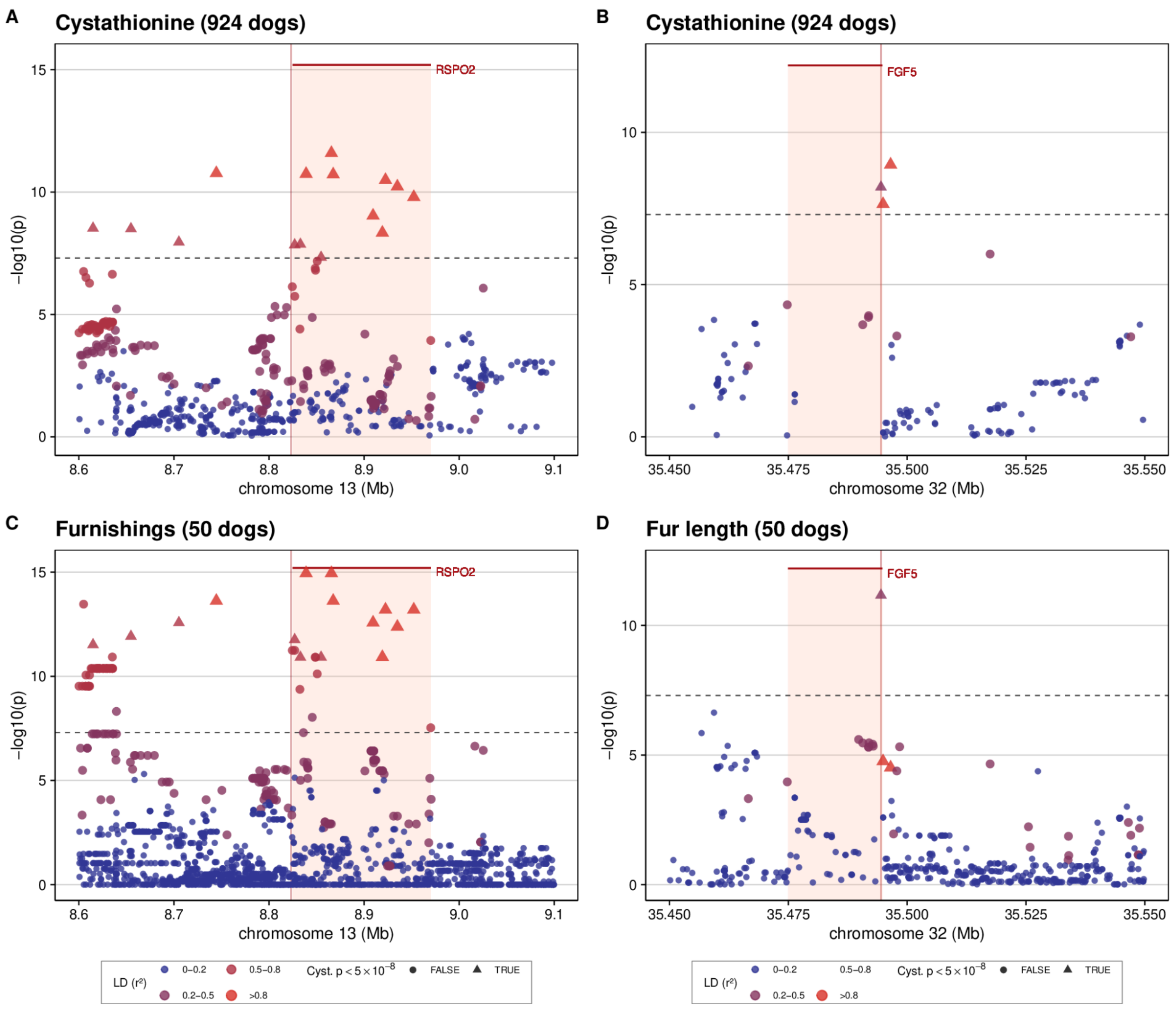

**Fig. S7. Cystathionine and fur type regional association plots at the RSPO2 and FGF5 locus.** SNPs in each region are colored according to their linkage disequilibrium ($r^2$) with the top cystathionine associated variant. Gene locations are light red boxes topped with horizontal red lines with labels, and the location of previously reported variants shown as vertical red lines(*49*). The horizontal dashed line is the genomewide significance threshold of 5e-8. (A) Cystathionine GWAS results for the *RSPO2* locus. (B) Cystathionine GWAS results for the *FGF5* locus. (C) Furnishings GWAS results for the *RSPO2* locus. (D) Fur length GWAS results for the *FGF5* locus.

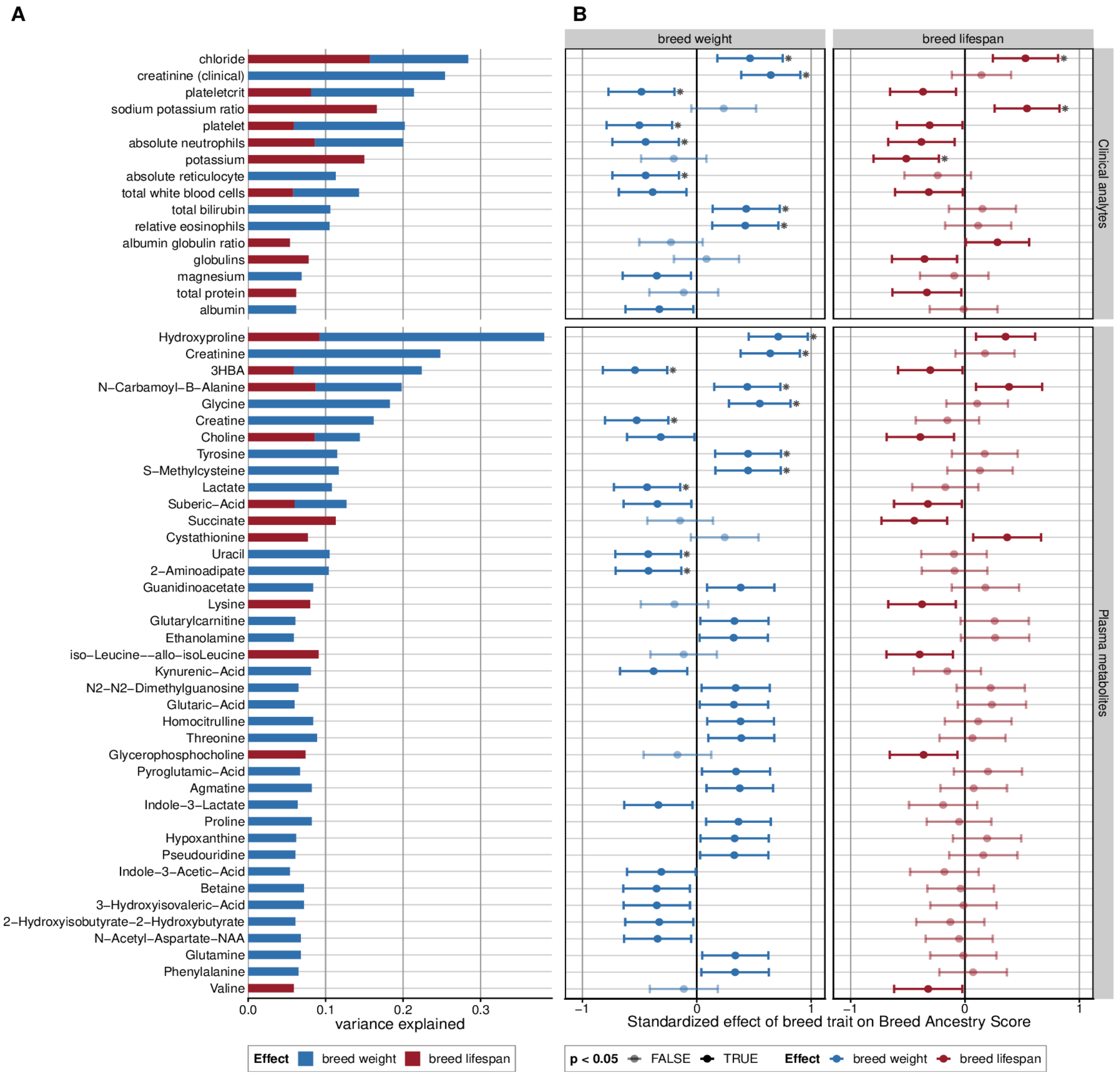

**Fig. S8. Effect of breed characteristics such as lifespan and weight on Breed Ancestry Scores of blood traits. (A)** Stacked bars show variance explained by breed weight and breed lifespan on Breed Ancestry Score, expressed as generalized eta-squared (ges), from the linear model (REML.t.val ~ breed.weight.kg + lifespan) **(B)** Effect estimates for the traits in **(A)**, shown as $\beta \pm 1.96$ s.e. from the model above, faceted by covariate; asterisks indicate $p \leq 0.05$.

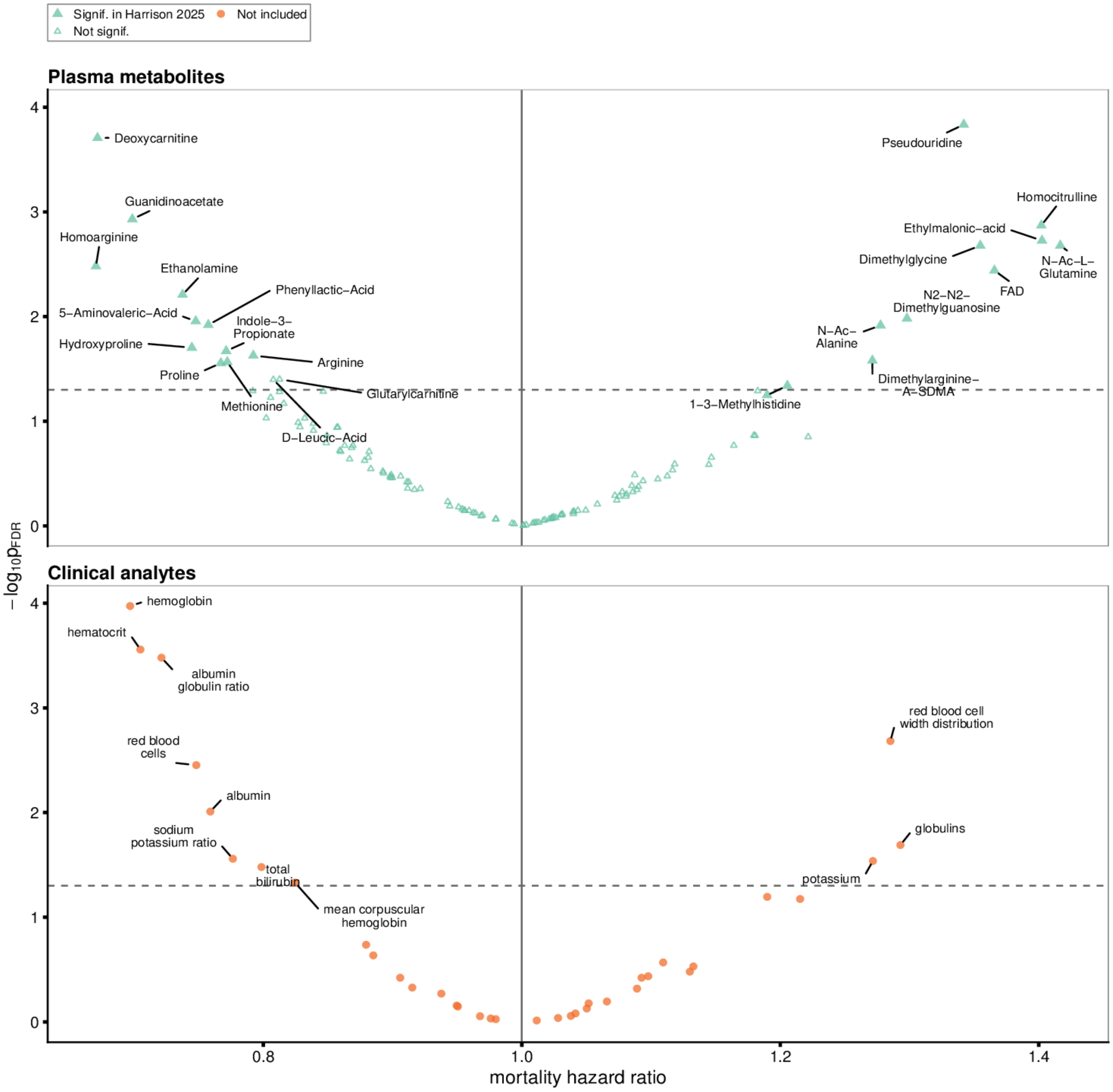

**Fig. S9. Cox regression hazard ratios of blood traits.** Each point is a blood phenotype, with Cox proportional hazard ratio in individual dogs (x) and its corresponding -$\log_{10}$(FDR-adjusted p-value) (y). Vertical dashed line at hazard ratio = 1 implies no difference in risk of death. Points to the right of this vertical dashed line are phenotypes that increase risk of death in individual dogs, while points to the left are phenotypes that decrease risk of death. Filled green triangles are dog metabolite biomarkers of mortality identified by Harrison et al.(*50*) using the same longitudinal data from the Dog Aging Project.

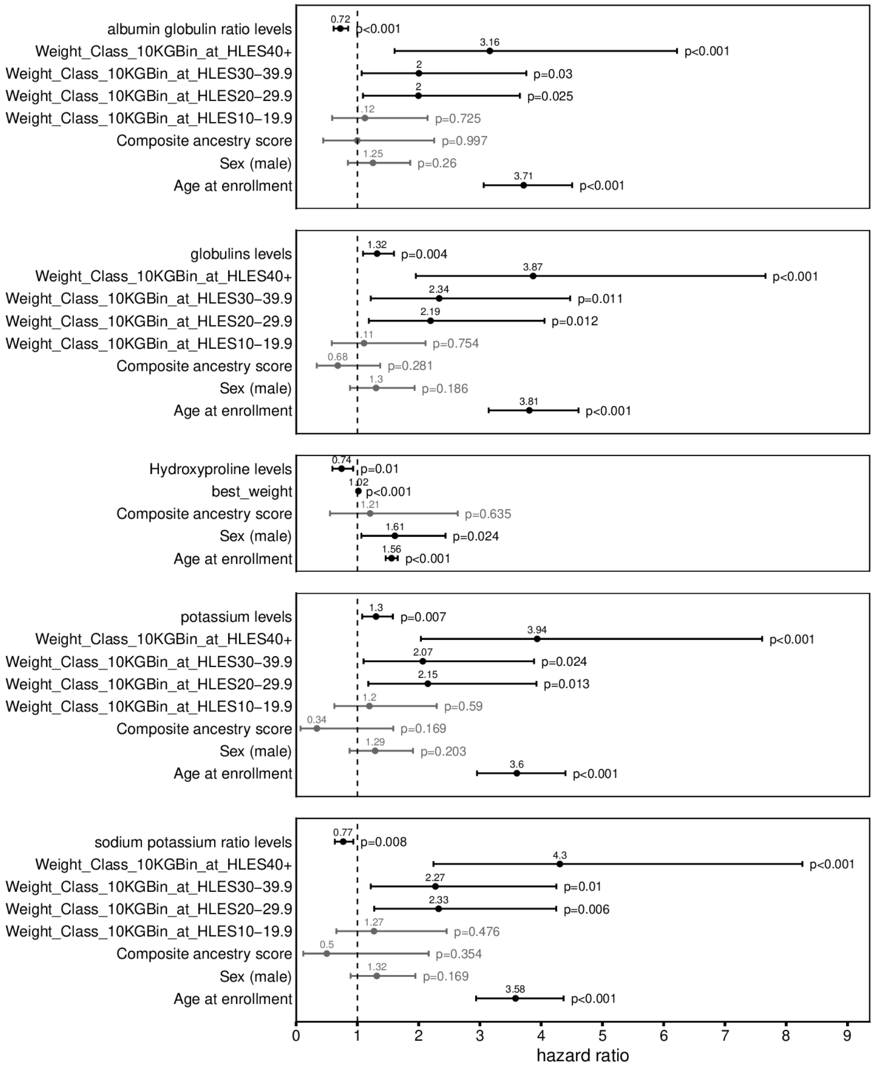

**Fig. S10. Cox regression analysis of protective and risk-associated blood traits.** The effect of composite ancestry, being male, enrollment age, body weight, and blood trait levels on mortality risk in individual dogs. Vertical dashed line at hazard ratio = 1 implies no difference in risk of death. Hazard ratio value shown above each point and the confidence interval shown for each effect. Corresponding p-value for each hazard ratio is shown to the right of the upper bound of the confidence interval. Composite ancestry score is calculated as a sum of proportion ancestry (Breed Ancestry Scores) from the top 9 breeds impacting the levels of each respective trait according to our LMER results.

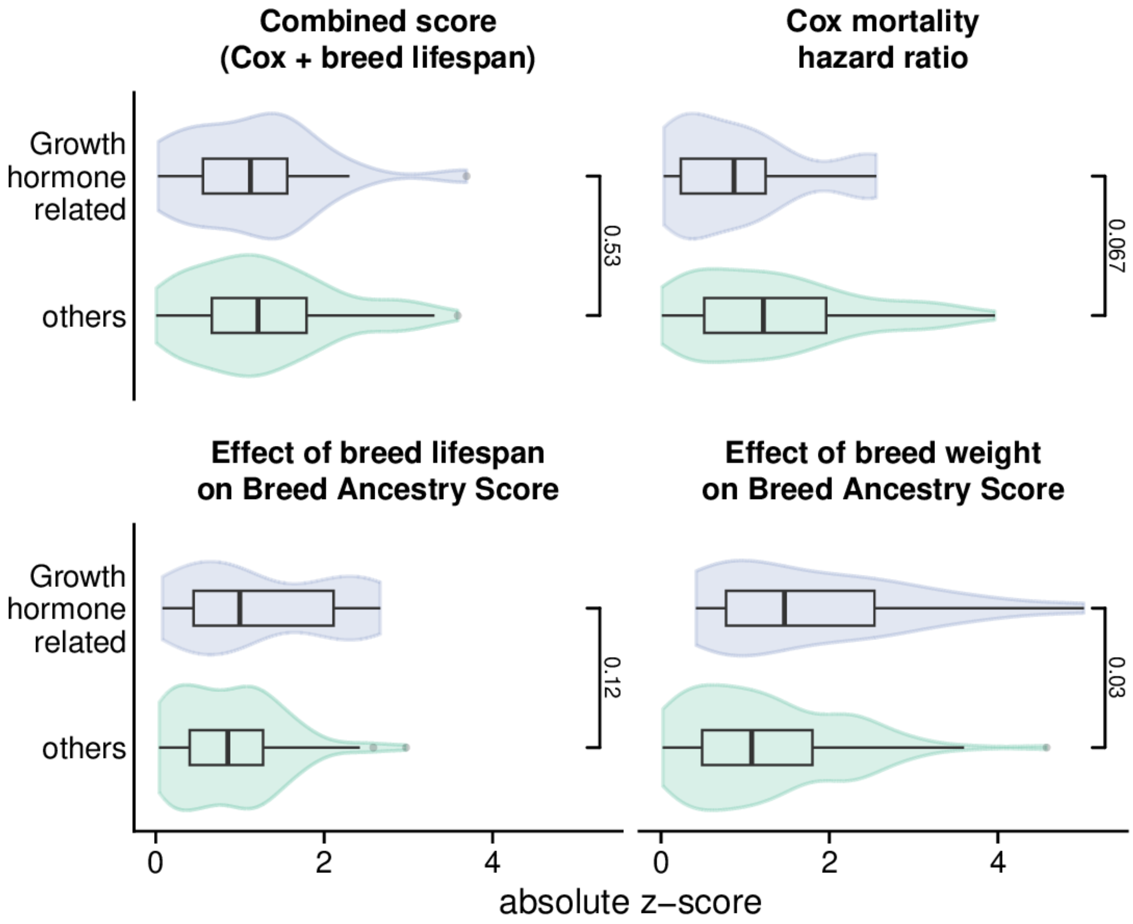

**Fig. S11. Growth hormone related metabolites are similar to other metabolites.** Plasma metabolites linked to the GH/IGF1 pathway (blue) do not differ from other metabolites in their combined z scores, Cox mortality hazard z scores, or breed lifespan effect z scores. However, breed body weight z scores are higher, suggesting that Breed Ancestry Scores for GH-linked metabolites are more explained by breed body size compared to other metabolites.

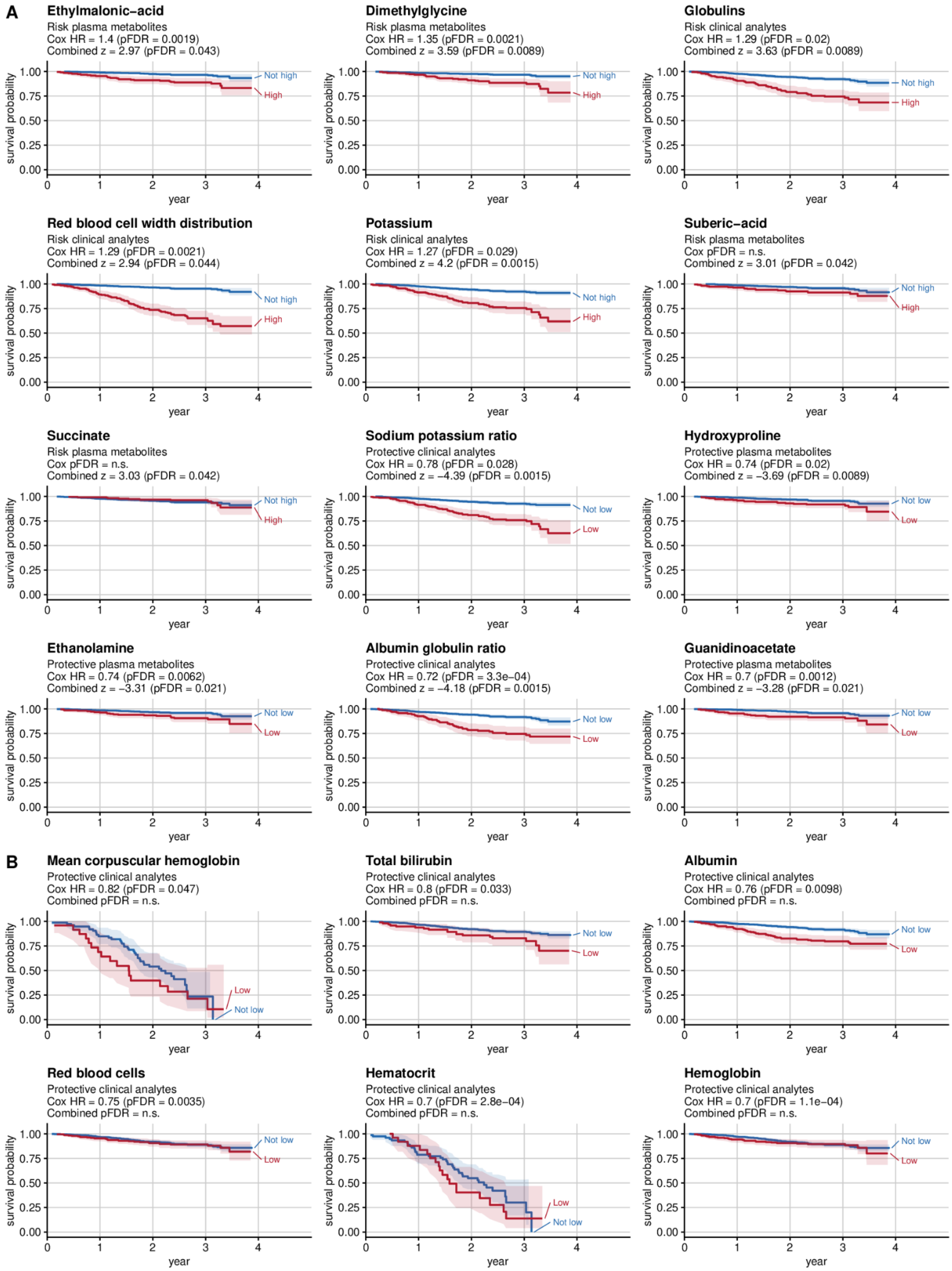

**Fig. S12. Kaplan-Meier survival curves for blood traits.** Kaplan-Meier curves for **(A)** all blood traits with $p_{FDR}<0.05$ in the combined analysis and **(B)** any other clinical analytes with $p_{FDR}<0.05$ in the Cox survival analysis (included because these were not part of an earlier publication analyzing only plasma metabolites (*50*). Time on the x-axis refers to follow-up time in years where time = 0 is the initial blood draw. Each phenotype is stratified by a combination of metabolite levels and broad age groups. Curves are colored according to risk of death with red implying the greatest risk, followed by blue for intermediate risk, then purple for low risk, and black for the least risk of death in individual dogs. For stratification labels, "Low" is defined as having metabolite values below or equal to the 25th percentile (Q1), while "not low" is defined as having metabolite levels greater than the 25th percentile. "High" includes dogs with metabolite values greater than the 75th percentile (Q3), and "not high" includes dogs with metabolite values less than or equal to the 75th percentile. Old dogs are defined as dogs greater than 7 years old at enrollment, whereas not old refers to dogs less than or equal to 7 years old.

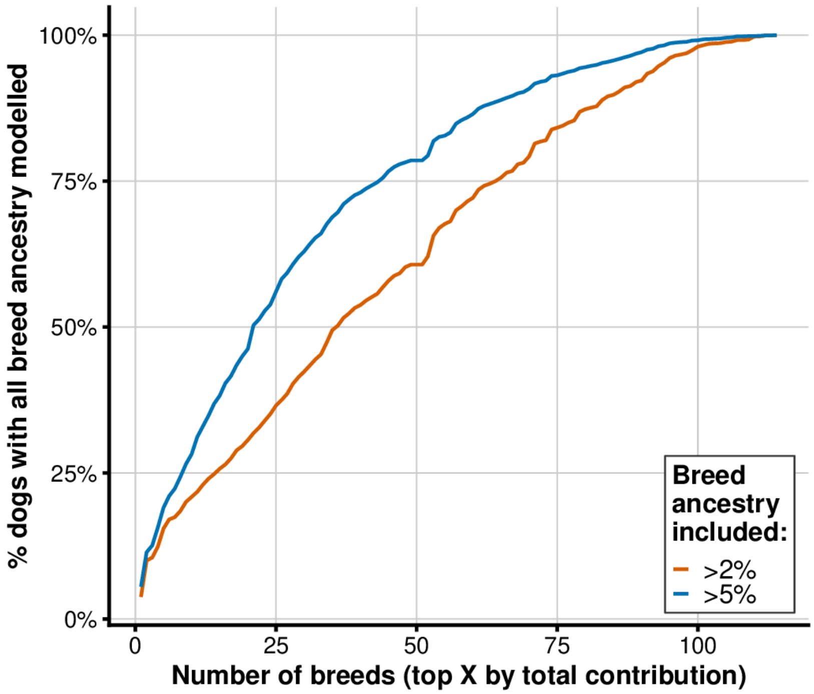

**Fig. S13. Number of breeds needed to model ancestry of Dog Aging Project dog population.** Number of breeds required to model all breeds contributing more than 2% (orange) or 5% (blue) of the ancestry found in any individual dog.

**Table S1.**

Correlation of body weight predicted by breed with measured body weight

| set | term | beta | se | statistic | p | r2 | n |
|---|---|---|---|---|---|---|---|
| Complex mixed-breed dogs (age > 2 years) | (Intercept) | -5.21 | 1.66 | -3.13604069 | 1.75e-03 | 0.67 | 1229 |
| Complex mixed-breed dogs (age > 2 years) | pred_weight | 1.11 | 0.02 | 49.50678417 | 1.39e-294 | 0.67 | 1229 |
| Complex mixed-breed dogs (age > 2 years) | sexmale | 3.19 | 0.35 | 9.157566439 | 2.17e-19 | 0.67 | 1229 |
| Complex mixed-breed dogs (age > 2 years) | Sterilization statussterilized | 3.4 | 1.58 | 2.145463201 | 3.21e-02 | 0.67 | 1229 |
| Mixed-breed dogs (age > 2 years) | (Intercept) | -2.63 | 0.88 | -2.994265223 | 2.78e-03 | 0.75 | 2451 |
| Mixed-breed dogs (age > 2 years) | pred_weight | 1.04 | 0.01 | 86.05283442 | 0.00e+00 | 0.75 | 2451 |
| Mixed-breed dogs (age > 2 years) | sexmale | 3.37 | 0.23 | 14.34311502 | 7.36e-45 | 0.75 | 2451 |
| Mixed-breed dogs (age > 2 years) | Sterilization statussterilized | 2.26 | 0.84 | 2.672273044 | 7.58e-03 | 0.75 | 2451 |
| Single-breed dogs (age > 2 years) | (Intercept) | -1.23 | 0.44 | -2.822254377 | 4.81e-03 | 0.84 | 2475 |
| Single-breed dogs (age > 2 years) | pred_weight | 1.03 | 0.01 | 112.8059364 | 0.00e+00 | 0.84 | 2475 |
| Single-breed dogs (age > 2 years) | sexmale | 3.89 | 0.25 | 15.83827342 | 6.93e-54 | 0.84 | 2475 |
| Single-breed dogs (age > 2 years) | Sterilization statussterilized | 0.54 | 0.34 | 1.598768086 | 1.10e-01 | 0.84 | 2475 |

**Table S2.**

Phenotypes in genetic analysis of the Dog Aging Project

| phenotype | plot_label | paper_phenotype_category | pheweb_phenotype_category | pheweb_phenotype_name | sample_size |
|---|---|---|---|---|---|
| krt_cbc_abs_eosinophils_sqrt_transformed | absolute eosinophils | Clinical analytes | Complete Blood Count (CBC) | absolute_eosinophils | 918 |
| krt_cbc_abs_lymphocytes_sqrt_transformed | absolute lymphocytes | Clinical analytes | Complete Blood Count (CBC) | absolute_lymphocytes | 918 |
| krt_cbc_abs_monocytes_sqrt_transformed | absolute monocytes | Clinical analytes | Complete Blood Count (CBC) | absolute_monocytes | 918 |
| krt_cbc_abs_neutrophils_sqrt_transformed | absolute neutrophils | Clinical analytes | Complete Blood Count (CBC) | absolute_neutrophils | 918 |
| krt_cbc_hct_sqrt_transformed | hematocrit | Clinical analytes | Complete Blood Count (CBC) | hematocrit | 923 |
| krt_cbc_hgb_sqrt_transformed | hemoglobin | Clinical analytes | Complete Blood Count (CBC) | hemoglobin | 923 |
| krt_cbc_mch_sqrt_transformed | mean corpuscular hemoglobin | Clinical analytes | Complete Blood Count (CBC) | mean_corpuscular_hemoglobin | 923 |
| krt_cbc_mchc_sqrt_transformed | mean corpuscular hemoglobin concentration | Clinical analytes | Complete Blood Count (CBC) | mean_corpuscular_hemoglobin_concentration | 923 |
| krt_cbc_mcv_sqrt_transformed | mean corpuscular volume | Clinical analytes | Complete Blood Count (CBC) | mean_corpuscular_volume | 923 |
| krt_cbc_mpv_sqrt_transformed | mean platelet volume | Clinical analytes | Complete Blood Count (CBC) | mean_platelet_volume | 919 |
| krt_cbc_pct_sqrt_transformed | plateletcrit | Clinical analytes | Complete Blood Count (CBC) | plateletcrit | 919 |
| krt_cbc_plt_sqrt_transformed | platelet | Clinical analytes | Complete Blood Count (CBC) | platelet | 715 |
| krt_cbc_rbc_sqrt_transformed | red blood cells | Clinical analytes | Complete Blood Count (CBC) | red_blood_cells | 923 |
| krt_cbc_rdw_sqrt_transformed | red blood cell width distribution | Clinical analytes | Complete Blood Count (CBC) | red_blood_cell_width_distribution | 923 |
| krt_cbc_rel_eosinophils_sqrt_transformed | relative eosinophils | Clinical analytes | Complete Blood Count (CBC) | relative_eosinophils | 918 |
| krt_cbc_rel_lymphocytes_sqrt_transformed | relative lymphocytes | Clinical analytes | Complete Blood Count (CBC) | relative_lymphocytes | 918 |
| krt_cbc_rel_monocytes_sqrt_transformed | relative monocytes | Clinical analytes | Complete Blood Count (CBC) | relative_monocytes | 918 |
| krt_cbc_rel_neutrophils_sqrt_transformed | relative neutrophils | Clinical analytes | Complete Blood Count (CBC) | relative_neutrophils | 918 |
| krt_cbc_retic_abs_sqrt_transformed | absolute reticulocyte | Clinical analytes | Complete Blood Count (CBC) | absolute_reticulocyte | 922 |
| krt_cbc_total_wbcs_sqrt_transformed | total white blood cells | Clinical analytes | Complete Blood Count (CBC) | total_white_blood_cells | 923 |
| krt_cp_alb_glob_ratio_value_ln_transformed | albumin globulin ratio | Clinical analytes | Blood Chemistry Panel | albumin_globulin_ratio | 953 |
| krt_cp_albumin_value_ln_transformed | albumin | Clinical analytes | Blood Chemistry Panel | albumin | 953 |
| krt_cp_alkp_value_ln_transformed | alkaline phosphatase | Clinical analytes | Blood Chemistry Panel | alkaline_phosphatase | 953 |
| krt_cp_alt_value_ln_transformed | alanine aminotransferase | Clinical analytes | Blood Chemistry Panel | alanine_aminotransferase | 953 |
| krt_cp_amylase_value_ln_transformed | amylase | Clinical analytes | Blood Chemistry Panel | amylase | 953 |

| phenotype | plot_label | paper_phenotype_category | pheweb_phenotype_category | pheweb_phenotype_name | sample_size |
|---|---|---|---|---|---|
| krt_cp_bilirubin_total_value_ln_transformed | total bilirubin | Clinical analytes | Blood Chemistry Panel | total_bilirubin | 953 |
| krt_cp_bun_value_ln_transformed | blood urea nitrogen | Clinical analytes | Blood Chemistry Panel | blood_urea_nitrogen | 953 |
| krt_cp_chloride_value_ln_transformed | chloride | Clinical analytes | Blood Chemistry Panel | chloride | 953 |
| krt_cp_cholesterol_value_ln_transformed | cholesterol | Clinical analytes | Blood Chemistry Panel | cholesterol | 953 |
| krt_cp_creatinine_value_ln_transformed | creatinine (clinical) | Clinical analytes | Blood Chemistry Panel | creatinine | 953 |
| krt_cp_globulins_value_ln_transformed | globulins | Clinical analytes | Blood Chemistry Panel | globulins | 953 |
| krt_cp_magnesium_value_ln_transformed | magnesium | Clinical analytes | Blood Chemistry Panel | magnesium | 953 |
| krt_cp_potassium_value_ln_transformed | potassium | Clinical analytes | Blood Chemistry Panel | potassium | 953 |
| krt_cp_sodium_value_ln_transformed | sodium | Clinical analytes | Blood Chemistry Panel | sodium | 953 |
| krt_cp_sp_ratio_value_ln_transformed | sodium potassium ratio | Clinical analytes | Blood Chemistry Panel | sodium_potassium_ratio | 953 |
| krt_cp_total_protein_value_ln_transformed | total protein | Clinical analytes | Blood Chemistry Panel | total_protein | 953 |
| 1-3-Methylhistidine | 1-3-Methylhistidine | Plasma metabolites | Amino acids and peptides | 1-3-Methylhistidine | 924 |
| 1-Methylnicotinamide | 1-Methylnicotinamide | Plasma metabolites | Pyridine alkaloids | 1-Methylnicotinamide | 924 |
| 2-Aminoadipate | 2-Aminoadipate | Plasma metabolites | Amino acids and peptides | 2-Aminoadipate | 924 |
| 2-Hydroxyglutarate | 2-Hydroxyglutarate | Plasma metabolites | Fatty acids | 2-Hydroxyglutarate | 924 |
| 2-Hydroxyisobutyrate-2-Hydroxybutyrate | 2-Hydroxyisobutyrate-2-Hydroxybutyrate | Plasma metabolites | Fatty acids | 2-Hydroxyisobutyrate-2-Hydroxybutyrate | 924 |
| 3-Hydroxyisovaleric-Acid | 3-Hydroxyisovaleric-Acid | Plasma metabolites | Fatty acids | 3-Hydroxyisovaleric-Acid | 924 |
| 3-Indoxyl-Sulfate | 3-Indoxyl-Sulfate | Plasma metabolites | Tryptophan alkaloids | 3-Indoxyl-Sulfate | 924 |
| 3-Methyl-3-Hydroxyglutaric-Acid | 3-Methyl-3-Hydroxyglutaric-Acid | Plasma metabolites | Fatty acids | 3-Methyl-3-Hydroxyglutaric-Acid | 924 |
| 3HBA | 3HBA | Plasma metabolites | Fatty acids | 3HBA | 924 |
| 5-Aminovaleric-Acid | 5-Aminovaleric-Acid | Plasma metabolites | Fatty acids | 5-Aminovaleric-Acid | 924 |
| 5-Methylcytidine | 5-Methylcytidine | Plasma metabolites | Pyrimidines | 5-Methylcytidine | 924 |
| 7-Methylguanine | 7-Methylguanine | Plasma metabolites | Purines | 7-Methylguanine | 924 |

| phenotype | plot_label | paper_phenotype_category | pheweb_phenotype_category | pheweb_phenotype_name | sample_size |
|---|---|---|---|---|---|
| Acetylcarnitine | Acetylcarnitine | Plasma metabolites | Fatty esters | Acetylcarnitine | 924 |
| Adenosine | Adenosine | Plasma metabolites | Purines | Adenosine | 924 |
| Adenylosuccinate | Adenylosuccinate | Plasma metabolites | Purines | Adenylosuccinate | 924 |
| Agmatine | Agmatine | Plasma metabolites | Ornithine alkaloids | Agmatine | 924 |
| Alanine | Alanine | Plasma metabolites | Amino acids and peptides | Alanine | 924 |
| Allantoin | Allantoin | Plasma metabolites | Cyclic ureas | Allantoin | 924 |
| Alpha-Ketoglutaric-Acid | Alpha-Ketoglutaric-Acid | Plasma metabolites | TCA acids | Alpha-Ketoglutaric-Acid | 924 |
| Anserine | Anserine | Plasma metabolites | Amino acids and peptides | Anserine | 924 |
| Arachidonate | Arachidonate | Plasma metabolites | Fatty acids | Arachidonate | 924 |
| Arginine | Arginine | Plasma metabolites | Amino acids and peptides | Arginine | 924 |
| Asparagine | Asparagine | Plasma metabolites | Amino acids and peptides | Asparagine | 924 |
| Aspartic-Acid | Aspartic-Acid | Plasma metabolites | Amino acids and peptides | Aspartic-Acid | 924 |
| Betaine | Betaine | Plasma metabolites | Amino acids and peptides | Betaine | 924 |
| Cadaverine | Cadaverine | Plasma metabolites | Fatty amines | Cadaverine | 924 |
| Carnitine | Carnitine | Plasma metabolites | Carnitines | Carnitine | 924 |
| Carnosine | Carnosine | Plasma metabolites | Amino acids and peptides | Carnosine | 924 |
| Cholesteryl-Sulfate | Cholesteryl-Sulfate | Plasma metabolites | Sterols | Cholesteryl-Sulfate | 924 |
| Choline | Choline | Plasma metabolites | Cholines | Choline | 924 |
| Citrulline | Citrulline | Plasma metabolites | Amino acids and peptides | Citrulline | 924 |
| Creatine | Creatine | Plasma metabolites | Amino acids and peptides | Creatine | 924 |
| Creatinine | Creatinine | Plasma metabolites | Azoles | Creatinine | 924 |
| Cystathionine | Cystathionine | Plasma metabolites | Amino acids and peptides | Cystathionine | 924 |
| Cystine | Cystine | Plasma metabolites | Amino acids and peptides | Cystine | 924 |
| Cytidine | Cytidine | Plasma metabolites | Pyrimidines | Cytidine | 924 |
| D-Leucic-Acid | D-Leucic-Acid | Plasma metabolites | Fatty acids | D-Leucic-Acid | 924 |

| phenotype | plot_label | paper_phenotype_category | pheweb_phenotype_category | pheweb_phenotype_name | sample_size |
|---|---|---|---|---|---|
| Deoxycarnitine | Deoxycarnitine | Plasma metabolites | Fatty esters | Deoxycarnitine | 924 |
| Dimethylarginine-A-SDMA | Dimethylarginine-A-SDMA | Plasma metabolites | Amino acids and peptides | Dimethylarginine-A-SDMA | 924 |
| Dimethylglycine | Dimethylglycine | Plasma metabolites | Amino acids and peptides | Dimethylglycine | 924 |
| Epinephrine | Epinephrine | Plasma metabolites | Tyrosine alkaloids | Epinephrine | 924 |
| Ethanolamine | Ethanolamine | Plasma metabolites | Amines | Ethanolamine | 924 |
| Ethylmalonic-acid | Ethylmalonic-acid | Plasma metabolites | Fatty acids | Ethylmalonic-acid | 924 |
| FAD | FAD | Plasma metabolites | Flavins | FAD | 924 |
| G6P | G6P | Plasma metabolites | Monosaccharides | G6P | 924 |
| Glucoronate | Glucoronate | Plasma metabolites | Monosaccharides | Glucoronate | 924 |
| Glucose | Glucose | Plasma metabolites | Monosaccharides | Glucose | 924 |
| Glutamic-acid | Glutamic-acid | Plasma metabolites | Amino acids and peptides | Glutamic-acid | 924 |
| Glutamine | Glutamine | Plasma metabolites | Amino acids and peptides | Glutamine | 924 |
| Glutaric-Acid | Glutaric-Acid | Plasma metabolites | Fatty acids | Glutaric-Acid | 924 |
| Glutarylcarnitine | Glutarylcarnitine | Plasma metabolites | Fatty esters | Glutarylcarnitine | 924 |
| Glycerol-3-P | Glycerol-3-P | Plasma metabolites | Organic phosphoric acids | Glycerol-3-P | 924 |
| Glycerophosphocholine | Glycerophosphocholine | Plasma metabolites | Organic phosphoric acids | Glycerophosphocholine | 924 |
| Glycine | Glycine | Plasma metabolites | Amino acids and peptides | Glycine | 924 |
| Guanidinoacetate | Guanidinoacetate | Plasma metabolites | Amino acids and peptides | Guanidinoacetate | 924 |
| Histidine | Histidine | Plasma metabolites | Amino acids and peptides | Histidine | 924 |
| Homoarginine | Homoarginine | Plasma metabolites | Amino acids and peptides | Homoarginine | 924 |
| Homocitrulline | Homocitrulline | Plasma metabolites | Amino acids and peptides | Homocitrulline | 924 |
| Hydroxyproline | Hydroxyproline | Plasma metabolites | Amino acids and peptides | Hydroxyproline | 924 |
| Hypotaurine | Hypotaurine | Plasma metabolites | Sulfinic acids | Hypotaurine | 924 |
| Hypoxanthine | Hypoxanthine | Plasma metabolites | Purines | Hypoxanthine | 924 |
| Imidazole-Propionate | Imidazole-Propionate | Plasma metabolites | Histidine alkaloids | Imidazole-Propionate | 924 |
| Indole-3-Acetic-Acid | Indole-3-Acetic-Acid | Plasma metabolites | Tryptophan alkaloids | Indole-3-Acetic-Acid | 924 |

| phenotype | plot_label | paper_phenotype_category | pheweb_phenotype_category | pheweb_phenotype_name | sample_size |
|---|---|---|---|---|---|
| Indole-3-Lactate | Indole-3-Lactate | Plasma metabolites | Tryptophan alkaloids | Indole-3-Lactate | 924 |
| Indole-3-Propionate | Indole-3-Propionate | Plasma metabolites | Tryptophan alkaloids | Indole-3-Propionate | 924 |
| Kynurenic-Acid | Kynurenic-Acid | Plasma metabolites | Anthranilic acid alkaloids | Kynurenic-Acid | 924 |
| L-Kynurenine | L-Kynurenine | Plasma metabolites | Amino acids and peptides | L-Kynurenine | 924 |
| Lactate | Lactate | Plasma metabolites | Short-chain fatty acid | Lactate | 924 |
| Leucine--D-Norleucine | Leucine--D-Norleucine | Plasma metabolites | Amino acids and peptides | Leucine--D-Norleucine | 924 |
| Linoleic-Acid | Linoleic-Acid | Plasma metabolites | Fatty acids | Linoleic-Acid | 924 |
| Linolenic-Acid | Linolenic-Acid | Plasma metabolites | Fatty acids | Linolenic-Acid | 924 |
| Lysine | Lysine | Plasma metabolites | Amino acids and peptides | Lysine | 924 |
| Margaric-Acid | Margaric-Acid | Plasma metabolites | Fatty acids | Margaric-Acid | 924 |
| Methionine | Methionine | Plasma metabolites | Amino acids and peptides | Methionine | 924 |
| Methionine-Sulfoxide | Methionine-Sulfoxide | Plasma metabolites | Amino acids and peptides | Methionine-Sulfoxide | 924 |
| N-Ac-Alanine | N-Ac-Alanine | Plasma metabolites | Amino acids and peptides | N-Ac-Alanine | 924 |
| N-Ac-Glutamate | N-Ac-Glutamate | Plasma metabolites | Amino acids and peptides | N-Ac-Glutamate | 924 |
| N-Ac-L-Glutamine | N-Ac-L-Glutamine | Plasma metabolites | Amino acids and peptides | N-Ac-L-Glutamine | 924 |
| N-Ac-Phenylalanine | N-Ac-Phenylalanine | Plasma metabolites | Amino acids and peptides | N-Ac-Phenylalanine | 924 |
| N-Ac-Tryptophan | N-Ac-Tryptophan | Plasma metabolites | Amino acids and peptides | N-Ac-Tryptophan | 924 |
| N-Acetyl-Aspartate-NAA | N-Acetyl-Aspartate-NAA | Plasma metabolites | Amino acids and peptides | N-Acetyl-Aspartate-NAA | 924 |
| N-Carbamoyl-B-Alanine | N-Carbamoyl-B-Alanine | Plasma metabolites | Organonitrogen compounds | N-Carbamoyl-B-Alanine | 924 |
| N2-N2-Dimethylguanosine | N2-N2-Dimethylguanosine | Plasma metabolites | Purines | N2-N2-Dimethylguanosine | 924 |
| N6-Acetyl-Lysine | N6-Acetyl-Lysine | Plasma metabolites | Amino acids and peptides | N6-Acetyl-Lysine | 924 |
| N6-Trimethyllysine | N6-Trimethyllysine | Plasma metabolites | Amino acids and peptides | N6-Trimethyllysine | 924 |
| Ornithine | Ornithine | Plasma metabolites | Amino acids and peptides | Ornithine | 924 |
| Orotate | Orotate | Plasma metabolites | Pyrimidines | Orotate | 924 |
| Oxalacetate | Oxalacetate | Plasma metabolites | TCA acids | Oxalacetate | 924 |

| phenotype | plot_label | paper_phenotype_category | pheweb_phenotype_category | pheweb_phenotype_name | sample_size |
|---|---|---|---|---|---|
| Oxidized-Glutathione | Oxidized-Glutathione | Plasma metabolites | Amino acids and peptides | Oxidized-Glutathione | 924 |
| PPA | PPA | Plasma metabolites | Phenylpropanoids | PPA | 924 |
| Palmitic-Acid | Palmitic-Acid | Plasma metabolites | Fatty acids | Palmitic-Acid | 924 |
| Pantothenate | Pantothenate | Plasma metabolites | Amino acids and peptides | Pantothenate | 924 |
| Phenylacetylglutamine | Phenylacetylglutamine | Plasma metabolites | Amino acids and peptides | Phenylacetylglutamine | 924 |
| Phenylalanine | Phenylalanine | Plasma metabolites | Amino acids and peptides | Phenylalanine | 924 |
| Phenyllactic-Acid | Phenyllactic-Acid | Plasma metabolites | Phenylpropanoids | Phenyllactic-Acid | 924 |
| Proline | Proline | Plasma metabolites | Amino acids and peptides | Proline | 924 |
| Pseudouridine | Pseudouridine | Plasma metabolites | Pyrimidines | Pseudouridine | 924 |
| Pyroglutamic-Acid | Pyroglutamic-Acid | Plasma metabolites | Amino acids and peptides | Pyroglutamic-Acid | 924 |
| Pyruvate | Pyruvate | Plasma metabolites | Short-chain fatty acid | Pyruvate | 924 |
| Reduced-Glutathione | Reduced-Glutathione | Plasma metabolites | Amino acids and peptides | Reduced-Glutathione | 924 |
| Riboflavin | Riboflavin | Plasma metabolites | Alloxazines and isoalloxazines | Riboflavin | 924 |
| S-Methylcysteine | S-Methylcysteine | Plasma metabolites | Amino acids and peptides | S-Methylcysteine | 924 |
| SAH | SAH | Plasma metabolites | Purines | SAH | 924 |
| Sarcosine | Sarcosine | Plasma metabolites | Amino acids and peptides | Sarcosine | 924 |
| Serine | Serine | Plasma metabolites | Amino acids and peptides | Serine | 924 |
| Suberic-Acid | Suberic-Acid | Plasma metabolites | Fatty acids | Suberic-Acid | 924 |
| Succinate | Succinate | Plasma metabolites | TCA acids | Succinate | 924 |
| Succinylcarnitine | Succinylcarnitine | Plasma metabolites | Fatty esters | Succinylcarnitine | 924 |
| Taurine | Taurine | Plasma metabolites | Sulfonic acids | Taurine | 924 |
| Thiamine | Thiamine | Plasma metabolites | Pyrimidines | Thiamine | 924 |
| Threonine | Threonine | Plasma metabolites | Amino acids and peptides | Threonine | 924 |
| Trigonelline | Trigonelline | Plasma metabolites | Pyridine alkaloids | Trigonelline | 924 |
| Trimethylamine-N-Oxide-TMAO | Trimethylamine-N-Oxide-TMAO | Plasma metabolites | Amine oxides | Trimethylamine-N-Oxide-TMAO | 924 |
| Tryptophan | Tryptophan | Plasma metabolites | Amino acids and peptides | Tryptophan | 924 |

| phenotype | plot_label | paper_phenotype_category | pheweb_phenotype_category | pheweb_phenotype_name | sample_size |
|---|---|---|---|---|---|
| Tyrosine | Tyrosine | Plasma metabolites | Amino acids and peptides | Tyrosine | 924 |
| Uracil | Uracil | Plasma metabolites | Pyrimidines | Uracil | 924 |
| Valine | Valine | Plasma metabolites | Amino acids and peptides | Valine | 924 |
| Xanthine | Xanthine | Plasma metabolites | Purines | Xanthine | 924 |
| alpha-Tocopherol | alpha-Tocopherol | Plasma metabolites | Quinones and hydroquinones | alpha-Tocopherol | 924 |
| gamma-Aminobutyrate | gamma-Aminobutyrate | Plasma metabolites | Fatty acids | gamma-Aminobutyrate | 924 |
| iso-Leucine--allo-isoLeucine | iso-Leucine--allo-isoLeucine | Plasma metabolites | Unclassified metabolite | iso-Leucine--allo-isoLeucine | 924 |
| isoValerylcarnitine | isoValerylcarnitine | Plasma metabolites | Fatty esters | isoValerylcarnitine | 924 |
| n-Formylmethionine | n-Formylmethionine | Plasma metabolites | Amino acids and peptides | n-Formylmethionine | 924 |
| afus_dora_10_runs_around_alot | Runs around a lot | Survey question | Eating Behavior | afus_dora_10_runs_around_alot | 4789 |
| afus_dora_11_interested_eating_after_meal | After a meal my dog is still interested in eating | Survey question | Eating Behavior | afus_dora_11_interested_eating_after_meal | 4789 |
| afus_dora_12_slow_eater | Takes their time to eat | Survey question | Eating Behavior | afus_dora_12_slow_eater | 4789 |
| afus_dora_13_eats_treats_quickly | Eats treats immediately | Survey question | Eating Behavior | afus_dora_13_eats_treats_quickly | 4789 |
| afus_dora_14_human_food_during_meals | Gets bits of human food when we are eating | Survey question | Eating Behavior | afus_dora_14_human_food_during_meals | 4789 |
| afus_dora_15_eat_anything | Would eat anything | Survey question | Eating Behavior | afus_dora_15_eat_anything | 4789 |
| afus_dora_16_very_fit | Is very fit | Survey question | Eating Behavior | afus_dora_16_very_fit | 4789 |
| afus_dora_17_human_food_often | Often gets human food | Survey question | Eating Behavior | afus_dora_17_human_food_often | 4789 |
| afus_dora_18_upset_stomach_some_foods | Gets an upset tummy on some foods | Survey question | Eating Behavior | afus_dora_18_upset_stomach_some_foods | 4789 |
| afus_dora_19_should_lose_weight | My dog could do with losing weight | Survey question | Eating Behavior | afus_dora_19_should_lose_weight | 4789 |
| afus_dora_1_excited_food | Gets excited when there is food around | Survey question | Eating Behavior | afus_dora_1_excited_food | 4789 |
| afus_dora_20_most_walks_on_leash | Walks are mostly on leash | Survey question | Eating Behavior | afus_dora_20_most_walks_on_leash | 4789 |
| afus_dora_22_diet_to_control_weight | I alter dog's food to control weight | Survey question | Eating Behavior | afus_dora_22_diet_to_control_weight | 4789 |

| phenotype | plot_label | paper_phenotype_category | pheweb_phenotype_category | pheweb_phenotype_name | sample_size |
|---|---|---|---|---|---|
| afus_dora_23_hungry_all_time | Seems to be hungry all the time | Survey question | Eating Behavior | afus_dora_23_hungry_all_time | 4789 |
| afus_dora_24_walks_high_energy | Walks involve a lot of energetic play | Survey question | Eating Behavior | afus_dora_24_walks_high_energy | 4789 |
| afus_dora_25_careful_dogs_weight | Careful about my dog's weight | Survey question | Eating Behavior | afus_dora_25_careful_dogs_weight | 4789 |
| afus_dora_26_sensitive_stomach | Has a sensitive stomach | Survey question | Eating Behavior | afus_dora_26_sensitive_stomach | 4789 |
| afus_dora_27_very_greedy | Is very greedy | Survey question | Eating Behavior | afus_dora_27_very_greedy | 4789 |
| afus_dora_28_vet_often_for_health_problems | Regularly sees vet for health problems | Survey question | Eating Behavior | afus_dora_28_vet_often_for_health_problems | 4789 |
| afus_dora_29_happy_dogs_weight | Happy with my dog's weight | Survey question | Eating Behavior | afus_dora_29_happy_dogs_weight | 4789 |
| afus_dora_2_most_walks_off_leash | Spends most of their walks off leash | Survey question | Eating Behavior | afus_dora_2_most_walks_off_leash | 4789 |
| afus_dora_30_measure_dogs_food | I measure dog's food | Survey question | Eating Behavior | afus_dora_30_measure_dogs_food | 4789 |
| afus_dora_31_regulate_exercise_for_weight | Careful to regulate exercise to keep them slim | Survey question | Eating Behavior | afus_dora_31_regulate_exercise_for_weight | 4789 |
| afus_dora_32_gets_lots_of_exercise | Gets a lot of exercise | Survey question | Eating Behavior | afus_dora_32_gets_lots_of_exercise | 4789 |
| afus_dora_33_upset_stomach_often | Often gets an upset tummy | Survey question | Eating Behavior | afus_dora_33_upset_stomach_often | 4789 |
| afus_dora_34_no_human_food_during_meals | Gets no food at human mealtimes | Survey question | Eating Behavior | afus_dora_34_no_human_food_during_meals | 4789 |
| afus_dora_35_eat_non_food_objects | Would eat non-food objects | Survey question | Eating Behavior | afus_dora_35_eat_non_food_objects | 4789 |
| afus_dora_3_human_leftovers_in_bowl | Gets human leftovers in their food bowl | Survey question | Eating Behavior | afus_dora_3_human_leftovers_in_bowl | 4789 |
| afus_dora_4_waits_for_scraps | Hangs around for scraps even if not getting them | Survey question | Eating Behavior | afus_dora_4_waits_for_scraps | 4789 |
| afus_dora_5_choosy_treats | Is choosy about treats | Survey question | Eating Behavior | afus_dora_5_choosy_treats | 4789 |
| afus_dora_6_waits_during_food_prep | Hangs around when i am preparing human food | Survey question | Eating Behavior | afus_dora_6_waits_during_food_prep | 4789 |
| afus_dora_7_turns_down_food | Turns down food if not hungry | Survey question | Eating Behavior | afus_dora_7_turns_down_food | 4789 |
| afus_dora_8_finishes_meal_quickly | Finishes meal straight away | Survey question | Eating Behavior | afus_dora_8_finishes_meal_quickly | 4789 |

| phenotype | plot_label | paper_phenotype_category | pheweb_phenotype_category | pheweb_phenotype_name | sample_size |
|---|---|---|---|---|---|
| afus_dora_9_inspects_unfamilar_foods | Inspects unfamiliar foods before eating | Survey question | Eating Behavior | afus_dora_9_inspects_unfamilar_foods | 4789 |
| afus_mdors_10_get_through_tough_times | Helps me get through tough times | Survey question | Dog Owner Relationship | afus_mdors_10_get_through_tough_times | 4789 |
| afus_mdors_11_dog_for_comfort | Is there whenever i need to be comforted | Survey question | Dog Owner Relationship | afus_mdors_11_dog_for_comfort | 4789 |
| afus_mdors_12_close_proximity | I would like to have my dog near me all the time | Survey question | Dog Owner Relationship | afus_mdors_12_close_proximity | 4789 |
| afus_mdors_13_constant_companionship | Provides me with constant companionship | Survey question | Dog Owner Relationship | afus_mdors_13_constant_companionship | 4789 |
| afus_mdors_14_there_for_me | If everyone else left me, my dog would still be there for me | Survey question | Dog Owner Relationship | afus_mdors_14_there_for_me | 4789 |
| afus_mdors_15_reason_to_get_up | Gives me a reason to get up in morning | Survey question | Dog Owner Relationship | afus_mdors_15_reason_to_get_up | 4789 |
| afus_mdors_16_never_apart | I wish my dog and I never had to be apart | Survey question | Dog Owner Relationship | afus_mdors_16_never_apart | 4789 |
| afus_mdors_17_constantly_attentive | Is constantly attentive to me | Survey question | Dog Owner Relationship | afus_mdors_17_constantly_attentive | 4789 |
| afus_mdors_18_share_secrets | How often: tell dog things you don't tell anyone else | Survey question | Dog Owner Relationship | afus_mdors_18_share_secrets | 4789 |
| afus_mdors_19_trauma_of_death | How traumatic when dog dies | Survey question | Dog Owner Relationship | afus_mdors_19_trauma_of_death | 4789 |
| afus_mdors_1_play_games | How often: play games with dog | Survey question | Dog Owner Relationship | afus_mdors_1_play_games | 4789 |
| afus_mdors_20_care_is_chore | How often: feel that looking after dog is a chore | Survey question | Dog Owner Relationship | afus_mdors_20_care_is_chore | 4789 |
| afus_mdors_21_interferes_with_activities | How often: stop you from doing things you want to | Survey question | Dog Owner Relationship | afus_mdors_21_interferes_with_activities | 4789 |
| afus_mdors_23_annoying_to_change_plans | Annoying when I have to change plans because of dog | Survey question | Dog Owner Relationship | afus_mdors_23_annoying_to_change_plans | 4789 |
| afus_mdors_24_stopped_enjoyable_activities | Bothers me that dog stops me from doing things I enjoy | Survey question | Dog Owner Relationship | afus_mdors_24_stopped_enjoyable_activities | 4789 |

| phenotype | plot_label | paper_phenotype_category | pheweb_phenotype_category | pheweb_phenotype_name | sample_size |
|---|---|---|---|---|---|
| afus_mdors_25_dont_like_ownership | There are major aspects of owning a dog I don't like | Survey question | Dog Owner Relationship | afus_mdors_25_dont_like_ownership | 4789 |
| afus_mdors_26_makes_mess | Makes too much mess | Survey question | Dog Owner Relationship | afus_mdors_26_makes_mess | 4789 |
| afus_mdors_27_costs_money | Costs too much money | Survey question | Dog Owner Relationship | afus_mdors_27_costs_money | 4789 |
| afus_mdors_28_hard_to_care_for | How hard is it to look after your dog | Survey question | Dog Owner Relationship | afus_mdors_28_hard_to_care_for | 4789 |
| afus_mdors_2_treats | How often: give dog treats | Survey question | Dog Owner Relationship | afus_mdors_2_treats | 4789 |
| afus_mdors_3_kiss | How often: kiss dog | Survey question | Dog Owner Relationship | afus_mdors_3_kiss | 4789 |
| afus_mdors_4_dog_in_car | How often: take dog in car | Survey question | Dog Owner Relationship | afus_mdors_4_dog_in_car | 4789 |
| afus_mdors_7_grooming_frequency | How often: groom dog | Survey question | Dog Owner Relationship | afus_mdors_7_grooming_frequency | 4789 |
| afus_mdors_8_visiting_other_people | How often: take dog to visit people | Survey question | Dog Owner Relationship | afus_mdors_8_visiting_other_people | 4789 |
| afus_mdors_9_buy_presents | How often: buy dog presents | Survey question | Dog Owner Relationship | afus_mdors_9_buy_presents | 4789 |
| cslb_active_6mo | How much time active vs. 6 months ago | Survey question | Canine Social and Learned Behavior | cslb_active_6mo | 6773 |
| cslb_avoid | How often: walk away while, or avoid, being petted | Survey question | Canine Social and Learned Behavior | cslb_avoid | 6773 |
| cslb_find_food | How often: difficulty finding food dropped on floor | Survey question | Canine Social and Learned Behavior | cslb_find_food | 6773 |
| cslb_pace | How often: pace up and down, circle, wander | Survey question | Canine Social and Learned Behavior | cslb_pace | 6773 |
| db_aggression_level_delivery_workers_at_home | Aggression when delivery workers approach your home | Survey question | Dog Behavior | db_aggression_level_delivery_workers_at_home | 7328 |
| db_aggression_level_familiar_dog_while_eating | Aggression when approached while eating by household dog | Survey question | Dog Behavior | db_aggression_level_familiar_dog_while_eating | 4692 |
| db_aggression_level_familiar_dog_while_playing | Aggression with familiar dog while playing with a toy | Survey question | Dog Behavior | db_aggression_level_familiar_dog_while_playing | 4902 |

| phenotype | plot_label | paper_phenotype_category | pheweb_phenotype_category | pheweb_phenotype_name | sample_size |
|---|---|---|---|---|---|
| db_aggression_level_on_leash_unknown_dog | Aggression when approached by unfamiliar dog on leash | Survey question | Dog Behavior | db_aggression_level_on_leash_unknown_dog | 7310 |
| db_aggression_level_on_leash_unknown_human | Aggression when approached by unfamiliar person on leash | Survey question | Dog Behavior | db_aggression_level_on_leash_unknown_human | 7342 |
| db_aggression_level_unknown_aggressive_dog | Aggression when barked, growled, or lunged at by another dog | Survey question | Dog Behavior | db_aggression_level_unknown_aggressive_dog | 7219 |
| db_aggression_level_unknown_human_near_yard | Aggression when strangers walk past and dog is outside | Survey question | Dog Behavior | db_aggression_level_unknown_human_near_yard | 7313 |
| db_attention_seeking_follows_humans_frequency | Tends to follow you about the house, from room to room | Survey question | Dog Behavior | db_attention_seeking_follows_humans_frequency | 7365 |
| db_attention_seeking_sits_close_to_humans_frequency | Tends to sit close when you are sitting | Survey question | Dog Behavior | db_attention_seeking_sits_close_to_humans_frequency | 7372 |
| db_barks_frequency | Barks persistently when alarmed or excited | Survey question | Dog Behavior | db_barks_frequency | 7370 |
| db_chases_birds_frequency | Chases birds, given the chance | Survey question | Dog Behavior | db_chases_birds_frequency | 7346 |
| db_chases_squirrels_frequency | Chases squirrels, rabbits, etc., given the chance | Survey question | Dog Behavior | db_chases_squirrels_frequency | 7313 |
| db_chases_tail_frequency | Chases own tail | Survey question | Dog Behavior | db_chases_tail_frequency | 7342 |
| db_chews_inappropriate_objects_frequency | Chews inappropriate objects | Survey question | Dog Behavior | db_chews_inappropriate_objects_frequency | 7350 |
| db_energetic_frequency | Active, energetic, always on the go | Survey question | Dog Behavior | db_energetic_frequency | 7325 |
| db_escapes_home_or_property_frequency | Escapes from home or yard | Survey question | Dog Behavior | db_escapes_home_or_property_frequency | 7299 |
| db_excitement_level_before_car_ride | Excitement before being taken on a car trip | Survey question | Dog Behavior | db_excitement_level_before_car_ride | 7358 |
| db_excitement_level_before_walk | Excitement before being taken for a walk | Survey question | Dog Behavior | db_excitement_level_before_walk | 7355 |
| db_fear_level_bathed_at_home | Fear when groomed or bathed | Survey question | Dog Behavior | db_fear_level_bathed_at_home | 7281 |
| db_fear_level_loud_noises | Fear of sudden or loud noises | Survey question | Dog Behavior | db_fear_level_loud_noises | 7352 |

| phenotype | plot_label | paper_phenotype_category | pheweb_phenotype_category | pheweb_phenotype_name | sample_size |
|---|---|---|---|---|---|
| db_fear_level_nails_clipped_at_home | Fear when having nails clipped | Survey question | Dog Behavior | db_fear_level_nails_clipped_at_home | 6993 |
| db_fear_level_unknown_aggressive_dog | Fear when barked, growled, or lunged at by unfamiliar dog | Survey question | Dog Behavior | db_fear_level_unknown_aggressive_dog | 7210 |
| db_fear_level_unknown_dogs | Fear when approached by unfamiliar dog | Survey question | Dog Behavior | db_fear_level_unknown_dogs | 7302 |
| db_fear_level_unknown_human_away_from_home | Fear when approached by unfamiliar person away from home | Survey question | Dog Behavior | db_fear_level_unknown_human_away_from_home | 7324 |
| db_fear_level_unknown_human_touch | Fear when unfamiliar person tries to touch | Survey question | Dog Behavior | db_fear_level_unknown_human_touch | 7334 |
| db_fear_level_unknown_objects_outside | Fear of unfamiliar objects on or near the sidewalk | Survey question | Dog Behavior | db_fear_level_unknown_objects_outside | 7338 |
| db_fear_level_unknown_situations | Fear of unfamiliar situations | Survey question | Dog Behavior | db_fear_level_unknown_situations | 7331 |
| db_hyperactive_frequency | Hyperactive, restless | Survey question | Dog Behavior | db_hyperactive_frequency | 7343 |
| db_left_alone_barking_frequency | Barking or whining when alone | Survey question | Dog Behavior | db_left_alone_barking_frequency | 7355 |
| db_left_alone_restlessness_frequency | Restlessness, agitation or pacing when alone | Survey question | Dog Behavior | db_left_alone_restlessness_frequency | 7310 |
| db_left_alone_scratching_frequency | Chewing or scratching at doors etc when alone | Survey question | Dog Behavior | db_left_alone_scratching_frequency | 7353 |
| db_playful_frequency | Playful, puppyish, boisterous | Survey question | Dog Behavior | db_playful_frequency | 7362 |
| db_pulls_leash_frequency | Pulls hard when on the leash | Survey question | Dog Behavior | db_pulls_leash_frequency | 7318 |
| db_training_distraction_frequency | Easily distracted | Survey question | Dog Behavior | db_training_distraction_frequency | 7370 |
| db_training_obeys_sit_command_frequency | Obeys sit command immediately | Survey question | Dog Behavior | db_training_obeys_sit_command_frequency | 7353 |
| db_training_obeys_stay_command_frequency | Obeys stay command immediately | Survey question | Dog Behavior | db_training_obeys_stay_command_frequency | 7302 |
| dd_weight_lbs | Dog's weight | Survey question | Dog Demographics | dd_weight_lbs | 7378 |

| phenotype | plot_label | paper_phenotype_category | pheweb_phenotype_category | pheweb_phenotype_name | sample_size |
|---|---|---|---|---|---|
| de_air_cleaner_present | Air cleaners present | Survey question | Dog Environment | de_air_cleaner_present | 7335 |
| de_air_freshener_present | Air fresheners present | Survey question | Dog Environment | de_air_freshener_present | 7338 |
| de_daytime_sleep_avg_hours | How many hours sleeping per day | Survey question | Dog Environment | de_daytime_sleep_avg_hours | 6623 |
| de_drinks_outdoor_water | Drinks from outdoor water sources | Survey question | Dog Environment | de_drinks_outdoor_water | 7378 |
| de_drinks_outdoor_water_frequency | How frequently drinks from outdoor water sources | Survey question | Dog Environment | de_drinks_outdoor_water_frequency | 7378 |
| de_eats_feces | Consume feces | Survey question | Dog Environment | de_eats_feces | 7073 |
| de_eats_grass_frequency | How often: eats grass | Survey question | Dog Environment | de_eats_grass_frequency | 7378 |
| de_gas_fireplace_present | Gas fireplace present | Survey question | Dog Environment | de_gas_fireplace_present | 7346 |
| de_hepa_present | HEPA filters in air circulation present | Survey question | Dog Environment | de_hepa_present | 7097 |
| de_incense_present | Incense or scented candles present | Survey question | Dog Environment | de_incense_present | 7354 |
| de_interacts_with_neighborhood_animals | Interacts with other animals in neighborhood | Survey question | Dog Environment | de_interacts_with_neighborhood_animals | 7378 |
| de_interacts_with_neighborhood_humans | Interacts with other humans in neighborhood | Survey question | Dog Environment | de_interacts_with_neighborhood_humans | 7378 |
| de_licks_chews_or_plays_with_non_toys | Regularly licks or chews non-toy items | Survey question | Dog Environment | de_licks_chews_or_plays_with_non_toys | 7378 |
| de_nighttime_sleep_avg_hours | How many hours sleeping per night | Survey question | Dog Environment | de_nighttime_sleep_avg_hours | 7240 |
| de_recent_toxins_or_hazards_ingested_frequency | How often: ingested poisons or hazardous materials | Survey question | Dog Environment | de_recent_toxins_or_hazards_ingested_frequency | 7378 |
| de_wood_fireplace_present | Wood burning fireplace/stove present | Survey question | Dog Environment | de_wood_fireplace_present | 7344 |
| df_appetite | How is dog's appetite | Survey question | Diet | df_appetite | 6965 |
| df_daily_supplements | Dog gets daily supplements | Survey question | Diet | df_daily_supplements | 6965 |
| df_ever_overweight | Ever overweight | Survey question | Diet | df_ever_overweight | 6660 |
| df_feedings_per_day | How many meals per day | Survey question | Diet | df_feedings_per_day | 6965 |

| phenotype | plot_label | paper_phenotype_category | pheweb_phenotype_category | pheweb_phenotype_name | sample_size |
|---|---|---|---|---|---|
| df_secondary_diet_component_used | Diet has second component >25% | Survey question | Diet | df_secondary_diet_component_used | 6965 |
| df_treats_frequency | How often: treats or other items to eat | Survey question | Diet | df_treats_frequency | 6965 |
| hs_chronic_condition_present | Any ongoing medical conditions | Survey question | Health Status | hs_chronic_condition_present | 7378 |
| hs_chronic_condition_recently_changed_or_treated | Change in medication for chronic condition | Survey question | Health Status | hs_chronic_condition_recently_changed_or_treated | 1853 |
| hs_general_health | Dog's health is ... | Survey question | Health Status | hs_general_health | 7378 |
| hs_new_condition_diagnosed_last_month | Were the conditions diagnosed in the past month | Survey question | Health Status | hs_new_condition_diagnosed_last_month | 855 |
| mp_dental_extraction | Teeth extracted | Survey question | Medication and Preventatives | mp_dental_extraction | 1746 |
| mp_dental_procedure_undergone | Any dental procedures | Survey question | Medication and Preventatives | mp_dental_procedure_undergone | 7378 |
| pa_activity_level | Dog's lifestyle | Survey question | Physical Activity | pa_activity_level | 7378 |
| pa_avg_activity_intensity | Intensity level when active | Survey question | Physical Activity | pa_avg_activity_intensity | 7378 |
| pa_avg_daily_active_hours | Hours active per day | Survey question | Physical Activity | pa_avg_daily_active_hours | 7378 |
| pa_cold_weather_daily_hours_outside | Time outdoors in cold weather | Survey question | Physical Activity | pa_cold_weather_daily_hours_outside | 6334 |
| pa_hot_weather_daily_hours_outside | Time outdoors in hot weather | Survey question | Physical Activity | pa_hot_weather_daily_hours_outside | 6877 |
| pa_moderate_weather_daily_hours_outside | Time outdoors in moderate weather | Survey question | Physical Activity | pa_moderate_weather_daily_hours_outside | 7377 |
| pa_off_leash_walk_average_pace_pct | How often off-leash | Survey question | Physical Activity | pa_off_leash_walk_average_pace_pct | 2975 |
| pa_off_leash_walk_avg_hours | Hours of off-leash walking | Survey question | Physical Activity | pa_off_leash_walk_avg_hours | 2975 |
| pa_off_leash_walk_brisk_pace_pct | How often off-leash activities at a brisk pace | Survey question | Physical Activity | pa_off_leash_walk_brisk_pace_pct | 2975 |
| pa_off_leash_walk_frequency | Frequency active off leash | Survey question | Physical Activity | pa_off_leash_walk_frequency | 7378 |
| pa_off_leash_walk_in_enclosed_area | Ever off leash in enclosed area | Survey question | Physical Activity | pa_off_leash_walk_in_enclosed_area | 2975 |
| pa_off_leash_walk_in_open_area | Ever off leash in open area | Survey question | Physical Activity | pa_off_leash_walk_in_open_area | 2975 |
| pa_off_leash_walk_returns_when_called_frequency | How frequently comes back when called while off leash | Survey question | Physical Activity | pa_off_leash_walk_returns_when_called_frequency | 2975 |

| phenotype | plot_label | paper_phenotype_category | pheweb_phenotype_category | pheweb_phenotype_name | sample_size |
|---|---|---|---|---|---|
| pa_off_leash_walk_run_pace_pct | How often off-leash activities at running pace | Survey question | Physical Activity | pa_off_leash_walk_run_pace_pct | 2975 |
| pa_on_leash_walk_average_pace_pct | How often on-leash activities at average pace | Survey question | Physical Activity | pa_on_leash_walk_average_pace_pct | 7110 |
| pa_on_leash_walk_avg_hours | Hours of leash-walking | Survey question | Physical Activity | pa_on_leash_walk_avg_hours | 7110 |
| pa_on_leash_walk_brisk_pace_pct | How often on-leash activities at a brisk pace | Survey question | Physical Activity | pa_on_leash_walk_brisk_pace_pct | 7110 |
| pa_on_leash_walk_frequency | How often active on a lead/leash | Survey question | Physical Activity | pa_on_leash_walk_frequency | 7378 |
| pa_other_aerobic_activity_avg_hours | Hours of other aerobic activity | Survey question | Physical Activity | pa_other_aerobic_activity_avg_hours | 3614 |
| pa_other_aerobic_activity_avg_intensity | Intensity of activity during aerobic activity | Survey question | Physical Activity | pa_other_aerobic_activity_avg_intensity | 3614 |
| pa_other_aerobic_activity_frequency | Frequency of other aerobic activity | Survey question | Physical Activity | pa_other_aerobic_activity_frequency | 7378 |
| pa_physical_games_frequency | How often: play games with physical activity | Survey question | Physical Activity | pa_physical_games_frequency | 7378 |
| pa_swim | Does your dog go swimming | Survey question | Physical Activity | pa_swim | 7378 |
| pa_swim_cold_weather_frequency | How often: swimming in cold weather | Survey question | Physical Activity | pa_swim_cold_weather_frequency | 1920 |
| pa_swim_hot_weather_frequency | How often: swimming in hot weather | Survey question | Physical Activity | pa_swim_hot_weather_frequency | 2005 |
| pa_swim_moderate_weather_frequency | How often: swimming in moderate weather | Survey question | Physical Activity | pa_swim_moderate_weather_frequency | 2325 |

**Table S3.**

Demographic information, fur traits, and normalized cystathionine values for 50 dogs with blood trait measurements and photographs

| dog_id | single_breed | Breed | sex | weight_kg | Fur Curl | Furnishings | Fur length | Cystathionine |
|---|---|---|---|---|---|---|---|---|
| 310 | TRUE | pug | female | 11.51927438 | 1 | 1 | 1 | -0.6026110809 |
| 695 | FALSE | rottweiler/airedale terrier | male | 37.64172336 | 1 | 1 | 2 | 0.2468598868 |
| 766 | TRUE | welsh terrier | male | 14.51247166 | 2 | 2 | 2 | 0.9252532918 |
| 1443 | TRUE | australian shepherd | female | 15.87301587 | 1 | 1 | 3 | -0.3308558253 |
| 1824 | TRUE | great dane | male | 63.49206349 | 1 | 1 | 1 | -0.029214752 |
| 2775 | TRUE | english shepherd | female | 29.02494331 | 1 | 1 | 2 | 0.7819874195 |
| 2895 | TRUE | akita | male | 47.61904762 | 1 | 1 | 2 | 0.1531169823 |
| 3035 | FALSE | dachshund/west highland white terrier | male | 6.802721088 | 2 | 2 | 3 | -0.2063971032 |
| 3586 | TRUE | labrador retriever | male | 34.46712018 | 1 | 1 | 2 | 0.415526004 |
| 4416 | TRUE | golden retriever | male | 36.05442177 | 1 | 1 | 2 | 0.2520251583 |
| 9113 | TRUE | labrador retriever | male | 26.48526077 | 1 | 1 | 2 | 0.2190681428 |
| 10312 | FALSE | unknown/unknown | female | 17.50566893 | 1 | 1 | 2 | -0.6931054257 |
| 12278 | TRUE | labrador retriever/poodle | male | 21.76870748 | 2 | 2 | 3 | 0.3298542319 |
| 16321 | TRUE | boxer | male | 29.47845805 | 1 | 1 | 1 | 0.6116976138 |
| 18521 | TRUE | poodle | female | 23.58276644 | 2 | 2 | 3 | 0.01730166118 |
| 19232 | TRUE | english setter | female | 21.31519274 | 1 | 1 | 3 | 0.4168443773 |
| 20155 | TRUE | bavarian mountain scent hound | male | 26.16780045 | 1 | 1 | 1 | 0.1669572435 |
| 25292 | TRUE | border collie | male | 24.03628118 | 1 | 1 | 2 | -0.1038846842 |
| 25538 | FALSE | catahoula leopard dog/bluetick coonhound | female | 25.3968254 | 1 | 1 | 1 | -0.1708490959 |
| 26492 | TRUE | greyhound | female | 28.11791383 | 1 | 1 | 1 | 0.8778133593 |
| 27486 | TRUE | english setter | male | 19.50113379 | 1 | 1 | 3 | 0.1134528437 |

| dog_id | single_bre ed | Breed | sex | weight_kg | Fur Curl | Furnishing s | Fur length | Cystathion ine |
|---|---|---|---|---|---|---|---|---|
| 28082 | FALSE | labrador retriever/great pyrenees | male | 39.9092970 5 | 1 | 1 | 2 | 0.05154028 727 |
| 29556 | TRUE | golden retriever | femal e | 28.1179138 3 | 1 | 1 | 3 | 0.30417684 86 |
| 31865 | FALSE | west highland white terrier/unknown | male | 6.80272108 8 | 2 | 2 | 3 | 1.08907702 6 |
| 33975 | FALSE | poodle/cocker spaniel | male | 10.8843537 4 | 2 | 2 | 3 | 0.52449942 66 |
| 35367 | FALSE | great pyrenees/border collie | femal e | 39.9092970 5 | 1 | 1 | 3 | 0.00891587 0711 |
| 39567 | FALSE | miniature schnauzer/yorks hire terrier | femal e | 4.08163265 3 | 1 | 2 | 2 | 0.10641384 48 |
| 42183 | FALSE | russell terrier/unknown | femal e | 6.34920634 9 | 1 | 2 | 2 | 0.21375778 21 |
| 42423 | TRUE | rhodesian ridgeback | male | 41.2698412 7 | 1 | 1 | 1 | - 0.55289385 68 |
| 43622 | TRUE | poodle (toy) | femal e | 2.04081632 7 | 2 | 2 | 3 | 0.91295457 45 |
| 44534 | TRUE | labrador retriever | femal e | 33.1065759 6 | 1 | 1 | 2 | 0.00839637 3529 |
| 54174 | TRUE | west highland white terrier | male | 10.7029478 5 | 1 | 2 | 2 | - 0.15476291 19 |
| 54218 | TRUE | australian cattle dog | femal e | 24.9433106 6 | 1 | 1 | 2 | - 0.28290681 85 |
| 54857 | FALSE | border collie/labrador retriever | male | 26.3038548 8 | 1 | 1 | 2 | - 0.18010809 16 |
| 54903 | FALSE | labrador retriever/treeing walker coonhound | male | 43.0839002 3 | 1 | 1 | 2 | - 1.33428596 1 |
| 55103 | TRUE | german shepherd dog | femal e | 47.6190476 2 | 1 | 1 | 2 | - 0.27680651 72 |
| 55370 | TRUE | bichon frise | femal e | 4.98866213 2 | 2 | 2 | 3 | 0.68509849 46 |
| 55566 | FALSE | chihuahua/jack russell terrier | femal e | 5.44217687 1 | 1 | 2 | 2 | 0.29224167 36 |
| 55684 | FALSE | poodle/golden retriever | male | 17.2335600 9 | 1 | 2 | 3 | 0.61728016 59 |
| 57351 | TRUE | poodle | male | 10.8843537 4 | 2 | 2 | 3 | - 0.13206516 12 |

| dog_id | single_breed | Breed | sex | weight_kg | Fur Curl | Furnishings | Fur length | Cystathionine |
|---|---|---|---|---|---|---|---|---|
| 62093 | FALSE | american pitbull terrier/labrador retriever | female | 24.94331066 | 1 | 1 | 1 | -0.2101897279 |
| 73365 | FALSE | chihuahua/rat terrier | female | 9.070294785 | 1 | 1 | 1 | -0.2829545208 |
| 76109 | TRUE | newfoundland | female | 63.49206349 | 1 | 1 | 3 | -0.2813746192 |
| 84841 | FALSE | german shepherd dog/labrador retriever | female | 11.33786848 | 1 | 1 | 2 | -0.9199955986 |
| 87040 | TRUE | labrador retriever | female | 24.48979592 | 1 | 1 | 2 | -0.1060013345 |
| 96234 | TRUE | cocker spaniel | male | 8.390022676 | 1 | 2 | 3 | 1.092841363 |
| 96928 | TRUE | mastiff | female | 18.14058957 | 1 | 1 | 2 | -0.8343893232 |
| 100293 | FALSE | american pitbull terrier/bullmastiff | female | 23.1292517 | 1 | 1 | 1 | -0.8821893615 |
| 109353 | TRUE | australian shepherd | male | 15.87301587 | 1 | 1 | 3 | 0.2396857423 |
| 111720 | TRUE | poodle | male | 28.11791383 | 2 | 2 | 3 | 0.332343815 |

**Table S4.**
Mortality hazard ratios of blood traits from human studies

| phenotype | plot label | PMID | Human hazard ratio | comment | Models: AgingAI | Models: phenoAge |
|---|---|---|---|---|---|---|
| krt_cp_globulins_value_ln_transformed | globulins | 25816467 | 1.92 | 3.7 - < 4 g/dL globulin level | TRUE | FALSE |
| krt_cbc_rbc_sqrt_transformed | red blood cells | 35490162 | 1.66 | low red blood cells | TRUE | FALSE |
| krt_cbc_abs_monocytes_sqrt_transformed | absolute monocytes | 35078679 | 1.53 | | TRUE | FALSE |
| krt_cbc_rel_monocytes_sqrt_transformed | relative monocytes | 35078679 | 1.53 | | TRUE | FALSE |
| krt_cbc_abs_neutrophils_sqrt_transformed | absolute neutrophils | 35078679 | 1.51 | | TRUE | FALSE |
| krt_cbc_rel_neutrophils_sqrt_transformed | relative neutrophils | 35078679 | 1.51 | | TRUE | FALSE |
| krt_cp_bun_value_ln_transformed | blood urea nitrogen | 36678332 | 1.48 | higher BUN compared to lower BUN | TRUE | FALSE |
| krt_cp_alt_value_ln_transformed | alanine aminotransferase | 35435277 | 1.45 | ALT levels of 10-19 IU/L (mid level) | TRUE | FALSE |
| krt_cp_potassium_value_ln_transformed | potassium | 33849496 | 1.43 | high potassium, multivariable adjusted | TRUE | FALSE |
| krt_cbc_hgb_sqrt_transformed | hemoglobin | 29378732 | 1.39 | high hemoglobin | TRUE | FALSE |
| krt_cp_bilirubin_total_value_ln_transformed | total bilirubin | 24728477 | 1.36 | lower bilirubin associated with higher hazard of mortality | TRUE | FALSE |
| krt_cp_magnesium_value_ln_transformed | magnesium | 28890274 | 1.34 | The multivariate-adjusted HRs (95% CIs) of all-cause mortality across increasing categories of Mg were 1.34 (1.02, 1.77), 0.94 (0.75, 1.18), 1.08 (0.97, 1.19), 1.00 (referent), 1.05 (0.95, 1.16), 0.96 (0.79, 1.15), and 0.98 (0.76, 1.26). | TRUE | FALSE |
| krt_cbc_abs_lymphocytes_sqrt_transformed | absolute lymphocytes | 35078679 | 1.2 | | TRUE | FALSE |
| krt_cbc_abs_eosinophils_sqrt_transformed | absolute eosinophils | 23765950 | 1.16 | mild eosinophilia | TRUE | FALSE |
| krt_cbc_rel_eosinophils_sqrt_ | relative | 23765950 | 1.16 | mild eosinophilia | TRUE | FALSE |

| phenotype | plot label | PMID | Human hazard ratio | comment | Models: AgingAI | Models: phenoAge |
|---|---|---|---|---|---|---|
| transformed | eosinophils | | | | | |
| krt_cp_chloride_value_ln_transformed | chloride | 36809243 | 2.41 | low serum chloride levels - hypochloremia (< 97 mmol/L) | TRUE | FALSE |
| krt_cp_cholesterol_value_ln_transformed | cholesterol | 10891962 | 1.23 | relative risk of Chicago Heart Association Detection Project in Industry (CHA) cohort with serum cholesterol level higher by 40 mg/dL (1.03 mmol/L) | TRUE | FALSE |
| krt_cbc_hct_sqrt_transformed | hematocrit | 23569195 | 2.12 | 15–29 (% hematocrit) males and females adjusted HR value | TRUE | FALSE |
| krt_cbc_mchc_sqrt_transformed | mean corpuscular hemoglobin concentration | 39093828 | 0.94 | continuous values, model III that adjusted for a comprehensive set of covariates | TRUE | FALSE |
| krt_cbc_mch_sqrt_transformed | mean corpuscular hemoglobin | 39093828 | 1.02 | continuous values, model III that adjusted for a comprehensive set of covariates | TRUE | FALSE |
| Krt_cbc_plt_sqrt_transformed | platelet | 26864338 | 2.17 | low platelet levels | TRUE | FALSE |
| krt_cp_sodium_value_ln_transformed | sodium | 37407935 | 1.8 | sodium levels < 135 mmol/L from National Kidney Disease Surveillance System in Ireland | TRUE | FALSE |
| krt_cp_total_protein_value_ln_transformed | total protein | 36317096 | 1.75 | low total protein levels of hospital in-patients without malignancies in Japan | TRUE | FALSE |
| krt_cp_albumin_value_ln_transformed | albumin | 29649489 | 1.57 | low albumin (3.8–3.9 g/dL) | TRUE | TRUE |
| krt_cp_alkp_value_ln_transformed | alkaline phosphatase | 19841303 | 1.43 | highest tertile of alkaline phosphatase | TRUE | TRUE |
| krt_cbc_total_wbcs_sqrt_trans | total white | 35078679 | 1.42 | | TRUE | TRUE |

| phenotype | plot label | PMID | Human hazard ratio | comment | Models: AgingAI | Models: phenoAge |
|---|---|---|---|---|---|---|
| formed | blood cells | | | | | |
| krt_cbc_rel_lymphocytes_sqrt_transformed | relative lymphocytes | 35078679 | 1.2 | | TRUE | TRUE |
| Creatinine | Creatinine | 37717037 | 1 | | TRUE | TRUE |
| krt_cp_creatinine_value_ln_transformed | creatinine (clinical) | 37717037 | 1 | | TRUE | TRUE |
| Glucose | Glucose | 31266835 | 1.56 | low fasting glucose | TRUE | TRUE |
| krt_cbc_mcv_sqrt_transformed | mean corpuscular volume | 39093828 | 1.02 | continuous values, model III that adjusted for a comprehensive set of covariates | TRUE | TRUE |
| krt_cbc_rdw_sqrt_transformed | red blood cell width distribution | 24173039 | 3.83 | higher RDW values (RDW >17%) | TRUE | TRUE |

**Table S5.**

Cox proportional hazard modeling with composite ancestry risk for risk-associated and protective blood traits

| term | estimate | std error | statistic | p value | conf low | conf high | phenotype |
|---|---|---|---|---|---|---|---|
| recruitmentAge | 3.7145 | 0.0986 | 13.3075 | 0.0000 | 3.0617 | 4.5065 | krt_cp_alb_glob_ratio_value_ln_transformed |
| Weight_Class_10KGBin_at_HLES10-19.9 | 1.1231 | 0.3300 | 0.3517 | 0.7250 | 0.5882 | 2.1445 | krt_cp_alb_glob_ratio_value_ln_transformed |
| Weight_Class_10KGBin_at_HLES20-29.9 | 1.9966 | 0.3082 | 2.2437 | 0.0249 | 1.0914 | 3.6526 | krt_cp_alb_glob_ratio_value_ln_transformed |
| Weight_Class_10KGBin_at_HLES30-39.9 | 2.0046 | 0.3207 | 2.1686 | 0.0301 | 1.0692 | 3.7583 | krt_cp_alb_glob_ratio_value_ln_transformed |
| Weight_Class_10KGBin_at_HLES40+ | 3.1618 | 0.3451 | 3.3359 | 0.0009 | 1.6077 | 6.2182 | krt_cp_alb_glob_ratio_value_ln_transformed |
| sexMale | 1.2544 | 0.2014 | 1.1254 | 0.2604 | 0.8453 | 1.8616 | krt_cp_alb_glob_ratio_value_ln_transformed |
| krt_cp_alb_glob_ratio_value | 0.7211 | 0.0840 | -3.8925 | 0.0001 | 0.6117 | 0.8502 | krt_cp_alb_glob_ratio_value_ln_transformed |
| alb_glob_risk_breeds_composite_ancestry | 0.9983 | 0.4154 | -0.0041 | 0.9968 | 0.4423 | 2.2534 | krt_cp_alb_glob_ratio_value_ln_transformed |
| recruitmentAge | 3.8066 | 0.0977 | 13.6865 | 0.0000 | 3.1434 | 4.6097 | krt_cp_globulins_value_ln_transformed |
| Weight_Class_10KGBin_at_HLES10-19.9 | 1.1087 | 0.3293 | 0.3134 | 0.7540 | 0.5815 | 2.1140 | krt_cp_globulins_value_ln_transformed |
| Weight_Class_10KGBin_at_HLES20-29.9 | 2.1942 | 0.3133 | 2.5080 | 0.0121 | 1.1873 | 4.0549 | krt_cp_globulins_value_ln_transformed |
| Weight_Class_10KGBin_at_HLES30-39.9 | 2.3354 | 0.3317 | 2.5572 | 0.0106 | 1.2191 | 4.4740 | krt_cp_globulins_value_ln_transformed |
| Weight_Class_10KGBin_at_HLES40+ | 3.8688 | 0.3486 | 3.8806 | 0.0001 | 1.9535 | 7.6621 | krt_cp_globulins_value_ln_transformed |
| sexMale | 1.3047 | 0.2012 | 1.3216 | 0.1863 | 0.8795 | 1.9355 | krt_cp_globulins_value_ln_transformed |
| krt_cp_globulins_value | 1.3216 | 0.0970 | 2.8747 | 0.0040 | 1.0928 | 1.5984 | krt_cp_globulins_value_ln_transformed |
| globulin_risk_breeds_composite_ancestry | 0.6802 | 0.3578 | -1.0772 | 0.2814 | 0.3374 | 1.3714 | krt_cp_globulins_value_ln_transformed |
| recruitmentAge | 3.6031 | 0.1015 | 12.6265 | 0.0000 | 2.9530 | 4.3963 | krt_cp_potassium_value_ln_transformed |
| Weight_Class_10KGBin_at_HLES10-19.9 | 1.1964 | 0.3324 | 0.5395 | 0.5895 | 0.6237 | 2.2952 | krt_cp_potassium_value_ln_transformed |
| Weight_Class_10KGBin_at_HLES20-29.9 | 2.1497 | 0.3070 | 2.4931 | 0.0127 | 1.1778 | 3.9237 | krt_cp_potassium_value_ln_transformed |
| Weight_Class_10KGBin_at_HLES30-39.9 | 2.0680 | 0.3215 | 2.2600 | 0.0238 | 1.1013 | 3.8833 | krt_cp_potassium_value_ln_transformed |
| Weight_Class_10KGBin_at_HLES40+ | 3.9353 | 0.3363 | 4.0737 | 0.0000 | 2.0357 | 7.6073 | krt_cp_potassium_value_ln_transformed |
| sexMale | 1.2896 | 0.1996 | 1.2742 | 0.2026 | 0.8721 | 1.9071 | krt_cp_potassium_value_ln_transformed |
| krt_cp_potassium_value | 1.3041 | 0.0977 | 2.7175 | 0.0066 | 1.0768 | 1.5793 | krt_cp_potassium_value_ln_transformed |
| potassium_risk_breeds_composite_ancestry | 0.3387 | 0.7878 | -1.3743 | 0.1693 | 0.0723 | 1.5862 | krt_cp_potassium_value_ln_transformed |
| recruitmentAge | 3.5818 | 0.1010 | 12.6273 | 0.0000 | 2.9383 | 4.3662 | krt_cp_sp_ratio_value_ln_transformed |

| term | estimate | std error | statistic | p value | conf low | conf high | phenotype |
|---|---|---|---|---|---|---|---|
| Weight_Class_10KGBin_at_HLES10-19.9 | 1.2708 | 0.3363 | 0.7126 | 0.4761 | 0.6574 | 2.4565 | krt_cp_sp_ratio_value_ln_transformed |
| Weight_Class_10KGBin_at_HLES20-29.9 | 2.3263 | 0.3070 | 2.7502 | 0.0060 | 1.2746 | 4.2458 | krt_cp_sp_ratio_value_ln_transformed |
| Weight_Class_10KGBin_at_HLES30-39.9 | 2.2745 | 0.3187 | 2.5786 | 0.0099 | 1.2179 | 4.2478 | krt_cp_sp_ratio_value_ln_transformed |
| Weight_Class_10KGBin_at_HLES40+ | 4.3049 | 0.3328 | 4.3863 | 0.0000 | 2.2422 | 8.2649 | krt_cp_sp_ratio_value_ln_transformed |
| sexMale | 1.3164 | 0.1998 | 1.3761 | 0.1688 | 0.8899 | 1.9472 | krt_cp_sp_ratio_value_ln_transformed |
| krt_cp_sp_ratio_value | 0.7684 | 0.0993 | -2.6522 | 0.0080 | 0.6325 | 0.9336 | krt_cp_sp_ratio_value_ln_transformed |
| sodium_potassium_risk_breeds_composite_ancestry | 0.5009 | 0.7465 | -0.9260 | 0.3544 | 0.1160 | 2.1637 | krt_cp_sp_ratio_value_ln_transformed |
| recruitmentAge | 1.5574 | 0.0321 | 13.8132 | 0.0000 | 1.4625 | 1.6584 | Hydroxyproline |
| best_weight | 1.0168 | 0.0030 | 5.4715 | 0.0000 | 1.0107 | 1.0229 | Hydroxyproline |
| sexMale | 1.6122 | 0.2112 | 2.2614 | 0.0237 | 1.0657 | 2.4389 | Hydroxyproline |
| Hydroxyproline | 0.7415 | 0.1157 | -2.5855 | 0.0097 | 0.5911 | 0.9302 | Hydroxyproline |
| hydroxyproline_risk_breeds_composite_ancestry | 1.2081 | 0.3985 | 0.4743 | 0.6353 | 0.5532 | 2.6384 | Hydroxyproline |

**Table S6.**
Cox proportional hazard modeling with composite ancestry risk for risk-associated and protective blood traits

| phenotype | linked with GH-IGF1 in rodent model | Direction in Ames | Direction in bGH | Direction in GHRKO | Direction in GHRH-KO | refPMCID |
|---|---|---|---|---|---|---|
| Alanine | FALSE | none | none | none | | PMC9959592 |
| Creatine | FALSE | none | | | | PMC4467904 |
| Histidine | FALSE | | none | none | | PMC9959592 |
| Leucine--D-Norleucine | FALSE | none | none | none | | PMC9959592 |
| Phenylalanine | FALSE | | none | none | | PMC9959592 |
| Serine | FALSE | | none | none | | PMC9959592 |
| Trimethylamine-N-Oxide-TMAO | FALSE | none | | | | PMC4467904 |
| Valine | TRUE | | none | none | negative | PMC7066919 |
| 2-Hydroxyisobutyrate-2-Hydroxybutyrate | TRUE | positive | | | | PMC4467904 |
| 3HBA | TRUE | negative | | | | PMC9959592 |
| Arginine | TRUE | | positive | none | | PMC12074916 |
| Asparagine | TRUE | | positive | none | | PMC9959592 |
| Aspartic-Acid | TRUE | | positive | none | | PMC12074916 |
| Choline | TRUE | positive | | | | PMC4467904 |
| Glucose | TRUE | positive | | | | PMC4467904 |
| Glutamine | TRUE | | positive | none | | PMC9959592 |
| Glycerophosphocholine | TRUE | positive | | | | PMC4467904 |
| Glycine | TRUE | none | negative | negative | | PMC9959592 |
| Hydroxyproline | TRUE | | positive | positive | | PMC9959592 |
| iso-Leucine--allo-isoLeucine | TRUE | positive | none | none | | PMC4467904 |
| Lactate | TRUE | positive | | | | PMC4467904 |
| Lysine | TRUE | | positive | none | | PMC9959592 |
| Methionine | TRUE | positive | positive | none | | PMC4467904 |
| Ornithine | TRUE | | positive | none | | PMC9959592 |
| Proline | TRUE | | positive | none | | PMC9959592 |
| Pyruvate | TRUE | positive | | | | PMC4467904 |
| Sarcosine | TRUE | negative | | | | PMC6280974 |
| Taurine | TRUE | | negative | none | | PMC9959592 |
| Threonine | TRUE | | positive | none | | PMC9959592 |
| Tryptophan | TRUE | | positive | negative | | PMC9959592 |
| Tyrosine | TRUE | | positive | none | | PMC9959592 |

**Table S**7.
Pearson's correlation coefficient of phenotypes with genome-wide significant associations at the *ALB* locus

| | **albumin** | **globulins** | **albumin globulin ratio** | **Tryptophan** | **L-Kynurenine** | **Indole-3-Lactate** | **Indole-3-Acetic-Acid** | **Indole-3-Propionate** | **Margaric-Acid** |
|---|---|---|---|---|---|---|---|---|---|
| **albumin** | 1.000 | -0.020 | 0.526 | 0.290 | 0.075 | 0.216 | 0.042 | 0.062 | -0.073 |
| **globulins** | -0.020 | 1.000 | -0.844 | -0.067 | -0.113 | -0.053 | -0.127 | -0.151 | -0.025 |
| **albumin globulin ratio** | 0.526 | -0.844 | 1.000 | 0.201 | 0.132 | 0.158 | 0.129 | 0.153 | -0.023 |
| **Tryptophan** | 0.290 | -0.067 | 0.201 | 1.000 | 0.604 | 0.479 | 0.413 | 0.238 | -0.325 |
| **L-Kynurenine** | 0.075 | -0.113 | 0.132 | 0.604 | 1.000 | 0.338 | 0.429 | 0.238 | -0.265 |
| **Indole-3-Lactate** | 0.216 | -0.053 | 0.158 | 0.479 | 0.338 | 1.000 | 0.353 | 0.091 | -0.264 |
| **Indole-3-Acetic-Acid** | 0.042 | -0.127 | 0.129 | 0.413 | 0.429 | 0.353 | 1.000 | 0.185 | -0.146 |
| **Indole-3-Propionate** | 0.062 | -0.151 | 0.153 | 0.238 | 0.238 | 0.091 | 0.185 | 1.000 | 0.049 |
| **Margaric-Acid** | -0.073 | -0.025 | -0.023 | -0.325 | -0.265 | -0.264 | -0.146 | 0.049 | 1.000 |

**Data S1 (separate file)**
All survey questions in the Dog Aging Project.

**Data S2 (separate file)**
SNP-based heritability results.

**Data S3 (separate file)**
Suggestive or genome-wide significant genomic loci and defined regions from association results.

**Data S4 (separate file)**
Dog genomic regions that overlap with human genomic regions for blood trait associations.

**Data S5 (separate file)**
Analysis of variance (ANOVA) of blood traits in single breed dogs.

**Data S6 (separate file)**
Regional associations of fur traits in RSPO2 and FGF5 regions in 50 dogs with blood trait measurements and photographs as well as genome-wide association results for cystathionine in RSPO2 and FGF5 regions in 924 dogs.

**Data S7 (separate file)**
Allele frequencies of 6 cystathionine-associated SNPs across breeds.

**Data S8 (separate file)**
Fixed effects of breed ancestry score from linear mixed effects regression (LMER) analysis of blood traits in all dogs with blood trait measurements.

**Data S9 (separate file)**
Analysis of variance (ANOVA) and effect estimates of breed weight and breed lifespan on Breed Ancestry Scores of blood traits.

**Data S10 (separate file)**
Cox proportional hazards regression modeling results for all-cause mortality risk based on blood traits in individual dogs.

**Data S11 (separate file)**
Combined Z-score from Breed Ancestry Scores and Cox regression hazards to identify protective and risk-associated blood traits.

**Data S12 (separate file)**
Breed-level phenotypes (fur characteristics, breed-average weight, breed-average height, and breed-based mortality risk).

**Data S13 (separate file)**
Demographic data on all dogs enrolled in the Dog Aging Project.

**Data S14 (separate file)**
Demographic, top genetic ancestry, and phenotypic data availability on sequenced dogs of the Dog Aging Project.

**Data S15 (separate file)**
Breed ancestry results of sequenced dogs.

**Data S16 (separate file)**
Survey questions included in our genetic analysis with our coded response and the original Dog Aging Project coded responses.

9. J. Yang, A. Bakshi, Z. Zhu, G. Hemani, A. A. E. Vinkhuyzen, S. H. Lee, M. R. Robinson, J. R. B. Perry, I. M. Nolte, J. V. van Vliet-Ostaptchouk, H. Snieder, LifeLines Cohort Study, T. Esko, L. Milani, R. Mägi, A. Metspalu, A. Hamsten, P. K. E. Magnusson, N. L. Pedersen, E. Ingelsson, N. Soranzo, M. C. Keller, N. R. Wray, M. E. Goddard, P. M. Visscher, Genetic variance estimation with imputed variants finds negligible missing

heritability for human height and body mass index. *Nat. Genet.* **47**, 1114–1120 (2015).

10. J. Yang, S. H. Lee, M. E. Goddard, P. M. Visscher, GCTA: a tool for genome-wide complex trait analysis. *Am. J. Hum. Genet.* **88**, 76–82 (2011).

11. Z. Chen, M. Boehnke, X. Wen, B. Mukherjee, Revisiting the genome-wide significance threshold for common variant GWAS. *G3* **11** (2021).

12. K. Lindblad-Toh, C. M. Wade, T. S. Mikkelsen, E. K. Karlsson, D. B. Jaffe, M. Kamal, M. Clamp, J. L. Chang, E. J. Kulbokas 3rd, M. C. Zody, E. Mauceli, X. Xie, M. Breen, R. K. Wayne, E. A. Ostrander, C. P. Ponting, F. Galibert, D. R. Smith, P. J. DeJong, E. Kirkness, P. Alvarez, T. Biagi, W. Brockman, J. Butler, C.-W. Chin, A. Cook, J. Cuff, M. J. Daly, D. DeCaprio, S. Gnerre, M. Grabherr, M. Kellis, M. Kleber, C. Bardeleben, L. Goodstadt, A. Heger, C. Hitte, L. Kim, K.-P. Koepfli, H. G. Parker, J. P. Pollinger, S. M. J. Searle, N. B. Sutter, R. Thomas, C. Webber, J. Baldwin, A. Abebe, A. Abouelleil, L. Aftuck, M. Ait-Zahra, T. Aldredge, N. Allen, P. An, S. Anderson, C. Antoine, H. Arachchi, A. Aslam, L. Ayotte, P. Bachantsang, A. Barry, T. Bayul, M. Benamara, A. Berlin, D. Bessette, B. Blitshteyn, T. Bloom, J. Blye, L. Boguslavskiy, C. Bonnet, B. Boukhgalter, A. Brown, P. Cahill, N. Calixte, J. Camarata, Y. Cheshatsang, J. Chu, M. Citroen, A. Collymore, P. Cooke, T. Dawoe, R. Daza, K. Decktor, S. DeGray, N. Dhargay, K. Dooley, K. Dooley, P. Dorje, K. Dorjee, L. Dorris, N. Duffey, A. Dupes, O. Egbiremolen, R. Elong, J. Falk, A. Farina, S. Faro, D. Ferguson, P. Ferreira, S. Fisher, M. FitzGerald, K. Foley, C. Foley, A. Franke, D. Friedrich, D. Gage, M. Garber, G. Gearin, G. Giannoukos, T. Goode, A. Goyette, J. Graham, E. Grandbois, K. Gyaltsen, N. Hafez, D. Hagopian, B. Hagos, J. Hall, C. Healy, R. Hegarty, T. Honan, A. Horn, N. Houde, L. Hughes, L. Hunnicutt, M. Husby, B. Jester, C. Jones, A. Kamat, B. Kanga, C. Kells, D. Khazanovich, A. C. Kieu, P. Kisner, M. Kumar, K. Lance, T. Landers, M. Lara, W. Lee, J.-P. Leger, N. Lennon, L. Leuper, S. LeVine, J. Liu, X. Liu, Y. Lokyitsang, T. Lokyitsang, A. Lui, J. Macdonald, J. Major, R. Marabella, K. Maru, C. Matthews, S. McDonough, T. Mehta, J. Meldrim, A. Melnikov, L. Meneus, A. Mihalev, T. Mihova, K. Miller, R. Mittelman, V. Mlenga, L. Mulrain, G. Munson, A. Navidi, J. Naylor, T. Nguyen, N. Nguyen, C. Nguyen, T. Nguyen, R. Nicol, N. Norbu, C. Norbu, N. Novod, T. Nyima, P. Olandt, B. O'Neill, K. O'Neill, S. Osman, L. Oyono, C. Patti, D. Perrin, P. Phunkhang, F. Pierre, M. Priest, A. Rachupka, S. Raghuraman, R. Rameau, V. Ray, C. Raymond, F. Rege, C. Rise, J. Rogers, P. Rogov, J. Sahalie, S. Settipalli, T. Sharpe, T. Shea, M. Sheehan, N. Sherpa, J. Shi, D. Shih, J. Sloan, C. Smith, T. Sparrow, J. Stalker, N. Stange-Thomann, S. Stavropoulos, C. Stone, S. Stone, S. Sykes, P. Tchuinga, P. Tenzing, S. Tesfaye, D. Thoulutsang, Y. Thoulutsang, K. Topham, I. Topping, T. Tsamla, H. Vassiliev, V. Venkataraman, A. Vo, T. Wangchuk, T. Wangdi, M. Weiand, J. Wilkinson, A. Wilson, S. Yadav, S. Yang, X. Yang, G. Young, Q. Yu, J. Zainoun, L. Zembek, A. Zimmer, E. S. Lander, Genome sequence, comparative analysis and haplotype structure of the domestic dog. *Nature* **438**, 803–819 (2005).

13. T. G. Richardson, G. M. Leyden, Q. Wang, J. A. Bell, B. Elsworth, G. Davey Smith, M. V. Holmes, Characterising metabolomic signatures of lipid-modifying therapies through drug target mendelian randomisation. *PLoS Biol.* **20**, e3001547 (2022).

14. M. K. Karjalainen, S. Karthikeyan, C. Oliver-Williams, E. Sliz, E. Allara, W. T. Fung, P.

Surendran, W. Zhang, P. Jousilahti, K. Kristiansson, V. Salomaa, M. Goodwin, D. A. Hughes, M. Boehnke, L. Fernandes Silva, X. Yin, A. Mahajan, M. J. Neville, N. R. van Zuydam, R. de Mutsert, R. Li-Gao, D. O. Mook-Kanamori, A. Demirkan, J. Liu, R. Noordam, S. Trompet, Z. Chen, C. Kartsonaki, L. Li, K. Lin, F. A. Hagenbeek, J. J. Hottenga, R. Pool, M. A. Ikram, J. van Meurs, T. Haller, Y. Milaneschi, M. Kähönen, P. P. Mishra, P. K. Joshi, E. Macdonald-Dunlop, M. Mangino, J. Zierer, I. E. Acar, C. B. Hoyng, Y. T. E. Lechanteur, L. Franke, A. Kurilshikov, A. Zhernakova, M. Beekman, E. B. van den Akker, I. Kolcic, O. Polasek, I. Rudan, C. Gieger, M. Waldenberger, F. W. Asselbergs, China Kadoorie Biobank Collaborative Group, Estonian Biobank Research Team, FinnGen, C. Hayward, J. Fu, A. I. den Hollander, C. Menni, T. D. Spector, J. F. Wilson, T. Lehtimäki, O. T. Raitakari, B. W. J. H. Penninx, T. Esko, R. G. Walters, J. W. Jukema, N. Sattar, M. Ghanbari, K. Willems van Dijk, F. Karpe, M. I. McCarthy, M. Laakso, M.-R. Järvelin, N. J. Timpson, M. Perola, J. S. Kooner, J. C. Chambers, C. van Duijn, P. E. Slagboom, D. I. Boomsma, J. Danesh, M. Ala-Korpela, A. S. Butterworth, J. Kettunen, Genome-wide characterization of circulating metabolic biomarkers. *Nature* **628**, 130–138 (2024).

15. E. L. Harshfield, E. B. Fauman, D. Stacey, D. S. Paul, D. Ziemek, R. M. Y. Ong, J. Danesh, A. S. Butterworth, A. Rasheed, T. Sattar, Zameer-Ul-Asar, I. Saleem, Z. Hina, U. Ishtiaq, N. Qamar, N. H. Mallick, Z. Yaqub, T. Saghir, S. N. H. Rizvi, A. Memon, M. Ishaq, S. Z. Rasheed, F.-U.-R. Memon, A. Jalal, S. Abbas, P. Frossard, D. Saleheen, A. M. Wood, J. L. Griffin, A. Koulman, Genome-wide analysis of blood lipid metabolites in over 5000 South Asians reveals biological insights at cardiometabolic disease loci. *BMC Med* **19**, 232 (2021).

16. N. Sinnott-Armstrong, Y. Tanigawa, D. Amar, N. Mars, C. Benner, M. Aguirre, G. R. Venkataraman, M. Wainberg, H. M. Ollila, T. Kiiskinen, A. S. Havulinna, J. P. Pirruccello, J. Qian, A. Shcherbina, FinnGen, F. Rodriguez, T. L. Assimes, V. Agarwala, R. Tibshirani, T. Hastie, S. Ripatti, J. K. Pritchard, M. J. Daly, M. A. Rivas, Genetics of 35 blood and urine biomarkers in the UK Biobank. *Nat Genet* **53**, 185–194 (2021).

17. D. Vuckovic, E. L. Bao, P. Akbari, C. A. Lareau, A. Mousas, T. Jiang, M.-H. Chen, L. M. Raffield, M. Tardaguila, J. E. Huffman, S. C. Ritchie, K. Megy, H. Ponstingl, C. J. Penkett, P. K. Albers, E. M. Wigdor, S. Sakaue, A. Moscati, R. Manansala, K. S. Lo, H. Qian, M. Akiyama, T. M. Bartz, Y. Ben-Shlomo, A. Beswick, J. Bork-Jensen, E. P. Bottinger, J. A. Brody, F. J. A. van Rooij, K. N. Chitrala, P. W. F. Wilson, H. Choquet, J. Danesh, E. Di Angelantonio, N. Dimou, J. Ding, P. Elliott, T. Esko, M. K. Evans, S. B. Felix, J. S. Floyd, L. Broer, N. Grarup, M. H. Guo, Q. Guo, A. Greinacher, J. Haessler, T. Hansen, J. M. M. Howson, W. Huang, E. Jorgenson, T. Kacprowski, M. Kähönen, Y. Kamatani, M. Kanai, S. Karthikeyan, F. Koskeridis, L. A. Lange, T. Lehtimäki, A. Linneberg, Y. Liu, L.-P. Lyytikäinen, A. Manichaikul, K. Matsuda, K. L. Mohlke, N. Mononen, Y. Murakami, G. N. Nadkarni, K. Nikus, N. Pankratz, O. Pedersen, M. Preuss, B. M. Psaty, O. T. Raitakari, S. S. Rich, B. A. T. Rodriguez, J. D. Rosen, J. I. Rotter, P. Schubert, C. N. Spracklen, P. Surendran, H. Tang, J.-C. Tardif, M. Ghanbari, U. Völker, H. Völzke, N. A. Watkins, S. Weiss, VA Million Veteran Program, N. Cai, K. Kundu, S. B. Watt, K. Walter, A. B. Zonderman, K. Cho, Y. Li, R. J. F. Loos, J. C. Knight, M. Georges, O. Stegle, E. Evangelou, Y. Okada, D. J. Roberts, M. Inouye, A. D. Johnson, P. L. Auer, W. J. Astle, A. P. Reiner, A. S. Butterworth, W. H. Ouwehand, G. Lettre, V. G. Sankaran, N. Soranzo, The

Polygenic and Monogenic Basis of Blood Traits and Diseases. *Cell* **182**, 1214–1231.e11 (2020).

18. A. Verma, J. E. Huffman, A. Rodriguez, M. Conery, M. Liu, Y.-L. Ho, Y. Kim, D. A. Heise, L. Guare, V. A. Panickan, H. Garcon, F. Linares, L. Costa, I. Goethert, R. Tipton, J. Honerlaw, L. Davies, S. Whitbourne, J. Cohen, D. C. Posner, R. Sangar, M. Murray, X. Wang, D. R. Dochtermann, P. Devineni, Y. Shi, T. N. Nandi, T. L. Assimes, C. A. Brunette, R. J. Carroll, R. Clifford, S. Duvall, J. Gelernter, A. Hung, S. K. Iyengar, J. Joseph, R. Kember, H. Kranzler, C. M. Kripke, D. Levey, S.-W. Luoh, V. C. Merritt, C. Overstreet, J. D. Deak, S. F. A. Grant, R. Polimanti, P. Roussos, G. Shakt, Y. V. Sun, N. Tsao, S. Venkatesh, G. Voloudakis, A. Justice, E. Begoli, R. Ramoni, G. Tourassi, S. Pyarajan, P. Tsao, C. J. O'Donnell, S. Muralidhar, J. Moser, J. P. Casas, A. G. Bick, W. Zhou, T. Cai, B. F. Voight, K. Cho, J. M. Gaziano, R. K. Madduri, S. Damrauer, K. P. Liao, Diversity and scale: Genetic architecture of 2068 traits in the VA Million Veteran Program. *Science* **385**, eadj1182 (2024).

19. S. Sakaue, M. Kanai, Y. Tanigawa, J. Karjalainen, M. Kurki, S. Koshiba, A. Narita, T. Konuma, K. Yamamoto, M. Akiyama, K. Ishigaki, A. Suzuki, K. Suzuki, W. Obara, K. Yamaji, K. Takahashi, S. Asai, Y. Takahashi, T. Suzuki, N. Shinozaki, H. Yamaguchi, S. Minami, S. Murayama, K. Yoshimori, S. Nagayama, D. Obata, M. Higashiyama, A. Masumoto, Y. Koretsune, FinnGen, K. Ito, C. Terao, T. Yamauchi, I. Komuro, T. Kadowaki, G. Tamiya, M. Yamamoto, Y. Nakamura, M. Kubo, Y. Murakami, K. Yamamoto, Y. Kamatani, A. Palotie, M. A. Rivas, M. J. Daly, K. Matsuda, Y. Okada, A cross-population atlas of genetic associations for 220 human phenotypes. *Nat Genet* **53**, 1415–1424 (2021).

20. J. P. Davis, J. R. Huyghe, A. E. Locke, A. U. Jackson, X. Sim, H. M. Stringham, T. M. Teslovich, R. P. Welch, C. Fuchsberger, N. Narisu, P. S. Chines, A. J. Kangas, P. Soininen, M. Ala-Korpela, J. Kuusisto, F. S. Collins, M. Laakso, M. Boehnke, K. L. Mohlke, Common, low-frequency, and rare genetic variants associated with lipoprotein subclasses and triglyceride measures in Finnish men from the METSIM study. *PLoS Genet* **13**, e1007079 (2017).

21. M.-H. Chen, L. M. Raffield, A. Mousas, S. Sakaue, J. E. Huffman, A. Moscati, B. Trivedi, T. Jiang, P. Akbari, D. Vuckovic, E. L. Bao, X. Zhong, R. Manansala, V. Laplante, M. Chen, K. S. Lo, H. Qian, C. A. Lareau, M. Beaudoin, K. A. Hunt, M. Akiyama, T. M. Bartz, Y. Ben-Shlomo, A. Beswick, J. Bork-Jensen, E. P. Bottinger, J. A. Brody, F. J. A. van Rooij, K. Chitrala, K. Cho, H. Choquet, A. Correa, J. Danesh, E. Di Angelantonio, N. Dimou, J. Ding, P. Elliott, T. Esko, M. K. Evans, J. S. Floyd, L. Broer, N. Grarup, M. H. Guo, A. Greinacher, J. Haessler, T. Hansen, J. M. M. Howson, Q. Q. Huang, W. Huang, E. Jorgenson, T. Kacprowski, M. Kähönen, Y. Kamatani, M. Kanai, S. Karthikeyan, F. Koskeridis, L. A. Lange, T. Lehtimäki, M. M. Lerch, A. Linneberg, Y. Liu, L.-P. Lyytikäinen, A. Manichaikul, H. C. Martin, K. Matsuda, K. L. Mohlke, N. Mononen, Y. Murakami, G. N. Nadkarni, M. Nauck, K. Nikus, W. H. Ouwehand, N. Pankratz, O. Pedersen, M. Preuss, B. M. Psaty, O. T. Raitakari, D. J. Roberts, S. S. Rich, B. A. T. Rodriguez, J. D. Rosen, J. I. Rotter, P. Schubert, C. N. Spracklen, P. Surendran, H. Tang, J.-C. Tardif, R. C. Trembath, M. Ghanbari, U. Völker, H. Völzke, N. A. Watkins, A. B.

Zonderman, VA Million Veteran Program, P. W. F. Wilson, Y. Li, A. S. Butterworth, J.-F. Gauchat, C. W. K. Chiang, B. Li, R. J. F. Loos, W. J. Astle, E. Evangelou, D. A. van Heel, V. G. Sankaran, Y. Okada, N. Soranzo, A. D. Johnson, A. P. Reiner, P. L. Auer, G. Lettre, Trans-ethnic and Ancestry-Specific Blood-Cell Genetics in 746,667 Individuals from 5 Global Populations. *Cell* **182**, 1198–1213.e14 (2020).

22. P. Surendran, I. D. Stewart, V. P. W. Au Yeung, M. Pietzner, J. Raffler, M. A. Wörheide, C. Li, R. F. Smith, L. B. L. Wittemans, L. Bomba, C. Menni, J. Zierer, N. Rossi, P. A. Sheridan, N. A. Watkins, M. Mangino, P. G. Hysi, E. Di Angelantonio, M. Falchi, T. D. Spector, N. Soranzo, G. A. Michelotti, W. Arlt, L. A. Lotta, S. Denaxas, H. Hemingway, E. R. Gamazon, J. M. M. Howson, A. M. Wood, J. Danesh, N. J. Wareham, G. Kastenmüller, E. B. Fauman, K. Suhre, A. S. Butterworth, C. Langenberg, Rare and common genetic determinants of metabolic individuality and their effects on human health. *Nat. Med.* **28**, 2321–2332 (2022).

23. G. Cadby, C. Giles, P. E. Melton, K. Huynh, N. A. Mellett, T. Duong, A. Nguyen, M. Cinel, A. Smith, G. Olshansky, T. Wang, M. Brozynska, M. Inouye, N. S. McCarthy, A. Ariff, J. Hung, J. Hui, J. Beilby, M.-P. Dubé, G. F. Watts, S. Shah, N. R. Wray, W. L. F. Lim, P. Chatterjee, I. Martins, S. M. Laws, T. Porter, M. Vacher, A. I. Bush, C. C. Rowe, V. L. Villemagne, D. Ames, C. L. Masters, K. Taddei, M. Arnold, G. Kastenmüller, K. Nho, A. J. Saykin, X. Han, R. Kaddurah-Daouk, R. N. Martins, J. Blangero, P. J. Meikle, E. K. Moses, Comprehensive genetic analysis of the human lipidome identifies loci associated with lipid homeostasis with links to coronary artery disease. *Nat Commun* **13**, 3124 (2022).

24. M. Pietzner, E. Wheeler, J. Carrasco-Zanini, A. Cortes, M. Koprulu, M. A. Wörheide, E. Oerton, J. Cook, I. D. Stewart, N. D. Kerrison, J. 'an Luan, J. Raffler, M. Arnold, W. Arlt, S. O'Rahilly, G. Kastenmüller, E. R. Gamazon, A. D. Hingorani, R. A. Scott, N. J. Wareham, C. Langenberg, Mapping the proteo-genomic convergence of human diseases. *Science* **374**, eabj1541 (2021).

25. X. Yin, L. S. Chan, D. Bose, A. U. Jackson, P. VandeHaar, A. E. Locke, C. Fuchsberger, H. M. Stringham, R. Welch, K. Yu, L. Fernandes Silva, S. K. Service, D. Zhang, E. C. Hector, E. Young, L. Ganel, I. Das, H. Abel, M. R. Erdos, L. L. Bonnycastle, J. Kuusisto, N. O. Stitziel, I. M. Hall, G. R. Wagner, FinnGen, J. Kang, J. Morrison, C. F. Burant, F. S. Collins, S. Ripatti, A. Palotie, N. B. Freimer, K. L. Mohlke, L. J. Scott, X. Wen, E. B. Fauman, M. Laakso, M. Boehnke, Genome-wide association studies of metabolites in Finnish men identify disease-relevant loci. *Nat Commun* **13**, 1644 (2022).

26. Y. Chen, T. Lu, U. Pettersson-Kymmer, I. D. Stewart, G. Butler-Laporte, T. Nakanishi, A. Cerani, K. Y. H. Liang, S. Yoshiji, J. D. S. Willett, C.-Y. Su, P. Raina, C. M. T. Greenwood, Y. Farjoun, V. Forgetta, C. Langenberg, S. Zhou, C. Ohlsson, J. B. Richards, Genomic atlas of the plasma metabolome prioritizes metabolites implicated in human diseases. *Nat. Genet.* **55**, 44–53 (2023).

27. E. V. Feofanova, M. R. Brown, T. Alkis, A. M. Manuel, X. Li, U. A. Tahir, Z. Li, K. M. Mendez, R. S. Kelly, Q. Qi, H. Chen, M. G. Larson, R. N. Lemaitre, A. C. Morrison, C. Grieser, K. E. Wong, R. E. Gerszten, Z. Zhao, J. Lasky-Su, NHLBI Trans-Omics for

Precision Medicine (TOPMed), B. Yu, Whole-Genome Sequencing Analysis of Human Metabolome in Multi-Ethnic Populations. *Nat Commun* **14**, 3111 (2023).

28. E. V. Feofanova, H. Chen, Y. Dai, P. Jia, M. L. Grove, A. C. Morrison, Q. Qi, M. Daviglus, J. Cai, K. E. North, C. C. Laurie, R. C. Kaplan, E. Boerwinkle, B. Yu, A Genome-wide Association Study Discovers 46 Loci of the Human Metabolome in the Hispanic Community Health Study/Study of Latinos. *Am J Hum Genet* **107**, 849–863 (2020).

29. W. J. Astle, H. Elding, T. Jiang, D. Allen, D. Ruklisa, A. L. Mann, D. Mead, H. Bouman, F. Riveros-Mckay, M. A. Kostadima, J. J. Lambourne, S. Sivapalaratnam, K. Downes, K. Kundu, L. Bomba, K. Berentsen, J. R. Bradley, L. C. Daugherty, O. Delaneau, K. Freson, S. F. Garner, L. Grassi, J. Guerrero, M. Haimel, E. M. Janssen-Megens, A. Kaan, M. Kamat, B. Kim, A. Mandoli, J. Marchini, J. H. A. Martens, S. Meacham, K. Megy, J. O'Connell, R. Petersen, N. Sharifi, S. M. Sheard, J. R. Staley, S. Tuna, M. van der Ent, K. Walter, S.-Y. Wang, E. Wheeler, S. P. Wilder, V. Iotchkova, C. Moore, J. Sambrook, H. G. Stunnenberg, E. Di Angelantonio, S. Kaptoge, T. W. Kuijpers, E. Carrillo-de-Santa-Pau, D. Juan, D. Rico, A. Valencia, L. Chen, B. Ge, L. Vasquez, T. Kwan, D. Garrido-Martín, S. Watt, Y. Yang, R. Guigo, S. Beck, D. S. Paul, T. Pastinen, D. Bujold, G. Bourque, M. Frontini, J. Danesh, D. J. Roberts, W. H. Ouwehand, A. S. Butterworth, N. Soranzo, The Allelic landscape of human blood cell trait variation and links to common complex disease. *Cell* **167**, 1415–1429.e19 (2016).

30. P. G. Hysi, M. Mangino, P. Christofidou, M. Falchi, E. D. Karoly, Nihr Bioresource Investigators, R. P. Mohney, A. M. Valdes, T. D. Spector, C. Menni, Metabolome Genome-Wide Association Study Identifies 74 Novel Genomic Regions Influencing Plasma Metabolites Levels. *Metabolites* **12** (2022).

31. G. Kichaev, G. Bhatia, P.-R. Loh, S. Gazal, K. Burch, M. K. Freund, A. Schoech, B. Pasaniuc, A. L. Price, Leveraging polygenic functional enrichment to improve GWAS power. *Am. J. Hum. Genet.* **104**, 65–75 (2019).

32. U. A. Tahir, D. H. Katz, J. Avila-Pachecho, A. G. Bick, A. Pampana, J. M. Robbins, Z. Yu, Z.-Z. Chen, M. D. Benson, D. E. Cruz, D. Ngo, S. Deng, X. Shi, S. Zheng, A. S. Eisman, L. Farrell, M. E. Hall, A. Correa, R. P. Tracy, P. Durda, K. D. Taylor, Y. Liu, W. C. Johnson, X. Guo, J. Yao, Y.-D. I. Chen, A. W. Manichaikul, F. L. Ruberg, W. S. Blaner, D. Jain, NHLBI Trans-Omics for Precision Medicine 1 Consortium, C. Bouchard, M. A. Sarzynski, S. S. Rich, J. I. Rotter, T. J. Wang, J. G. Wilson, C. B. Clish, P. Natarajan, R. E. Gerszten, Whole Genome Association Study of the Plasma Metabolome Identifies Metabolites Linked to Cardiometabolic Disease in Black Individuals. *Nat Commun* **13**, 4923 (2022).

33. G. Thareja, A. Belkadi, M. Arnold, O. M. E. Albagha, J. Graumann, F. Schmidt, H. Grallert, A. Peters, C. Gieger, T. Q. G. P. R. Consortium, K. Suhre, Differences and commonalities in the genetic architecture of protein quantitative trait loci in European and Arab populations. *Hum Mol Genet* **32**, 907–916 (2023).

34. S. Jeon, H. Choi, Y. Jeon, W.-H. Choi, H. Choi, K. An, H. Ryu, J. Bhak, H. Lee, Y. Kwon, S. Ha, Y. J. Kim, A. Blazyte, C. Kim, Y. Kim, Y. Kang, Y. J. Woo, C. Lee, J. Seo, C. Yoon,

D. Bolser, O. Biro, E.-S. Shin, B. C. Kim, S.-Y. Kim, J.-H. Park, J. Jeon, D. Jung, S. Lee, J. Bhak, Korea4K: whole genome sequences of 4,157 Koreans with 107 phenotypes derived from extensive health check-ups. *Gigascience* **13** (2024).

44. S. Al-Samerria, H. Xu, M. E. Diaz-Rubio, J. Phelan, C. Su, K. Ma, A. Newen, K. Li, S. Yamada, A. L. Negron, F. Wondisford, S. Radovick, Biomarkers of GH deficiency identified in untreated and GH-treated Pit-1 mutant mice. *Front. Endocrinol. (Lausanne)*

**16**, 1539797 (2025).
